# The subcommissural organ regulates brain development via secreted peptides

**DOI:** 10.1101/2024.03.30.587415

**Authors:** Tingting Zhang, Daosheng Ai, Pingli Wei, Ying Xu, Zhanying Bi, Fengfei Ma, Fengzhi Li, Xing-jun Chen, Zhaohuan Zhang, Xiaoxiao Zou, Zongpei Guo, Yue Zhao, Jun-Liszt Li, Meng Ye, Ziyan Feng, Xinshuang Zhang, Lijun Zheng, Jie Yu, Chunli Li, Tianqi Tu, Hongkui Zeng, Jianfeng Lei, Hongqi Zhang, Tao Hong, Li Zhang, Benyan Luo, Zhen Li, Chao Xing, Chenxi Jia, Lingjun Li, Wenzhi Sun, Woo-ping Ge

**Author notes:** To whom the correspondence should be addressed: Woo-ping Ge, Ph.D. (Lead contact), Wenzhi Sun, Ph.D., Lingjun Li, Ph.D.

## Abstract

The subcommissural organ (SCO) is a gland located at the entrance of the aqueduct of Sylvius in the brain. It exists in species as distantly related as amphioxus and humans, but its function is largely unknown. To explore its function, we compared transcriptomes of SCO and non-SCO brain regions and found three genes, *Sspo*, *Car3*, and *Spdef*, that are highly expressed in the SCO. Mouse strains expressing Cre recombinase from endogenous promoter/enhancer elements of these genes were used to genetically ablate SCO cells during embryonic development, resulting in severe hydrocephalus and defects in neuronal migration and development of neuronal axons and dendrites. Unbiased peptidomic analysis revealed enrichment of three SCO-derived peptides, namely thymosin beta 4, thymosin beta 10, and NP24, and their reintroduction into SCO-ablated brain ventricles substantially rescued developmental defects. Together, these data identify a critical role for the SCO in brain development.

---

The SCO is an evolutionarily conserved ependymal gland located under the posterior commissure of the brain at the entrance to the cerebral aqueduct above the third ventricle^1–3^. To date, no specific transgenic mouse lines are available for gene manipulation in SCO cells. As such, most of our knowledge about SCO function has been gleaned from analyses of SCO lesions, animal models, or observations of patients with naturally occurring defects in the brain regions containing the SCO^4,5^. For example, H-Tx model rats develop congenital hydrocephalus^6^. Early studies indicated that such phenotypes are associated with abnormal SCO development owing to failure of the closure of the cerebral aqueduct^7^. Mutant hyh (hydrocephalus with hop gait) mice develop postnatal hydrocephalus with absence of the central canal in the spinal cord and stenosis of the cerebral aqueduct^8^. Further studies with hyh mice revealed impairment of both the SCO and formation of the Reissner Fiber (RF)^9,10^. The RF is an insoluble filament mainly formed by SCO-secreted glycoproteins called SCO-spondins^10^. However, there has been no agreement on SCO or RF function. The lack of the RF in hyh mice may cause distortion and collapse of the ependyma^11^. In zebrafish, the RF participates in spine morphogenesis^12^, and its dysfunction leads to scoliosis phenotypes^13^. The RF in zebrafish was observed to contribute to the mechanosensitivity of neurons that contact the cerebrospinal fluid (CSF)^14^.

In addition to the correlation between SCO malformation and congenital hydrocephalus and scoliosis, it has been proposed to be involved in sodium excretion, diuresis, osmoregulation, and water intake in the brain^9,10,15^, although these findings remain controversial^10,16^. During early brain development in vertebrates, the SCO and its secreted proteins help facilitate axonal guidance in the neural tube midline^17,18^, the regeneration of the neural tube after injury^17^, and the modulation of neuronal aggregation^19^. In addition, soluble factors secreted by the SCO may contribute to neurogenesis in adult mammals^3^. However, most of the information concerning SCO function has come from cell-culture experiments or SCO xenografting and isografting models^20,21^.

The conventional approach to studying the function of a particular brain region is to surgically lesion the area, e.g., by electrolysis^22^. However, the SCO is a tiny gland in the central brain located at the entrance of the aqueduct of Sylvius^10^, making it exceptionally challenging to create lesions without affecting cells in neighboring brain regions. Fortunately, however, genetic ablation via Cre recombinase–mediated expression of Diphtheria toxin fragment A (DTA) in targeted cells represents an efficient and specific way to evaluate the function of a group of cells in the body with minimal impact on adjacent cells^23^. Using this approach to compare the transcriptome of mouse SCO with that of non-SCO brain regions, we were able to identify three genes (*Spdef*, *Car3*, *Sspo*) that are highly expressed in the SCO. Using the promoters for these genes, we developed three Cre mouse strains (*Spdef-Cre*, *Car3-Cre*, *Sspo-Cre*) and one CreER strain (*Sspo-CreER*) for studies of SCO development and function.

## Results

### Generation of SCO-specific Cre mouse lines

We carried out a transcriptomic analysis of the SCO (and hippocampus, for comparison; Fig. 1a) to identify genes that are expressed specifically in the SCO; this analysis yielded >200 genes that were highly expressed specifically in the SCO (Fig. 1b, c). The relative levels of the mRNAs expressed at the whole-brain level were analyzed using an *in situ* hybridization (ISH) dataset from the Allen Mouse Brain Atlas, aiming to narrow the number of candidate genes that showed specific SCO labeling. We identified three genes, namely *Sspo*, *Car3*, and *Spdef*, that were expressed in the SCO and that met our four criteria: 1) mRNA abundance was among the top 10% of all mRNAs expressed in the SCO; 2) the *p*-value was among the top 200 when compared with hippocampal mRNAs; 3) the fold change (upregulated) relative to hippocampal mRNAs was among the top 100; 4) the ISH results from our RNAscope multicolor fluorescence *in situ* hybridization (mFISH) analysis were verified by comparison with ISH data from the Allen Institute ISH database. Strong signals for the corresponding probes of these three candidate genes were limited to the SCO region, implicating their specificity for SCO cells (Fig. 1d–f, Supplementary Fig. 1).

**Fig. 1.**
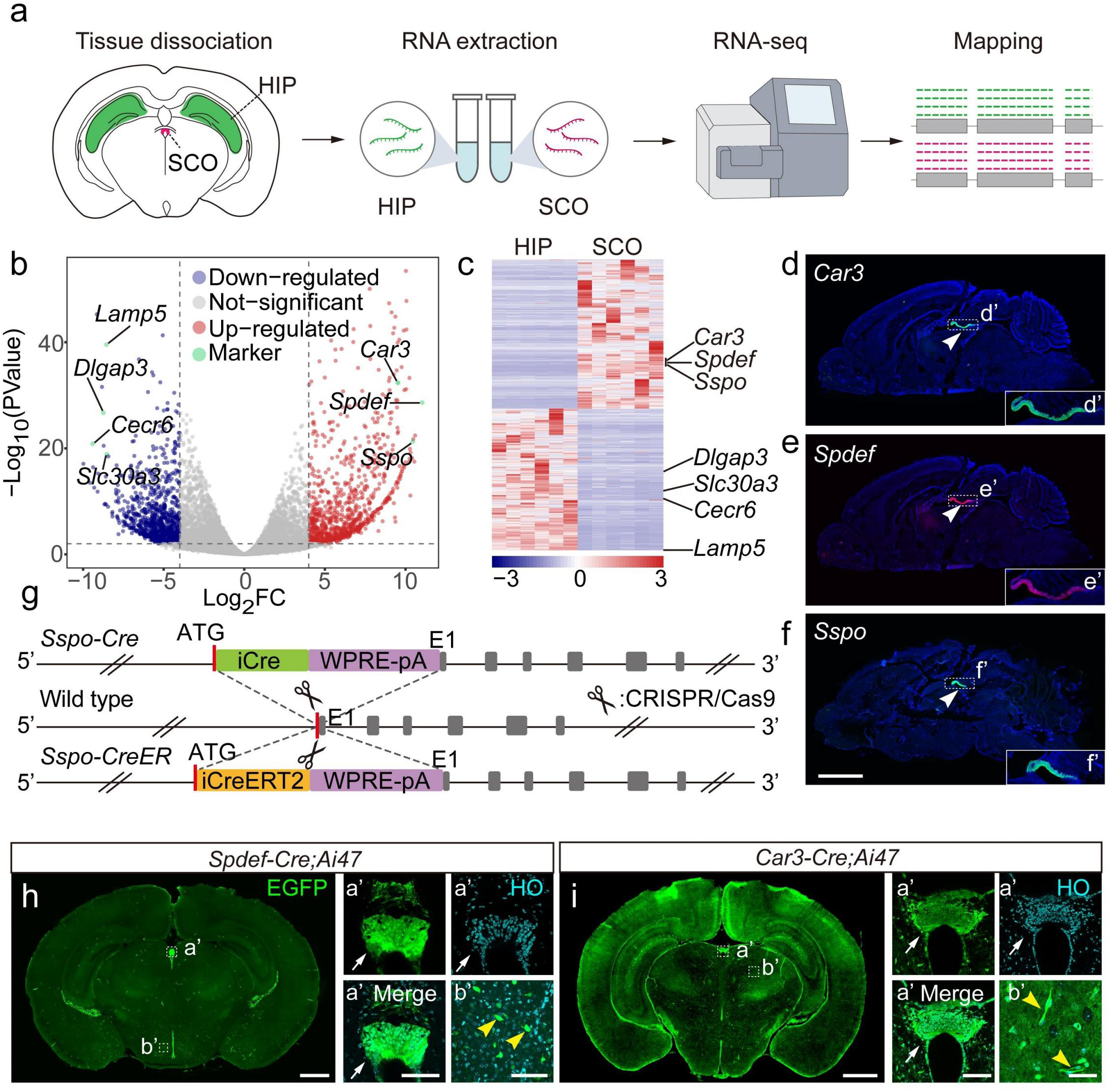
Identifying genes expressed specifically in SCO cells and developing corresponding Cre/CreER mouse lines. **a,** Strategy for screening genes for which expression was enriched in SCO cells compared with cells from the hippocampus (HIP). The SCO and HIP of adult mice (postnatal days 56– 70) were excised for bulk RNA-seq and analysis. **b, c,** Analysis of differentially expressed genes (upregulated or downregulated) in each of the SCO and HIP. b, Volcano plot; c, Clustered heatmap. *Sspo*, *Spdef*, and *Car3* were among the most highly specifically upregulated genes identified in the SCO. *Dlgap3*, *Slc30a3*, *Cecr6*, and *Lamp5* were four representative genes significantly upregulated in the HIP but downregulated in the SCO. Scale indicates fold expression level. **d–f,** Fluorescence imaging of mRNAs transcribed from *Car3* (d), *Spdef* (e), and *Sspo* (f) as determined with RNAscope mFISH of the adult mouse brain. Green, pseudocolor of Opal^TM^ 520; red, pseudocolor of Opal^TM^ 570; blue, nuclei stained by Hoechst 33342 (HO). Arrowheads in d–f indicate boxed areas enlarged in insets d’–f’. Images in d and e were from the same brain section stained with two probes. **g,** Strategy for generating mouse strains *Sspo-Cre* and *Sspo-CreER*. **h, i,** EGFP expression profiles for cells in the brain of *Spdef-Cre;Ai47* (h) and *Car3-Cre;Ai47* mice (i). Green, cells expressing EGFP; light blue, nuclei labeled by Hoechst 33342 (HO). Arrows in (h), SCO; yellow arrowheads in (h), neurons in the hypothalamus; white arrows in (i), SCO; yellow arrowheads in (i), blood vessels. a’, b’, insets. Scale bars, 2 mm (d–f), 1 mm (h–i) and 100 μm (h–a’, h–b’, i–a’, i–b’).

To further assess SCO-specific gene expression and evaluate SCO function, we generated three Cre transgenic mouse lines: *Car3-Cre*, *Spdef-Cre*, and *Sspo-Cre* (Fig. 1g and Supplementary Fig. 2a). To manipulate gene expression in the SCO from mice at different developmental stages, we also generated mouse line *Sspo-CreER* (Fig. 1g). For each line, a part of the sequence of the corresponding gene was replaced with Cre; thus, the expression level and profile of Cre reflected that of the inserted gene. To evaluate the specificity of these mouse lines, mice were crossed with one of two reporter mouse lines, namely *Ai47* (LSL-Rosa-EGFP) or *Ai14* (LSL-Rosa-tdTomato)^24,25^. SCO cells were labeled with EGFP in *Spdef-Cre;Ai47* mice, whereas only a small percentage of neurons were labeled (Fig. 1h). *Car3-Cre;Ai47* mice could also be used for SCO labeling, but some blood vessels expressed EGFP (Fig. 1i). We observed not only SCO cells but also some choroid-plexus cells and ependymal cells labeled in both *Spdef-Cre;Ai47* and *Car3-Cre;Ai47* mice (Supplementary Fig. 2b). After crossing *Sspo-CreER* mice with *Ai47* mice to induce Cre-dependent EGFP expression in *Sspo-*expressing cells via administration of tamoxifen during embryonic to adolescent stages, *Sspo* was expressed only in SCO cells from the embryonic stage (Fig. 2a, b, f and Supplementary Fig. 3) to adolescent stage (Fig. 2c–e). In addition, no Cre was expressed in the spinal cord. In *Sspo-Cre;Ai47* mice (Fig. 2g), EGFP also was highly expressed in the SCO. We further stained brain sections from *Sspo-Cre;Ai47* and *Sspo-CreER;Ai47* mice with an antibody specific for Foxj1, S100β, or β-catenin for labeling ependymal cells or choroid-plexus cells (Fig. 2 and Supplementary Fig. 3, 4). No EGFP^+^ ependymal or choroid-plexus cells were detected in brain sections of *Sspo-CreER;Ai47* mice (Fig. 2f and Supplementary Fig. 3). In *Sspo-Cre;Ai47* mice, only 0.09% of the ependymal cells and 0.33% of the choroid-plexus cells were EGFP^+^ (Fig. 2h, i and Supplementary Fig. 4). However, EGFP^+^ cells were not detected in ependyma around the 3^rd^ and 4^th^ ventricles or central canal (Supplementary Fig. 3, 4). SCO-spondin has been reported to be expressed in this area in some species^26^; however, no green fluorescence was detected in the floor plate of *Sspo-Cre;Ai47* mice (E12 to E18) (Supplementary Fig. 5). These results indicated that mouse lines *Sspo-Cre* and *Sspo-CreER* were ideal for SCO genetic manipulation.

**Fig. 2.**
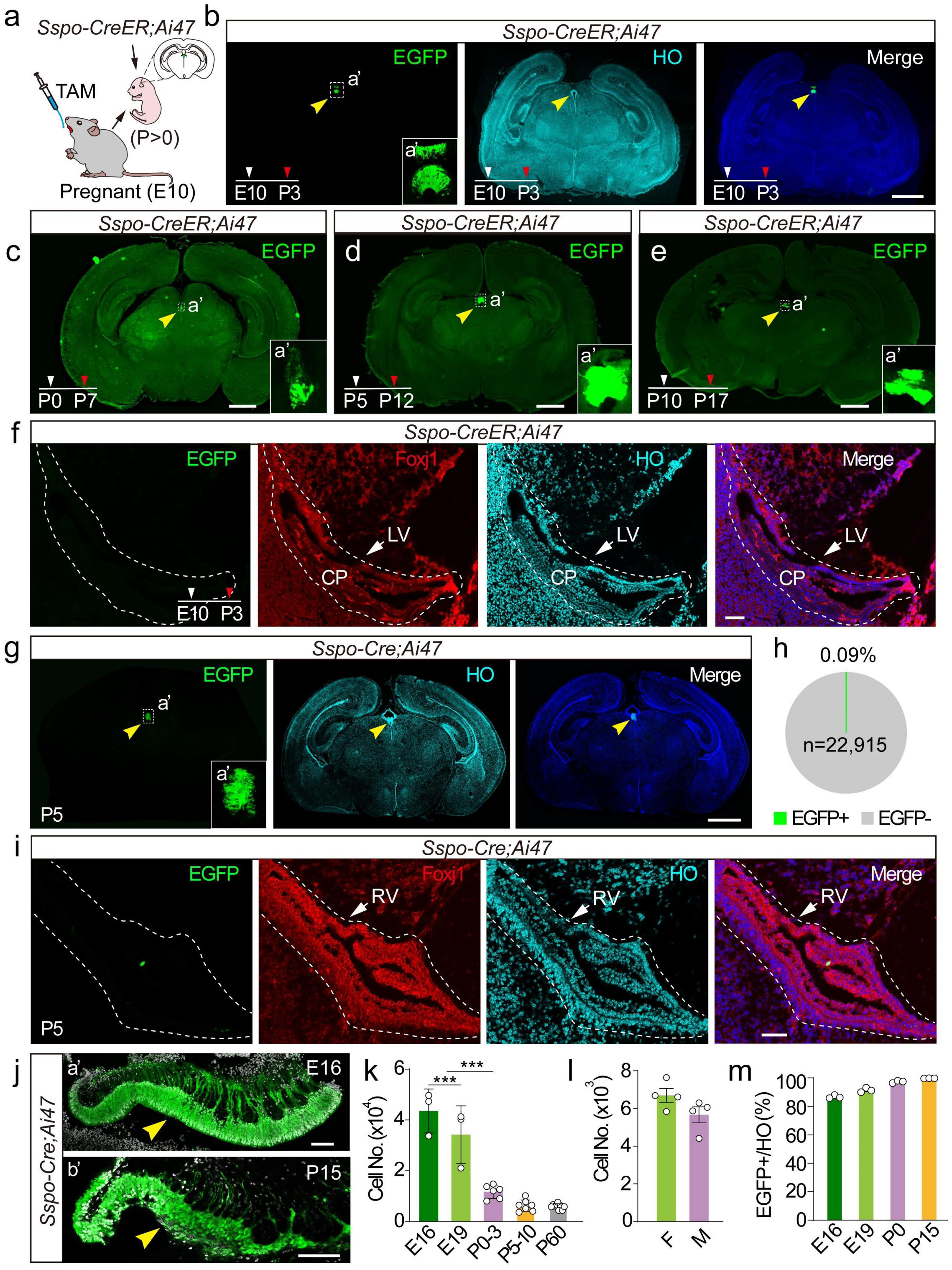
Specificity of *Sspo-CreER* and *Sspo-Cre* mouse strains for SCO labeling. **a,** Procedure for administering tamoxifen (TAM) to E10 *Sspo-CreER;Ai47* mice. **b–e,** EGFP expression profiles from the brain of *Sspo-CreER;Ai47* mice after tamoxifen was administered at E10, P0, P5, or P10; brains were collected at P3, P7, P12, and P17, respectively. Yellow arrowheads, SCO; a’, insets. **f,** Image of lateral ventricle from the brain of a *Sspo-CreER;Ai47* mouse at P3 after tamoxifen was administered at E10. Green, EGFP; red, ependymal cells stained with anti-Foxj1; LV, lateral ventricle; CP, choroid plexus; dashed lines, LV margin; **g,** EGFP expression in the brain of a *Sspo-Cre;Ai47* mouse at P5. Green, EGFP; yellow arrowheads, SCO. a’, insets. **h,** Percentage of ependymal cells with EGFP fluorescence in *Sspo-Cre;Ai47* mice at P5 (n = 22,915 ependymal cells in total). **i,** Images of LV from the brain of a *Sspo-Cre;Ai47* mouse at P5. Green, EGFP fluorescence; red, ependymal cells stained with anti-Foxj1; dashed lines, LV margin. **j,** Morphology of SCO in sagittal brain sections of *Sspo-Cre;Ai47* mice at E16 (a’) and P15 (b’). Green, EGFP; white, nuclei stained by Hoechst 33342 (HO); yellow arrowheads, SCO. **k,** Number of SCO cells in mice of different ages (n = 3–8 per group) (E16 and E19, n = 3 mice per group; P0–3, n = 6 mice; P5–10, n = 7 mice; P60, n = 8 mice). **l,** Number of SCO cells in the brain of adult male (M) and female (F) mice (n = 4 mice per group). **m,** Percentage of SCO cells exhibiting EGFP fluorescence in *Sspo-Cre;Ai47* mice (E16–P15, n = 3 mice per group). Scale bars, 1 mm (b–e, g), 100 μm (j–b’) and 50 μm (f, i, j–a’). Two-tailed unpaired *t-*test, *\*\*\*p* < 0.001. Error bars, s.e.m. White arrowheads, the starting point of tamoxifen injection; red arrowheads, time points for mouse sacrifice (b–f); green, EGFP fluorescence (b–j); light blue, blue or white, nuclei stained by Hoechst 33342 (HO, b, f, g, i, j).

The mouse SCO is first recognizable on embryonic day (E)10–11^27^, but it is unclear how the number of SCO cells changes during brain development. Therefore, we counted SCO cells at different embryonic and postnatal stages based on Hoechst 33342 staining and EGFP expression in wild-type and *Sspo-Cre;Ai47* mice (Fig. 2j and Supplementary Fig. 6). After postnatal day 5 (P5), the number of SCO cells decreased to ∼6,000 and remained relatively small thereafter, with no gender difference (Fig. 2 k, l). For *Sspo-Cre;Ai47* mice at different developmental stages, 86–99% of SCO cells were EGFP^+^ (Fig. 2m). Further, use of *Sspo-CreER;Ai47* mice to sparsely label SCO cells with a relatively lower dosage of tamoxifen revealed that most EGFP^+^ cells sent their processes to the 3^rd^ ventricle and were in contact with CSF (Supplementary Fig. 7). This indicated that most SCO cells were specialized ependymal cells.

### Genetic ablation of SCO cells leads to brain abnormalities

To study SCO function in the brain, the SCO was subjected to genetic ablation by crossing *Sspo-Cre* or *Sspo-CreER* mice with Cre-dependent *DTA* mice. Cre was expressed only in SCO cells of *Sspo-CreER* and *Sspo-Cre* mice (Fig. 2). In *Sspo-Cre;DTA* mice, the SCO was experimentally ablated from E11 to E12. In *Sspo-CreER;DTA* mice, ablation was performed at different developmental stages depending on the time of tamoxifen injection. As a result, *Sspo-CreER;DTA* and *Sspo-Cre;DTA* mice could be used to evaluate the role of the SCO with or without tamoxifen application, respectively (Fig. 3a), enabling targeted SCO ablation in mice at different embryonic or postnatal stages. Actually, we did not detect a difference in cortical layer structures, cortical cell density and body weight between the wild-type and *Sspo-Cre*/*Sspo-CreER* heterozygous mice (Supplementary Fig. 8), and thus these mice were used as the control. However, *Sspo-Cre;DTA* mice were significantly smaller than wild-type mice (Fig. 3b, c). Overall survival of *Sspo-Cre;DTA* mice dropped dramatically from P8, and ∼80% of mice died before P24 (Fig. 3d). Furthermore, *Sspo-Cre;DTA* mice that underwent SCO ablation had a bulged head and enlarged brain (Fig. 3e–g). Labeling of cellular nuclei with Hoechst 33342 revealed significant enlargement of the lateral ventricles in *Sspo-Cre;DTA* mice (Fig. 3h, i), and nearly all SCO cells were lost by P22 (Fig. 3h, i). Furthermore, SCO-spondin was present in CSF of control neonatal pups (P0–P3) but absent in CSF of *Sspo-cre;DTA* mice. Moreover, RF was detectable in the central canal of *Sspo-Cre* mice but was absent in *Sspo-Cre;DTA* mice. These results documented that the SCO was ablated in *Sspo-Cre;DTA* mice (Supplementary Fig. 9, 10 and Supplementary Table 1). To further understand how SCO ablation caused brain developmental abnormalities, magnetic resonance imaging (MRI) was carried out with *Sspo-Cre;DTA* mice at several postnatal stages. These mice exhibited clear progression of hydrocephalus with significant enlargement of ventricles (Fig. 3j–l). We used anti-S100β to stain ependymal cells and anti-β-catenin to stain choroid-plexus cells in control and *Sspo-Cre;DTA* mice (P0 and P12), revealing no differences in cell density or morphology of ependyma and choroid plexus, and the ependymal cells in the aqueduct and central canal of *Sspo-Cre;DTA* mice were well-aligned, cilia-rich, and possessed complete cellular structures (Supplementary Fig. 11, 12). Moreover, there were no apparent differences in the ventricular foramina, cerebral (Sylvian) aqueduct or central canal between wild-type and *Sspo-Cre;DTA* mice at P0, P3, P6 or P12 (Supplementary Fig. 13). Therefore, the hydrocephalus defects after SCO ablation were possibly non-communicating.

**Fig. 3.**
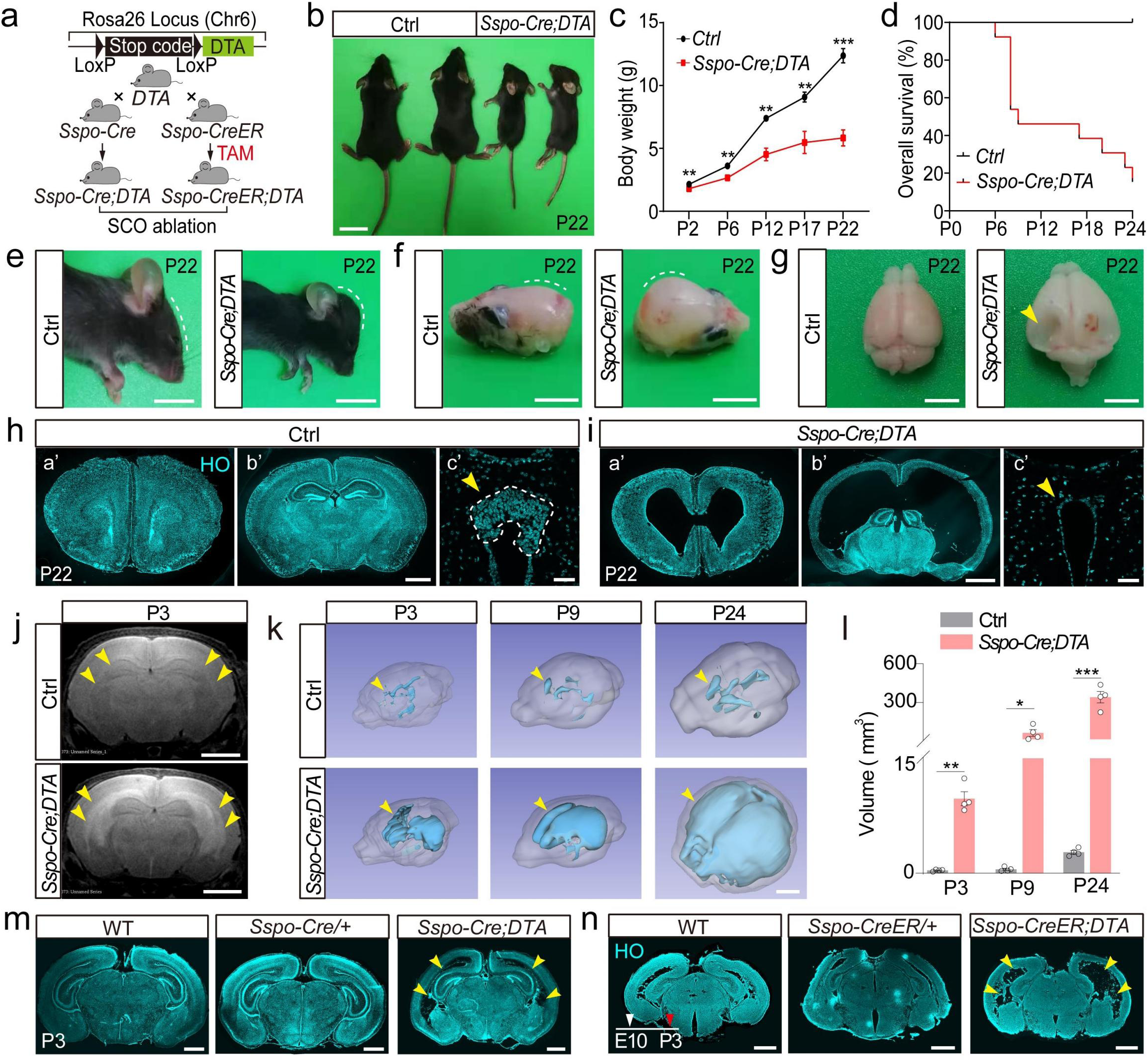
Phenotypes observed in SCO-ablated mice. **a,** Breeding strategy for *Sspo-Cre;DTA* and *Sspo-CreER;DTA* mice. **b,** Images of wild-type (Ctrl, P22, left) and *Sspo-Cre;DTA* mice (P22, right). **c,** Body weight of *Sspo-Cre;DTA* (red) and control mice (black) at P2, P6, P12, P17, and P22. n = 9 (P2), 9 (P6), 7 (P12), 6 (P17), 5 (P22, *Sspo-Cre;DTA*) and 6 (P22, Ctrl). **d,** Overall survival of *Sspo-Cre;DTA* mice (red, n = 13) and control mice (black, n = 13). **e–g,** Images of the head and brain of *Sspo-Cre;DTA* and control mice at P22. Dashed lines, skull curvature; arrowhead, collapsed cerebral cortex in the brain of a *Sspo-Cre;DTA* mouse. **h,** Images of coronal brain sections (a’, b’, different locations) from control mice at P22. Dashed line (c’), SCO margin; yellow arrowheads (c’), location of the SCO; light blue, nuclei stained with Hoechst 33342 (HO). **i,** Images of brain sections (a’, b’) from *Sspo-Cre;DTA* mice at P22. Yellow arrowhead (c’), location of ablated SCO; light blue, nuclei stained with Hoechst 33342 (HO). **j–k,** Representative MRI images (j, P3) and 3D reconstructed images (k, P3, P9, and P24) of *Sspo-Cre;DTA* and control mice. Yellow arrowheads (j, k), lateral ventricle; translucent white, 3D outline of the whole brain; blue, 3D outline of the lateral and third ventricles. **l,** Volumes of ventricles in control (Ctrl) and *Sspo-Cre;DTA* mice at P3, P9, and P24, n = 4 mice per group). **m, n,** Representative coronal brain sections of *Sspo-Cre;DTA* (m), *Sspo-CreER;DTA* (n) mice and corresponding controls. Tamoxifen was administered at E10, and mice were sacrificed at P3. WT, wild type; light blue, nuclei stained by Hoechst 33342 (HO); yellow arrowheads, lateral ventricles; white arrowhead, the starting point of tamoxifen injection; red arrowhead, the time point for mouse sacrifice. Scale bars, 1 cm (b), 5 mm (e–g), 2 mm (j, k), 1 mm (h–a’ and b’, i–a’ and b’, m, n), and 50 μm (h–c’ and i–c’). Two-tailed unpaired *t-*test, \**p* < 0.05, \*\**p* < 0.01, *\*\*\*p* < 0.001. Error bars, s.e.m.

To determine whether the SCO contributes to embryonic brain development, we induced *Sspo-CreER;DTA* with tamoxifen from E10 and observed similarly expanded ventricles at P3 (Fig. 3n), indicating that the SCO contributes to fetal brain development. Initiation of SCO ablation at P3 and brain collection at P37 revealed a slight expansion of the 3^rd^ ventricle (Supplementary Fig. 14). However, when SCO ablation was initiated at P25 and brains collected at P70, we did not observe an apparent ventricular expansion in the brain (Supplementary Fig. 14). Further, there were no apparent differences in neuronal density, distribution, or morphology in the cerebral cortex (Supplementary Fig. 15, 16). These results suggested that the SCO plays a critical role in early brain development.

### SCO contributes to neuronal development *in vivo*

Studies with chicken embryos have shown that the SCO promotes both neuronal adhesion and neurite growth *in vitro*^21,28^. To determine how SCO ablation would alter neuronal development in mice, neurons or neuronal progenitors were stained with an antibody against NeuN, SATB2, CTIP2 or TBR1 in *Sspo-Cre;DTA* mice (Fig. 4a–j). In the P3 brain, cortical neuron loss was slight (Fig. 4a, b), whereas loss was much greater at P22, at which time the six cortical layers could not be easily distinguished (Fig. 4c, d). Further, neuronal death increased significantly in the P3 brain but less so at P8 (Supplementary Fig. 17). Therefore, after SCO ablation, cortical neurons were gradually lost postnatally. Furthermore, the cortex was thinner at E18, when hydrocephalus had yet to manifest (Supplementary Fig. 18). Antibodies also were used for labeling neurons in different layers of the cerebral cortex of *Sspo-Cre;DTA* mice. These antibodies included SATB2 (layers II–VI, Fig. 4e–g), CTIP2 (layers V–VI, Supplementary Fig. 18), and TBR1 (layer VI, Fig. 4h–j). The numbers of SATB2^+^, TBR1^+^ and CTIP2^+^ neurons were significantly reduced in the cerebral cortex at E18 (Fig. 4e–j, Supplementary Fig. 18) and P22 (Supplementary Fig. 19).

**Fig. 4.**
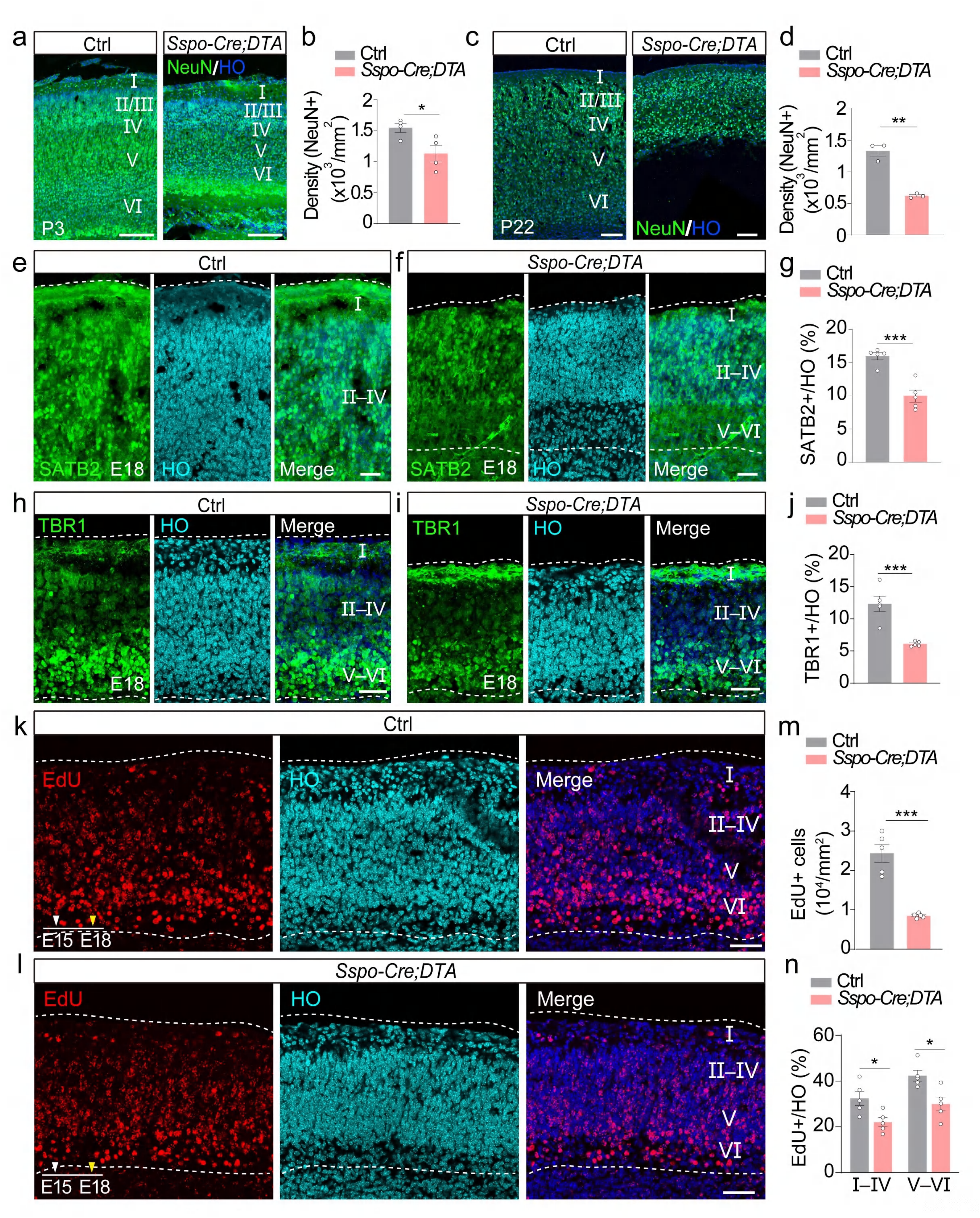
Neuronal properties of the mouse brain after SCO ablation. **a–d,** Images of coronal brain sections and density of NeuN^+^ cells from the cerebral cortex of *Sspo-Cre;DTA* and control mice at P3 (b, n = 4 regions per group) and P22 (d, n = 3 regions per group). Green, neurons stained with anti-NeuN; **e–f,** Images of coronal sections from the cerebral cortex of *Sspo-Cre;DTA* and control mice at E18. Green, neurons stained with anti-SATB2. **g,** Percentage of SATB2^+^ cells among total cells in the whole cerebral cortex from *Sspo-Cre;DTA* and control mice at E18 (n = 5 regions per group). **h–i,** Images of coronal sections from the cerebral cortex of *Sspo-Cre;DTA* and control mice at E18. Green, neurons stained with anti-TBR1. **j,** Percentage of TBR1^+^ cells among total cells in the whole cerebral cortex from *Sspo-Cre;DTA* and control mice at E18 (n = 5 regions per group). **k–l,** Images of coronal sections from the cerebral cortex of *Sspo-Cre;DTA* and control mice at E18. Red, cells stained with EdU. **m,** Density of EdU^+^ cells in the cerebral cortex of *Sspo-Cre;DTA* and control mice at E18 (n = 5 regions per group). **n,** Percentage of EdU^+^ cells among total cells of cortical layers I–V and V–VI from *Sspo-Cre;DTA* and control mice at E18 (n = 5 regions per group). Scale bars, 200 μm (a), 100 μm (c), 50 μm (e, f, h, i, k and l). Two-tailed unpaired *t-*test, \**p* < 0.05, \*\**p* < 0.01, *\*\*\*p* < 0.001. Error bars, s.e.m. Light blue or blue (a, c, e, f, h, i, k, l), nuclei stained with Hoechst 33342 (HO); I–VI (a, c, e, f, h, i, k, l), six layers of the cerebral cortex; white arrowheads (k, l), the time point of EdU injection; yellow arrowheads (k, l), the time point of mouse sacrifice; dashed lines (e, f, h, i, k, l), cortical margin.

Next, we injected pregnant *Sspo-Cre;DTA* mice once with 5-ethynyl-2’-deoxyuridine (EdU) at E15 and collected each pup brain at E18 for staining (Fig. 4k–n). The number of EdU^+^ cells in the cerebral cortex decreased dramatically (Fig. 4k–m). Further, the percentage of EdU^+^ cells was lower in both the superficial and deeper layers compared with wild-type mice (Fig. 4k, l, n). Finally, some NeuN^+^ neurons appeared in the white matter between layer VI and the subventricular zone of P3 mice (Supplementary Fig. 20). This observation indicated that a portion of neurons generated from the subventricular zone might not have successfully migrated to the appropriate cortical layer(s) during embryonic stages and more neurons underwent apoptosis postnatally in *Sspo-Cre;DTA* mice, which is consistent with the reduced number of neurons at P3. These results implied that neuronal properties may have been altered before hydrocephalus onset although hydrocephalus might have caused neuronal death in later developmental stages.

### SCO contributes to neuronal development and survival

To further examine the contribution of the SCO to neuronal development, neurons in transwells were co-cultured with a resected SCO (SCO^W^) or hippocampal tissue (Hip) or cultured alone (SCO^W/O^) (Fig. 5a). Neuronal density in the SCO^W^, Hip, and SCO^W/O^ groups was comparable before day 6 *in vitro* (DIV6) but then gradually decreased in the SCO^W/O^ and Hip groups. At DIV13, the density was significantly greater in the SCO^W^ group, indicating the SCO could maintain neuronal survival (Fig. 5b, c). This result was consistent with observations made in the cortex of *Sspo-Cre;DTA* mice (Fig. 5i–q). Incubation of neurons with anti-Tuj1 (neurite marker) revealed reduced complexity of neurites of the SCO^W/O^ and Hip groups (Fig. 5d). To evaluate neuronal morphology *in vitro*, neurons were infected with Lentivirus-H1-EGFP (Fig. 5e). The number of neurites (including dendrites and axons) were approximately doubled in the SCO^W^ group (Fig. 5f–h), suggesting that the SCO promotes neurite growth.

**Fig. 5.**
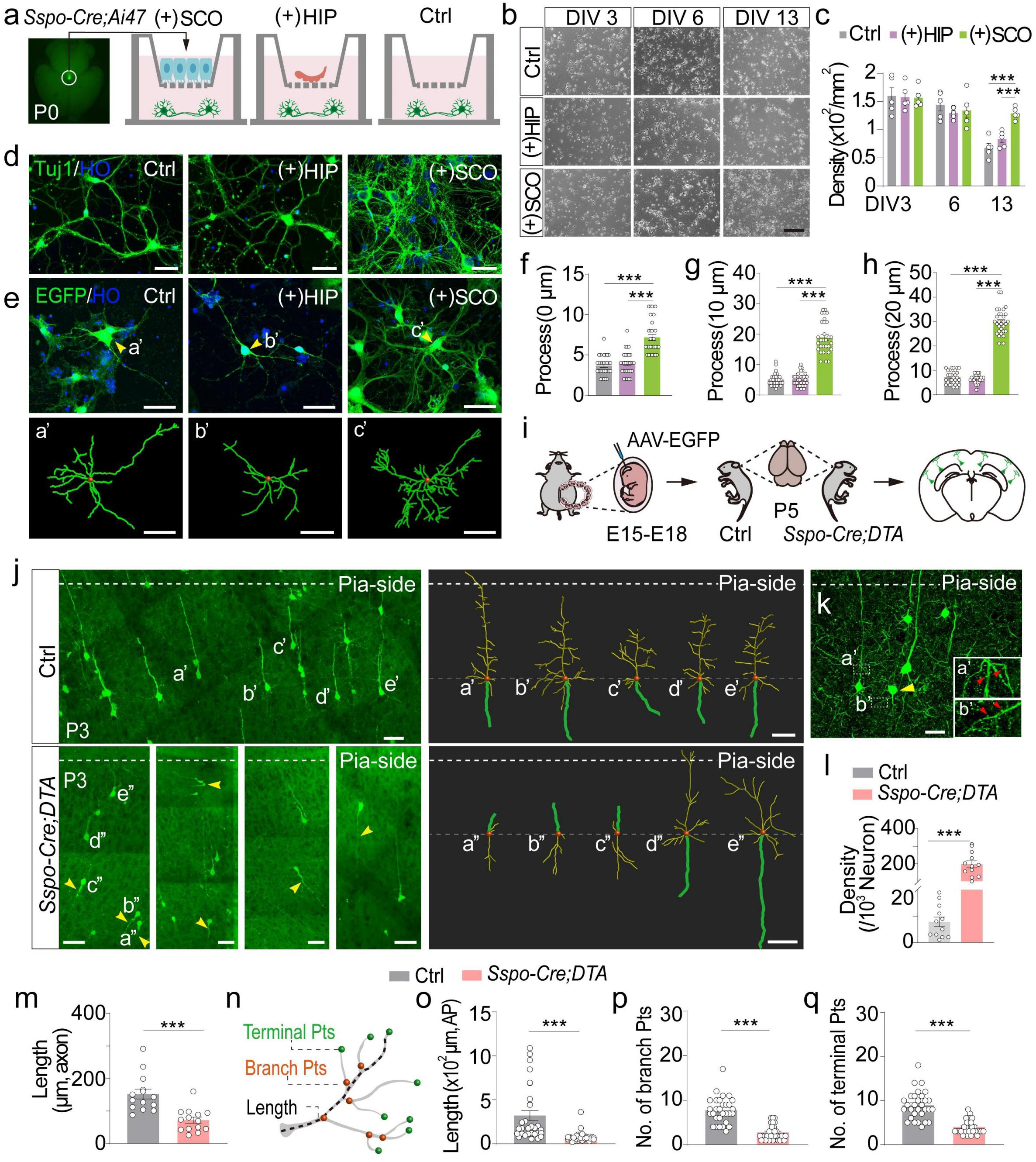
SCO promotes neuronal development. **a,** Strategy for culturing neurons in the absence or presence of the SCO. The SCO was resected from *Sspo-Cre;Ai47* pups. HIP, hippocampus. **b, c,** Images and density of neurons co-cultured with SCO (lower), hippocampus (middle, HIP), or alone (upper) at DIV3, 6 and 13 (n = 5 wells per group). **d**, **e**, Morphology of neurons. Green, neurons stained with anti-Tuj1 (d) or EGFP in neurons infected by Lentivirus-H1-EGFP (e); blue, Hoechst 33342 (HO, nuclei). a’–c’, Morphology of three representative neurons in three groups. **f–h,** Number of neurites after Lentivirus-H1-EGFP injection, located at the soma (f, 0 μm) or 10 μm (g) or 20 μm away from the soma (h). n = 25 neurons per group. **i,** Procedure for sparse neuronal labeling. AAV-CAG-EGFP was injected at E15–18, and the brains were collected at P5. **j,** Morphology of neurons in the brain of *Sspo-Cre;DTA* and control mice. a’–e’, a’’–e’’, representative neurons in control and *Sspo-Cre;DTA* mice, respectively. a’’–c’’, abnormal neurons with apical dendrites directed toward the corpus callosum and axons facing the pial surface; arrowheads, neurons with abnormal polarity; soma, red; axons, green; dendrites, yellow. **k,** Morphology of sparsely labeled neurons in the brain of a *Sspo-Cre;DTA* mouse. Green, anti-GFP; yellow arrowheads, neurons with abnormal polarity; red arrowheads, spines of a neuron with normal (a’) or abnormal (b’) polarity. **l, m,** Density of neurons with apical dendrites facing the side of the corpus callosum (n = 12 regions per group) and the length of axons (n = 14 neurons per group) in *Sspo-Cre;DTA* and control mice at P5. **n–q,** Analysis of apical dendrites at P5 including the length of apical dendrites (o), and the number of branch points (p) and terminal points (q). n = 30 neurons per group. Schematic (n) showing the method for measuring the length of an apical dendrite (dashed line), branch points (Branch Pts, orange), and terminal points (Terminal Pts, green). Scale bars, 100 μm (b) and 50 μm (d, e, j, k). Two-tailed unpaired *t-*test, *\*\*\*p* < 0.001. Error bars, s.e.m.

Previous results from SCO co-culturing and grafting studies have established that the SCO promotes neuronal development^20,21^, but the impact of SCO ablation on neuronal development *in vivo* remains largely unknown. To address this issue, a low dose of adeno-associated virus (AAV)-CAG-EGFP was injected into the lateral ventricle of *Sspo-Cre;DTA* mice as well as controls (*DTA*, *Sspo-Cre*, and wild type) at E15–18. Neuronal morphology in the cerebral cortex was then assessed (Fig. 5i). At P5, the *Sspo-Cre;DTA* brain had a high percentage of cortical neurons with abnormally polarized axons and apical dendrites (Fig. 5j–l). A 3D reconstruction revealed that apical dendrites were shorter in *Sspo-Cre;DTA* mice (Fig. 5n, o). *Sspo-Cre;DTA* neurons also had fewer branches, with abnormally polarized axons (Fig. 5m, n, p, q). We further stained brain sections with anti-EGFP to observe spines in dendrites and boutons in axons to establish neuron polarity in the cerebral cortex. Neurons indeed had inverted polarity in *Sspo-Cre;DTA* mice (Fig. 5k). These results demonstrated that the SCO promotes neuronal survival and development both *in vitro* and *in vivo*.

### Peptides secreted by SCO promote neuronal development

To elucidate the molecular mechanism underlying SCO regulation of neuronal development, RNA sequencing and mass spectrometry (MS)-based peptidomics were carried out to assess the involvement of SCO-derived secretory molecules. EGFP**^+^** SCO cells in the brain of *Sspo-CreER;Ai47* mice sent processes to the third ventricle (Fig. 6a, Supplementary Fig. 7). Golgi were abundant in SCO cells adjacent to the third ventricle (Fig. 6a), indicating that SCO cells are highly secretory cells. Transcriptome analysis of each of SCO and hippocampus revealed that the positive regulation of peptide-hormone secretion pathway was highly enriched in the SCO (Fig. 6b and Supplementary Fig. 21). We thus searched for SCO peptides that might contribute to neuronal development. A systems-level peptidomic analysis of SCOs from 120 mouse brains and corresponding hippocampal tissue (Fig. 6c) identified 2,788 (hippocampus) and 3,588 (SCO) *de novo* peptide-associated features (Fig. 6d, e). Interestingly, the vast majority (∼92%) of features found in SCO samples were unknown compared with only ∼12% for the hippocampus (Fig. 6d–e). The SCO features were further searched using the peptide databases Neuropep and SwePep, resulting in the annotation of 12 mature peptides that were highly enriched in the SCO (Supplementary Table 2). Of these, 9 peptides exhibited 100% sequence coverage, including manserin (Fig. 6f), which regulates neuroendocrine pathways. To our knowledge, this peptide has not previously been identified in mice. The preprohormones of these 12 peptides belong to a variety of subfamilies, including the thymosin beta (Tβ), granin (chromogranin/secretogranin), melanin-concentrating hormone, opioid, and somatostatin families (Fig. 6g). Among these, the Tβ family is related to embryonic development^29,30^. Moreover, Tβ10 and Tβ4 showed unique post-translation modifications and glycosylation, with one O-glycosylated peptide from Tβ10 and eight from Tβ4. Comparison with glycosylated peptides identified in the hippocampus (control) sample revealed four O-glycosylated peptides from Tβ4 that were specific to the SCO. Therefore, we selected Tβ4 and Tβ10 for further investigation along with a previously unidentified bioactive 24-residue peptide that we named NP24 (DVGSYQEKVDVVLGPIQLQSPSKE), detected with high cleavage probability (*p* = 0.9906) in the SCO group.

**Fig. 6.**
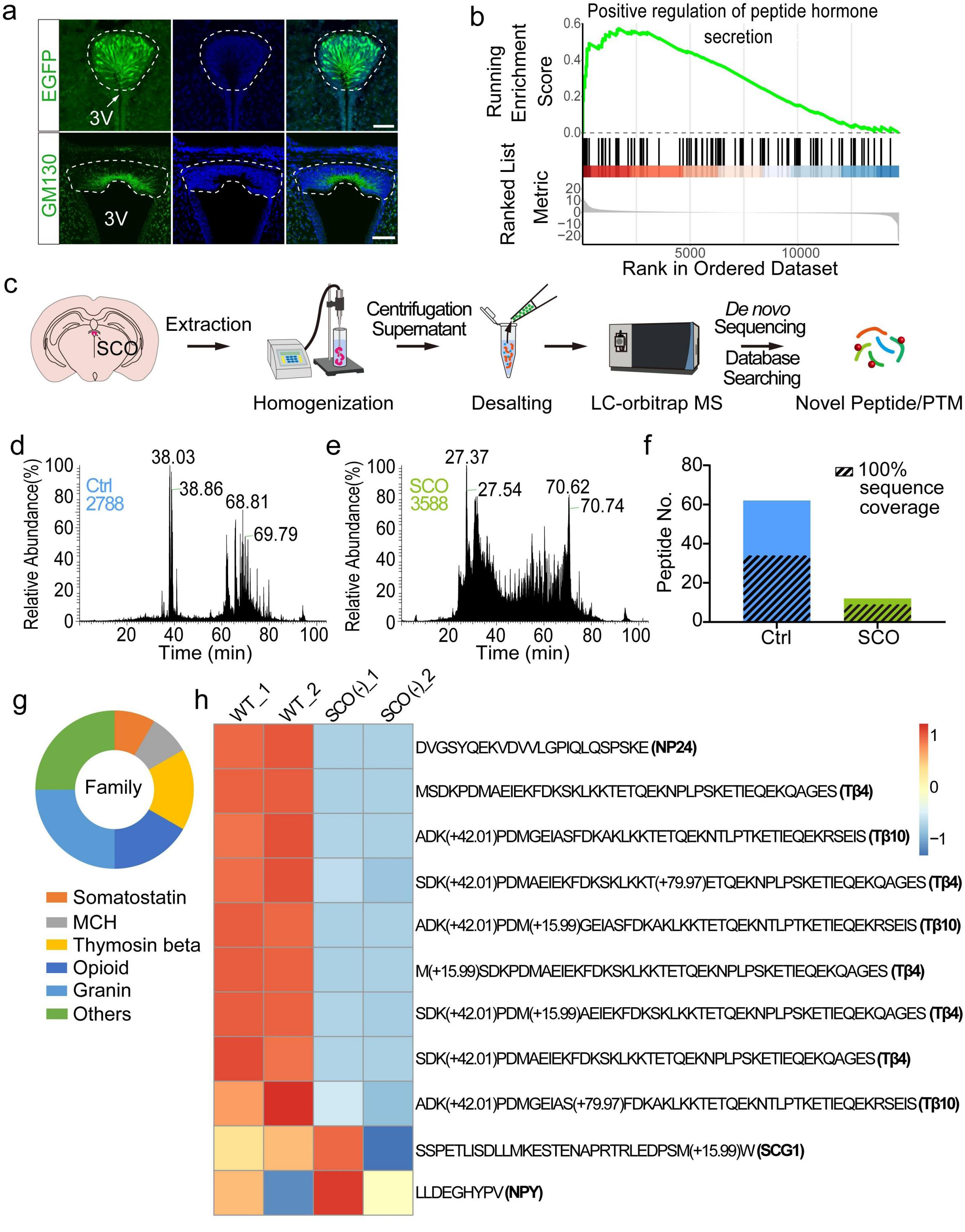
Detection of neuropeptides in SCO and their release into cerebral spinal fluid. **a,** Morphology of individual SCO cells with polarized long processes extending to the third ventricle (3V, upper row) and gathered Golgi enriched in terminals of polarized processes (GM130, lower row). Dashed lines, SCO margin; green (upper row), EGFP fluorescence of SCO cells in *Sspo-CreER;Ai47*; green (lower row), Golgi labeled with anti-GM130; blue, nuclei labeled by Hoechst 33342 (HO). **b,** Gene Set Enrichment Analysis showing that the pathway ‘Positive regulation of peptide hormone secretion’ was enriched in the SCO. Transcriptomic data were from samples of the SCO and control hippocampus of P0 mice. **c**, Workflow for peptide extraction and peptidomic analysis with LC-MS/MS on an Orbitrap MS platform. MS, mass spectrometry; LC-MS/MS, liquid chromatography-tandem mass spectrometry. **d, e,** Peptidomic sequencing results from tandem mass spectra of the control samples (hippocampus, d) and SCO samples (e) using a *de novo* sequencing strategy. Peptide features totaling 2,788 and 3,588 were identified in the SCO and control samples, respectively. **f, g,** Number of peptides identified in the SCO and control samples (f, Ctrl, 62 peptides with 34 of them exhibiting 100% sequence coverage; SCO, 12 peptides with 9 of them exhibiting 100% sequence coverage) and peptide family distribution of identified SCO peptides (g). MCH, melanin-concentrating hormone. **h,** SCO-secreted Tβ4, Tβ10 and NP24 in cerebral spinal fluid of wild-type (WT) and *Sspo-Cre;DTA* mice at E18–P0. WT_1 and WT_2, two samples of mixed cerebral spinal fluid of WT mice (∼30 pups or embryos per sample); SCO (-)_1 and SCO (-)_2, two samples of mixed cerebral spinal fluid of *Sspo-Cre;DTA* mice (∼30 pups or embryos per sample). Secretogranin-1 (SCG1) and neuropeptide Y (NPY) were detected as internal controls. Scale bars, 50 μm (EGFP, a) and 100 μm (GM130, a).

Next, we collected and pooled CSF from 20–30 wild-type or *Sspo-Cre;DTA* perinatal mice (E18–P0); peptides Tβ4, Tβ10 and NP24 were detected in wild-type CSF (Supplementary Fig. 22). Notably, the concentrations of Tβ4 and Tβ10 were substantially lower in CSF from *Sspo-Cre;DTA* mice (Fig. 6h, Supplementary Table 3). Notably, the concentration of five different Tβ4 segments was only 0–13% of that measured in wild-type CSF, and the concentration of Tβ10 was only 0–4% that of wild-type. NP24 was not detected in CSF from *Sspo-Cre;DTA* mice. These results demonstrated that SCO is the main source of these three peptides in CSF. Indeed, the results indicated a potential additional source of Tβ4 and Tβ10 in the CSF of brain and that nearly all NP24 is derived from SCO.

The potential roles of these three SCO peptides in neuronal development were examined by applying Tβ4, Tβ10, and NP24 to primary cultured neurons, which significantly increased neuron density and prolonged their survival (Fig. 7a, b and Supplementary Fig. 23) and prolonged their survival. In addition, the peptide-treated neurons had more complex morphology, with longer processes and more branches and terminal points (Fig. 7c–e and Supplementary Fig. 23). Finally, RNA-seq of neurons incubated for 2 days with these peptides revealed that they promoted neuronal development via activation of specific pathways (Supplementary Fig. 24).

**Fig. 7.**
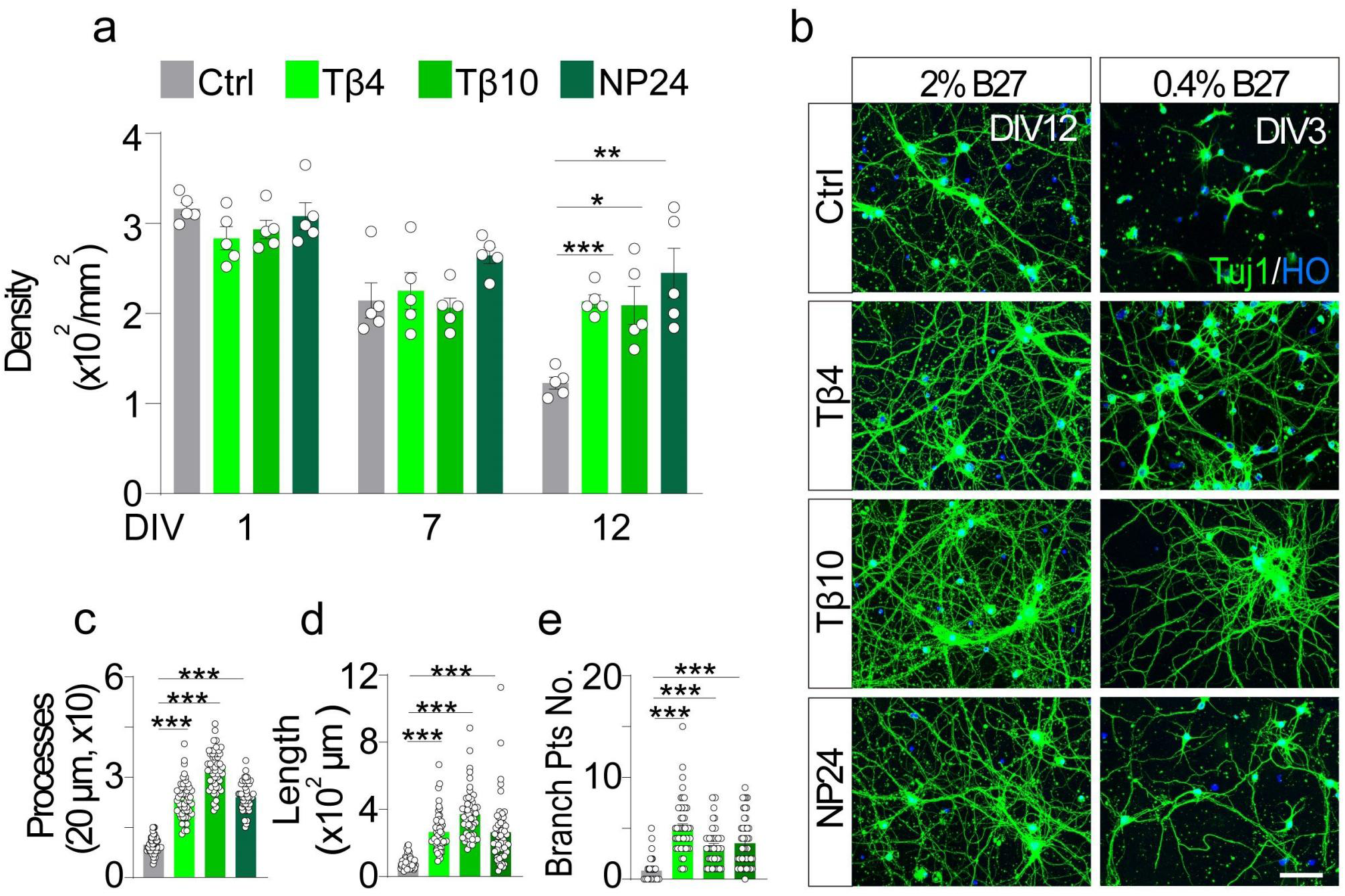
SCO-secreted peptides promote neuronal development *in vitro*. **a,** Density of neurons cultured in the absence (Ctrl) or presence of Tβ4, Tβ10 and NP24 at DIV1, DIV7 and DIV12. n = 5 regions per group. **b,** Representative images of neurons in the absence or presence of Tβ4, Tβ10 and NP24 at DIV 12 supplied with 2% B27 (left) or at DIV3 supplied with 0.4% B27 (right). Green, neurons labeled with anti-Tuj1; blue, nuclei labeled by Hoechst 33342 (HO). **c,** Number of neurites at 20 μm away from the soma (DIV12) in cultured neurons. Neurons were cultured with 2% B27 in the absence or presence of Tβ4, Tβ10 or NP24. The points where the neurite intersected a dotted circle were counted. Ctrl, 9.80±2.45 µm; Tβ4, 23.26±5.53 µm; Tβ10, 32.42±5.84 µm; NP24, 24.02±2.26 µm (n = 50 neurons per group). **d, e,** Morphological analysis of neurons cultured in the absence or presence of Tβ4, Tβ10 or NP24. Parameters included the length of the apical dendrite (n = 50 neurons per group) and the number of dendritic branch points (n = 50 neurons per group) in neurons cultured with 0.4% B27. Pts, points. Scale bars, 50 μm (b). Two-tailed unpaired *t-*test, \**p <* 0.05, \*\**p <* 0.01, *\*\*\*p <* 0.001. Error bars, s.e.m.

To test whether application of the Tβ4-Tβ10-NP24 cocktail could rescue the phenotype of *Sspo-Cre;DTA* mice, the cocktail was injected into the lateral ventricle of mice once every 3 days from P0 (3 µg each time). The cocktail-treated mice survived significantly longer than did the saline-treated *Sspo-Cre;DTA* mice at P24 (overall survival 75% vs. 33.3%, respectively; Fig. 8a), and they also attained a higher mean body weight (Fig. 8b). MRI revealed that the cocktail-treated mice exhibited less severe ventricle expansion and reduced hydrocephalus (Fig. 8c–g), which was verified through staining cell nuclei in coronal brain sections (Fig. 8e–f). Notably, the ventricular area in the cocktail-treated mice was reduced by ∼60% compared with saline-treated controls (Fig. 8g). Further, the percentage of neurons with abnormally polarized axons and shorter dendrites was vastly reduced in cocktail-treated mice (Fig. 8h–j). These results indicated that SCO-secreted peptides might contribute to neuronal development and help mitigate abnormalities in SCO-ablated mice. We applied individual peptides (at two different concentrations) to the brain of *Sspo-Cre;DTA* mice for rescue tests; each of the three peptides could also partially rescue the phenotype observed in SCO-ablated mice including longer survival (Fig. 8k–m) and reduced ventricular volume (Fig. 8n–q). However, no individual peptide could fully rescue the phenotype, indicated that other peptides or proteins from SCO might also contribute to rescuing the *Sspo-Cre;DTA* phenotype.

**Figure 8.**
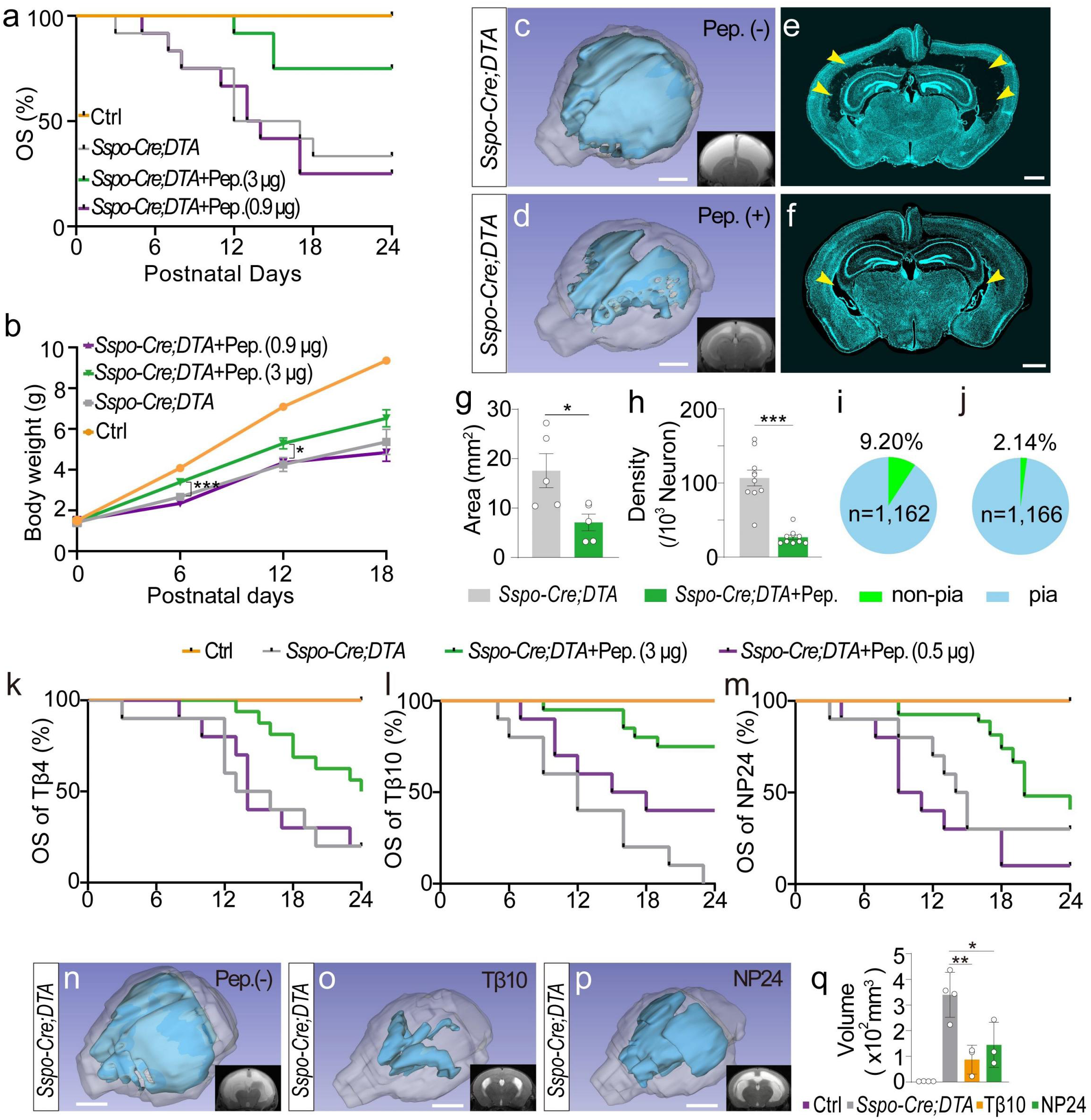
SCO-secreted peptides rescue phenotypes in SCO-ablated mice. **a,** Overall survival (OS) of control (orange), *Sspo-Cre;DTA* (gray) and *Sspo-Cre;DTA* mice injected with a cocktail of SCO peptides (Tβ4, Tβ10, NP24) comprising 0.9 µg (purple) total for all three peptides or 3 µg each time (green); n = 12 mice per group. Pep., peptide cocktail. **b,** Body weight of control (orange), *Sspo-Cre;DTA* (gray) and *Sspo-Cre;DTA* mice injected with the SCO peptide cocktail of 0.9 µg (purple) and 3 µg each time (green) at P0, P6, P12, or P18 (n = 8 for P0, P6 and P12 mice; n = 6 for P18). **c, d,** MRI and 3D reconstruction of the ventricle in *Sspo-Cre;DTA* mice without (c) or with (d, 3 µg) injection the SCO peptide cocktail at P24. **e–g,** Ventricular area measured from *Sspo-Cre;DTA* mice without (e) or with (f, 3 µg each time) injection of the SCO peptide cocktail at P24. n = 5 mice per group (g). Blue, Hoechst 33342 (HO, nuclei); arrowheads, lateral ventricles. **h–j,** Density (n = 10 regions per group) and percentage (i, *Sspo-Cre;DTA* without SCO peptide injection; j, *Sspo-Cre;DTA* mice with cocktail injection, 3 µg each time) of neurons exhibiting non-pia-projected polarization among all labeled neurons. Non-pia, neurons with axons directed toward the corpus callosum; pia, neurons with axons directed toward the pial surface. **k–m,** Overall survival (OS) of control (orange), *Sspo-Cre;DTA* (gray) and *Sspo-Cre;DTA* mice injected with 0.5 µg (purple) or 3 µg (green) of Tβ4 or Tβ10, or NP24 each time. Tβ4, n = 10, 10, 10, 16 mice; Tβ10, n = 10, 10, 10, 20 mice; NP24, n = 10, 10, 10, 27 mice. **n–q,** 3D reconstruction and quantification of ventricular volume of *Sspo-Cre;DTA* mice after injection of Tβ10 (o, 3 µg each time) or NP24 (p, 3 µg each time) or without injection of SCO peptides (n). q, n = 4 mice for control and *Sspo-Cre;DTA*; n = 3 mice for *Sspo-Cre;DTA+*Tβ10 and *Sspo-Cre;DTA+*NP24. Scale bars, 2 cm (c, d, n–p) and 1 mm (e, f). Two-tailed unpaired *t-*test, \**p <* 0.05, \*\**p <* 0.01, *\*\*\*p <* 0.001. Error bars, s.e.m.

## Discussion

The genetic mouse models we developed make it possible to ablate cells specifically in the SCO and thus address its function in the brain. Our findings suggest that SCO cells are *de facto* secretory cells, and indeed we identified three SCO-secreted peptides, namely Tβ4, Tβ10, and a new peptide NP24, that contribute to neurite development and neuronal survival *in vitro*.

SCO cells are bipolar: their apical poles face the third ventricle, whereas their basal processes face the subarachnoid space and contact local blood vessels^2,16^. Ultrastructural analysis has suggested that the SCO is a high-output secretory gland^3^. A number of proteins, including SCO-spondins, transthyretin, and probably fibroblast growth factor, have been reported to be released from SCO cells^3^. Our proteomic data revealed the presence of SCO-spondin in the CSF of control mice, whereas it was absent from the CSF of SCO-ablated mice. Several oligopeptides derived from the SCO-spondin sequence are bioactive, and their positive roles in neurite growth have been confirmed *in vitro*^21,31^. However, it is unknown whether these SCO-spondin oligopeptides are secreted from the SCO under physiological conditions. Although we detected known peptides in CSF, e.g., SCG1, NPY, and pro-SAAS, we did not detect secretory peptides related to SCO-spondin. Thus, SCO-spondin–derived peptides might be present at low concentrations or may be rapidly degraded by enzymes in CSF after release.

Tβ4, Tβ10 and NP24 are small peptides, and therefore they could potentially reach different cortical layers and act on developing neurons or neuronal progenitors directly through the CSF. We found that all three peptides promoted neuronal development through activation of specific pathways, including neurogenesis and neuronal survival. Some of these functions are similar to those of SCO-spondin, indicating they cooperate to promote growth and development.

The cocktail of peptides that we injected into mice for the rescue experiment included 1 μg of each peptide (i.e., 3 μg total). Platelets and leukocytes release Tβ4 into wound fluid with a final concentration of 13 μg/ml^32,33^. Given that the weight of the developing brain of a mouse pup is 0.2–0.3 g (P2–10), the estimated concentration of Tβ4 that we used in *Sspo-Cre;DTA* mice (1 μg/brain, i.e., 3–5 μg/ml) was comparable to the physiological concentration (13 μg/ml). Based on our liquid chromatography-MS (LC-MS) analysis of CSF, we estimated the following concentrations of the peptides: Tβ4, ∼5.7 μg/ml; Tβ10, ∼2.3 μg/ml; and NP24, ∼0.023 μg/ml (Supplementary Table 3). Considering that enzyme-mediated digestion is always a possibility in CSF and that the amounts of peptides might decrease during collection and preparation, their concentrations under physiological conditions are likely higher than what we measured.

SCO comprises two cell layers: the ependymal layer, which originates from the epithelium in direct contact with CSF of the third ventricle, and the hypendymal cells situated beneath the ependymal layers of the SCO^19,34^. We determined that most SCO cells were EGFP^+^ in *Sspo-Cre;Ai47* mice. In addition, the majority of EGFP^+^ cells in SCO were S100β^+^ or Foxj1^+^. The use of *Sspo-CreER;Ai47* mice for sparse labeling of SCO cells revealed individual cell morphologies and confirmed that most EGFP^+^ cells extended processes to the third ventricle, indicating their specialization as ependymal cells in direct contact with CSF.

Emerging evidence has linked abnormal neurodevelopment to hydrocephalus rather than fluid hydrodynamics^35–38^. Our results indicate that SCO ablation primarily leads to brain malformations during early development, resulting in a thinner cortex and hydrocephalus. The phenotype of our SCO-ablated mice aligns partially with observations in previous congenital or mutant animal models, such as hyh mice, SUMS/np mice, and H-Tx rats^4,8^. The SCO is often dysfunctional or malformed in these models, although they also exhibit dysfunction in non-SCO brain regions, making it uncertain whether the SCO is solely responsible for hydrocephalus. Our observations revealed a thinner cortex in *Sspo-Cre;DTA* mice at E18, before hydrocephalus onset. Our *in vitro* and *in vivo* investigations suggest that the SCO plays a vital role in neuronal and cortical development, a finding consistent with prior studies^20,21^. Nevertheless, some disparities exist between the phenotype observed in our SCO-ablation model and those documented in previous mutant models or animals subjected to SCO immunoblockade^39^. For instance, *Sspo-Cre;DTA* mice did not develop aqueductal stenosis after specific SCO ablation. Analysis of ependymal cells within the aqueduct revealed no discernible differences between control and *Sspo-Cre;DTA* mice. In homozygous *Sspo-Cre* and *Sspo-CreER* mice, which mimic *Sspo* gene knockout and lack SCO-spondin in their CSF, there was no obvious aqueductal stenosis or disruption of ependymal cells or differences in body weight or brain morphology.

The SCO-RF complex is involved in spinal-cord development in zebrafish ^12–14^. Some rodent models, including hyh and H-Tx, have brain abnormalities including the SCO and RF^6,11,40,41^. Our results for *Sspo-Cre;DTA* mice revealed apparent abnormalities in the cerebral cortex and lateral ventricles when the SCO was ablated at embryonic stages; however, there was no obvious phenotype in the brain or change in overall survival if the SCO was ablated after P25. These results indicate that the SCO plays a critical role in early brain development, and its role in adults might differ^3,9,20^. Previous studies indicated that the monoamine concentration in CSF increases sharply in the absence of RF ^9,42,43^. Further, dysfunction of the adult neurogenic niche also has been proposed and investigated by SCO-grafting^3^. These observations are consistent with our results that the number of SCO cells decreases during embryonic and postnatal development. We cannot exclude the possibility that this decrease is attributable to incomplete experimental ablation of the SCO in adult *Sspo-CreER;DTA* mice.

Although hydrocephalus is commonly attributed to primary defects in CSF homeostasis, emerging evidence suggests that congenital hydrocephalus may stem not solely from the overproduction of certain factors in CSF but rather from anomalies in neurodevelopment^38^. In addition, additional evidence has linked abnormal neurodevelopment to hydrocephalus rather than fluid hydrodynamics^35–38^. Moreover, disruptions in the cerebral cortex or neural stem cells can result in secondary ventricular enlargement without primary defects in CSF circulation^35,38,44–47^. Abnormal neurogenesis also can arise from defects in the ventricular zone in certain of the aforementioned animal models of congenital hydrocephalus^47^.

It was recently reported that deletion or mutation of Trim71 in mouse neuroepithelial cells causes prenatal hydrocephalus. Indeed, congenital hydrocephalus mutations can cause abnormalities in neuroepithelial cell differentiation and neurogenesis^38^. Results from human patients also indicate that genetic disruption of early brain development—rather than impaired CSF circulation—is the primary cause of sporadic congenital hydrocephalus ^35,36^. We observed that E18 *Sspo-Cre;DTA* mice experienced cortex thinning, i.e., preceding the onset of hydrocephalus. We posit that a diminution of SCO secretion capacity may lead to abnormalities in neurogenesis, neuronal development, and cortex formation that could result in secondary hydrocephalus. Although hydrocephalus in SCO-ablated mice may be linked to SCO-RF defects, concurrent defects in cortical development may contribute to postnatal ventricle expansion.

## Acknowledgement

We thank lab members in Ge laboratories, Dr. Bridget Samuels, and Dr. Tim Taylor for their feedback and critical reading of the manuscript. We thank Qingchun Guo, Xinwei Gao, Mingyue Jia in the imaging core of CIBR for assistance with imaging and data analysis. We thank Wenlong Li, Shufang Huang, and Ruirui Shen at the animal core facility for assistance with animal care and purchasing. We thank Jinmei Chen and Xuefang Zhang at the Genomics Center for sequencing and fluorescence-activated cell sorting. We thank the mass spectrometry facility of the Phoenix Center for use of their instrumentation. We thank Dr. Fen Huang for suggestions. This work was supported in part by grants from the STI2030-Major Projects (2022ZD0204700), the Natural Science Foundation of China, and Beijing Scholars to W.G. (32170964), startup funds from CIBR, the National Key R&D Program of China to C.J. (No. 2021YFA1302601), and a grant from the U.S. National Institutes of Health (NIH, R01DK071801) to L.L.. The Orbitrap instruments were purchased through support from an NIH shared instrument grant (NIH-NCRR S10RR029531).

## Author Contributions

W.G. conceived and supervised the project. T.Z., D.A., P.W., L.L., C. J., W.S., and W.G. designed experiments. T.Z. performed most experiments, including mouse genetics, cell ablation, neuronal culture, immunostaining, imaging, RNAscope mFISH, mouse breeding and genotyping, CSF collection, and rescue experiments, among others. P.W., Y.X., F.M., Y.Z., T.Z. and C.J. performed peptidomics and proteomics and data analysis. W.G. Z.Z., T.T. and X. Zhang performed SCO bulk RNA sequencing; C.X., D.A., and W.G. analyzed the RNA sequencing data. W.G. and T.Z. determined the genes for genetic mouse strains. T.Z. and J.L. assisted in MRI data acquisition and analysis, T.T. assisted in MRI imaging, Z.B. and T.Z. performed AAV viral labeling and peptide injection and assisted in slice imaging. X.-J.C., J-L.L., J.Y., and L.Zheng. assisted in brain slice imaging and electrophysiology. F.L. assisted in neuronal culture. M.Y.assisted in RNA seq., C.L.assisted in EM, X.Zou. and Z.F. assisted in mouse breeding, genotyping and immunostaining. Z.G. assisted in CSF collection. W.G., W.S., L.L., B.L., C.J., T.H., Z.L., L. Zhang, H. Zhang and H.Zeng. provided reagents. T.Z. and W.G. wrote the manuscript. All authors discussed, reviewed, and edited the manuscript.

## Supplementary Figure Legends

**Supplementary Fig. 1**

***Sspo* mRNA expression pattern in the brain of C57B6/J mice.**

Fluorescence images were acquired with RNAscope mFISH. The brightest green fluorescence was mainly located at the side facing the third ventricle. a’, b’, c’, insets. Scale bars, 50 µm (a, b, c); 10 µm (a’, b’, c’).

**Supplementary Fig. 2**

**Strategy for generating *Spdef-Cre* and *Car3-Cre* mice and the profile of EGFP expression in ependymal cells of *Spdef-Cre;Ai47* and *Car3-Cre;Ai47* mice.**

**a,** The Cre sequence was inserted before the first exon of *Spdef* or *Car3* but after ATG. **b,** Images of a brain section from a *Spdef-Cre;Ai47* mouse at P3. **c,** Images of a brain section from a *Car3-Cre;Ai47* mouse at P15. Green, cells expressing EGFP; light blue and blue, nuclei labeled by Hoechst 33342 (HO). b’, c’, insets. Scale bars, 1 mm (b, c); 100 µm (b’, c’).

**Supplementary Fig. 3**

**Images of ependymal cells and choroid-plexus cells of *Sspo-CreER;Ai47* mice.**

**a,** Image of lateral ventricle. Tamoxifen was administered at E10, and mice were sacrificed at P3. No EGFP signals were detected in ependymal cells or choroid-plexus cells of *Sspo-CreER;Ai47* mice. LV, lateral ventricle; CP, choroid plexus; dashed lines, the margin of left lateral ventricle. **b,** Image of the fourth ventricle (4V). **c,** Image of the third ventricle (3V). **d,** Image of central canal in the spinal cord. CC, central canal. **e,** Image of choroid plexus (CP). Green, EGFP fluorescence; red, ependymal cells stained with anti-Foxj1 (a) or anti-S100β (b, d), and CP cells stained with anti-β-catenin (e); light blue and blue, nuclei stained by Hoechst 33342 (HO); white arrowhead (a), the starting point of tamoxifen injection; red arrowhead, time points of mouse sacrifice. Scale bars, 50 µm (a, e); 100 µm (b, c); 25 µm (d).

**Supplementary Fig. 4**

**EGFP expression in ependyma and choroid plexus from the brain of *Sspo-Cre;Ai47* mice.**

**a,** Images of lateral ventricle (LV) from the brain of a *Sspo-Cre;Ai47* mouse at P5. Dashed lines, the margin of left LV. **b,** Images of the fourth ventricle (4V). **c,** Images of the third ventricle (3V) from a *Sspo-Cre;Ai47* mouse at P5. **d,** Image of central canal in the spinal cord from a *Sspo-Cre;Ai47* mouse at P5. CC, central canal. **e,** Images of choroid plexus (CP) from a *Sspo-Cre;Ai47* mouse at P5. **f,** Percentage of CP cells expressing EGFP fluorescence in *Sspo-Cre;Ai47* mice (n = 10,569 CP cells in total). Only a very small number of regions with EGFP signal were detected in CP cells of *Sspo-Cre;Ai47* mice. Green, EGFP fluorescence; red, ependymal cells stained with anti-Foxj1 (a, c) or anti-S100β (b, d), and CP cells stained with anti-β-catenin (e); light blue and blue, nuclei stained by Hoechst 33342 (HO). Scale bars, 50 µm (a, c, e); 100 µm (b); 25 µm (d).

**Supplementary Fig. 5**

**EGFP expression in the floor plate of the brain in *Sspo-Cre;Ai47* mice.**

**a–d,** Images of the sagittal brain sections of *Sspo-Cre;Ai47* mice at embryonic days 12 (a), 15 (b), 16 (c), and 18 (d) depict EGFP fluorescence in SCO and floor plate. Green, EGFP fluorescence; light blue and blue, nuclei stained with Hoechst 33342 (HO). Scale bars are 300 µm.

**Supplementary Fig. 6**

**Images of SCO from *Sspo-Cre;Ai47* mice at different developmental stages.**

**a–f,** Images of SCO in coronal brain sections of E12 (a), E16 (b), E19 (c), P0 (d), P5 (e) and P15 (f) *Sspo-Cre;Ai47* mice. **g–j,** Images of SCO in sagittal brain sections of E12 (g), E19 (h), P0 (i) and P5 (j) *Sspo-Cre;Ai47* mice. Green, EGFP fluorescence; arrowheads, the location of SCO; white, nuclei stained with Hoechst 33342 (HO). Scale bars, 300 µm (a, g); 50 µm (b–f); 100 µm (h–j).

**Supplementary Fig. 7**

**Morphology of individual SCO cells in the brain of *Sspo-CreER;Ai47* mice.**

Mice received tamoxifen from their lactating mothers (once, 75 µg/g body weight) from P0 and were analyzed at P5 and P21. With a low dosage of tamoxifen, SCO cells were sparsely labeled, and the morphology of individual SCO cells could be observed. Most EGFP cells were specialized ependymal cells that sent their processes to the third ventricle and contacted central spinal fluid. **a,** Images of different coronal sections of SCO at P5. **b,** Percentage of ependymal cells and hypendymal cells in SCO of *Sspo-CreER;Ai47* mice at P5.

Quantification was based on the coronal SCO images from the images obtained from a confocal microscope (94.95%, n = 198 EGFP^+^ cells in total). **c,** Images of different sagittal sections of SCO at P21. **d,** Percentage of ependymal cells and hypendymal cells in SCO of *Sspo-CreER;Ai47* mice at P21. Quantification was based on the sagittal SCO images obtained from a confocal microscope (98.54%, n = 1,236 EGFP^+^ cells in total). Green, EGFP fluorescence; light blue and blue, nuclei stained with Hoechst 33342 (HO). Scale bars, 50 µm.

**Supplementary Fig. 8**

**Body weight and brain MRI of C57BL/6, *Sspo-Cre* and *Sspo-CreER* mice.**

**a,** Body weight of C57BL/6 (WT, green), *Sspo-Cre* heterozygous (purple) and *Sspo-CreER* heterozygous mice (orange) at P21 (n = 10 mice per group). **b,** Representative MRI images of the brain from C57BL/6, *Sspo-Cre* heterozygous and *Sspo-CreER* heterozygous mice at P21. **c,** Representative images of brain sections of C57BL/6, *Sspo-Cre* heterozygous and *Sspo-CreER* heterozygous mice at P21. Light blue, nuclei stained with Hoechst 33342 (HO). Scale bar, 2 mm (b), 1 mm (c). Two-tailed unpaired *t-*test, error bars, s.e.m.

**Supplementary Fig. 9**

**Detection of SCO-spondin in the cerebral spinal fluid of *Sspo-Cre;DTA* and control mice. a,** Identification of a unique peptide matching SCO-spondin in the cerebral spinal fluid of control mice at P0–3, named VKPSYC(+57.02)SC(+57.02)LDLLTGK; vertical red line, retention time (RT) = 25.82 min. **b**, Identification of another unique peptide matching SCO-spondin in the cerebral spinal fluid of control mice at P0–3, named LDLLTGK (vertical red line, RT = 25.56 min). **c, d,** Detection of SCO-spondin in the cerebral spinal fluid of control and *Sspo-Cre;DTA* mice at P0–3. Ctrl_1– 4 represents four samples of mixed cerebral spinal fluid from control mice (20 pups), whereas *Sspo-Cre;DTA*_1–4 represents four samples of mixed cerebral spinal fluid from *Sspo-Cre;DTA* mice (20 pups; n = 4 samples per group).

Microtubule-associated protein 6 (MAP6) and apolipoprotein E (APOE) were used as internal controls. Two-tailed unpaired *t-*test, *\*\*\*p* < 0.001. Error bars, s.e.m.

**Supplementary Fig. 10**

**Transmission electron microscopy images of the central canal in the spinal cord of *Sspo-Cre;DTA* and control mice.**

**a, b,** Images showing the central canal in the spinal cord of control mice. **c, d,** Images showing the ependymal cells located in the aqueduct of *Sspo-Cre;DTA* mice. The red arrowhead indicates Reissner’s fiber.

**Supplementary Fig. 11**

**Images of lateral ventricle and choroid-plexus sections of *Sspo-Cre;DTA* and control mice.**

**a,** Images of coronal lateral ventricle sections in the brain of *Sspo-Cre;DTA* and control mice at P0. **b,** Image of coronal lateral ventricle sections in the brain of *Sspo-Cre;DTA* and control mice at P12. **c,** Number of ependymal cells per millimeter in the brain of *Sspo-Cre;DTA* and control mice at P0 (Ctrl, 348.89±62.26/mm; *Sspo-Cre;DTA*, 375.56±54.39/mm, n = 10 regions). **d,** Number of ependymal cells per millimeter in the brain of *Sspo-Cre;DTA* and control mice at P12 (Ctrl, 257.78±61.96/mm, *Sspo-Cre;DTA*, 233.33±42.95/mm, n = 10 regions). **e,** Image of choroid plexus in the brain of control and *Sspo-Cre;DTA* mice at P3.

Dashed lines (c, Ctrl), the margin of lateral ventricle; red, ependymal cells stained with anti-S100β; green, choroid-plexus cells stained with anti-β-catenin; light blue, nuclei stained with Hoechst 33342 (HO). a’–d’, insets. Scale bar, 250 µm (a, b), 25 µm (a-a’, b’, b-a’, b’), 50 µm (e). Two-tailed unpaired *t-*test, error bars, s.e.m.

**Supplementary Fig. 12**

**Morphology of ependymal cells in the aqueduct and central canal of *Sspo-Cre;DTA* and control mice.**

**a, b,** Transmission electron microscopy images displaying the morphology of ependymal cells from the aqueduct of control (a) and *Sspo-Cre;DTA* (b) mice. **c, d,** Images showing the morphology of ependymal cells from the central canal of control (c) and *Sspo-Cre;DTA* (d) mice.

**Supplementary Fig. 13**

**Images of interventricular foramina, cerebral aqueduct, and central canal sections in *Sspo-Cre;DTA* and control mice.**

**a,** Images of coronal interventricular foramina in the brain of control and *Sspo-Cre;DTA* mice at P0. 3V, the third ventricle; yellow arrowheads, the location of interventricular foramina. **b– e,** Images and width measurements of the coronal cerebral aqueduct in the brain of control and *Sspo-Cre;DTA* mice at P3, P6, and P12 (n = 9 mice per group). Yellow arrowheads, the location of cerebral aqueducts. **f,** Images of coronal central canal sections in the brain of control and *Sspo-Cre;DTA* mice at P0. **g,** Schematic indicating the location of central canal sections in (f). Light blue, nuclei stained with Hoechst 33342 (HO). a’– b’, insets. Scale bar, 250 µm (a), 100 µm (a’, b’), 50 µm (b, f). Two-tailed unpaired *t-*test, error bars, s.e.m.

**Supplementary Fig. 14**

**Images of brain sections of *Sspo-CreER;DTA* mice at postnatal stages.**

Tamoxifen was injected into *Sspo-CreER;DTA* and control mice at P3 (for 4 consecutive days) or P25 (for 5 consecutive days), and brains were collected at P37 and P70, respectively. **a–d,** Representative images of brain sections of *Sspo-CreER;DTA* and control mice at P37 and P70. Light blue, nuclei stained with Hoechst 33342 (HO). Scale bar, 1 mm (a–d). White arrowheads, the starting point of tamoxifen injection; red arrowheads, time points for mouse sacrifice. a’ and b’, insets.

**Supplementary Fig. 15**

**Images of brain sections of *Sspo-CreER;DTA* and control mice at P21.**

**a, b,** Images of coronal brain sections of control (a) and *Sspo-CreER;DTA* mice (b) at P21. Green, neurons stained with anti-NeuN; light blue and blue, nuclei stained with Hoechst 33342 (HO). **c,** Density of NeuN^+^ cells in the cerebral cortex of *Sspo-CreER;DTA* and control mice at P21 (Ctrl, 1928.00±123.37/mm^2^, *Sspo-CreER;DTA*, 1921.60±296.01/mm^2^; n = 5 regions per group). **d, e,** Images of coronal brain sections of control (d) and *Sspo-CreER;DTA* mice (e) at P21. Green, neurons stained with anti-MAP2; light blue and blue, nuclei stained with Hoechst 33342 (HO). Tamoxifen was injected at P0 (intragastrically, see methods). I–VI, six layers of the cerebral cortex; dashed lines, cortical margin; white arrowheads, the starting point of tamoxifen injection; red arrowheads, time points for mouse sacrifice. Scale bar, 200 µm (a, b), 100 µm (d, e). Two-tailed unpaired *t-*test, error bars, s.e.m.

**Supplementary Fig. 16**

**Morphology of neurons in the brain of *Sspo-CreER;DTA* and control mice at P21.** Neurons were sparsely labeled by AAV-CAG-EGFP in the brain of *Sspo-Cre;DTA* and control mice. AAV-CAG-EGFP was injected at P0, and the brains were collected and sectioned at P21. **a,** Morphology of sparsely labeled neurons in control mice at P21. **b,** Morphology of sparsely labeled neurons in *Sspo-Cre;DTA* mice at P21. Tamoxifen was injected at P0 (intragastrically, see methods). Green, neurons labeled with AAVs-CAG-EGFP; white arrowheads, the starting point of tamoxifen injection; red arrowheads, time points of mouse sacrifice. a’ and b’, insets. Scale bar, 100 µm (a, b).

**Supplementary Fig. 17**

**Images of sections of the cerebral cortex of *Sspo-Cre;DTA* and control mice.**

TUNEL staining was applied to label apoptotic neurons in the cerebral cortex of *Sspo-Cre;DTA* and control mice at P3 (a) and P8 (b). **a,** Images of coronal sections of brain from the cerebral cortex of *Sspo-Cre;DTA* and control mice at P3. **b,** Images of coronal sections of brain from the cerebral cortex of *Sspo-Cre;DTA* and control mice at P8. Yellow arrowheads, TUNEL^+^NeuN^+^ cells; green, cells positive for TUNEL staining; red, neurons stained with anti-NeuN; light blue and blue, nuclei stained with Hoechst 33342 (HO); I–V, five layers of the cerebral cortex. **c,** Density of TUNEL^+^NeuN^+^ cells in the cerebral cortex of *Sspo-Cre;DTA* and control mice at P3 and P8 (P3, Ctrl, 15.20±9.26/mm^2^, *Sspo-Cre;DTA*, 236.00±32.50/mm^2^; P8, Ctrl, 3.20±2.99/mm^2^, *Sspo-Cre;DTA*, 10.40±4.08/mm^2^, n = 5 regions per group). Light blue, nuclei stained with Hoechst 33342 (HO). Scale bar, 200 µm (a, b). \**p < 0.05*, *\*\*\*p < 0.001.* Error bars, s.e.m.

**Supplementary Fig. 18**

**Images of brain sections of *Sspo-Cre;DTA* and control mice at E18.**

**a,** Representative images of coronal brain sections of a control mouse at E18.

**b,** Representative images of coronal brain sections of a *Sspo-Cre;DTA* mouse at E18, with a thinner cortex than that of the control mouse. Green, neurons stained with anti-CTIP2; light blue and blue, nuclei stained with Hoechst 33342 (HO); I–VI, six layers of the cerebral cortex; dashed lines, cortex margin. a’–d’, insets. Scale bar, 200 µm (a, b), 50 µm (a’, b’). **c,** Cortical thickness of *Sspo-Cre;DTA* and control mice at E18 (Ctrl, 0.29±0.06 mm, *Sspo-Cre;DTA*, 0.20±0.06 mm, n = 5 regions per group). **d,** Percentage of CTIP2^+^ cells among total cells of whole cerebral cortex from *Sspo-Cre;DTA* and control mice at E18 (Ctrl, 40.46±1.68%, *Sspo-Cre;DTA*, 27.38±3.35%, n = 5 regions per group). Two-tailed unpaired *t-*test, *\*p < 0.05*, *\*\*\*p < 0.001*. Error bars, s.e.m.

**Supplementary Fig. 19**

**Images of sections from the cerebral cortex of *Sspo-Cre;DTA* and control mice at P22.**

**a,** Images of coronal sections from the cerebral cortex of a *Sspo-Cre;DTA* and a control mouse at P22. Dashed lines, the margin of TBR1^+^ cell distribution; green, cells stained with anti-TBR1; blue, nuclei stained with Hoechst 33342 (HO). **b,** Density of TBR1^+^ cells in the cerebral cortex of *Sspo-Cre;DTA* and control mice at P22 (Ctrl, 1902.00±145.62/mm^2^, *Sspo-Cre;DTA*, 1172.00±47.92mm^2^, n = 4 regions per group). Scale bar, 100 µm (a). Two-tailed unpaired *t-*test, *\*\*p < 0.01.* Error bars, s.e.m.

**Supplementary Fig. 20**

**Representative images of the corpus callosum from *Sspo-Cre;DTA* and control mice at P3.**

Images of brain sections with the corpus callosum (white matter) in a control mouse (left) and *Sspo-Cre;DTA* mouse (right). Green, neurons stained with anti-NeuN; blue, nuclei stained with Hoechst 33342 (HO). Arrowheads, dislocated neurons observed in the corpus callosum. WM, white matter; LV, lateral ventricle. a’, insets. Scale bar, 100 µm.

**Supplementary Fig. 21**

**Pathway enrichment visualization.**

Top 25 Gene Ontology pathways from a Gene Set Enrichment Analysis of the SCO and hippocampus (adjusted *p*-values < 0.05). The dot plot depicts the activity of the enriched signaling pathways (activated) and suppressed pathways in SCO. The color represents the adjusted *p*-values (brighter red represents more significant), and the size of the terms represents the number of genes for which expression differed significantly (larger size represents greater expression).

**Supplementary Fig. 22**

**Sequences measured with mass spectrometry.**

Sequences of Tβ4 (a), Tβ10 (b), and NP24 (c) detected in the cerebral spinal fluid of C57BL/6 mice at E18–P0.

**Supplementary Fig. 23**

**Density and morphology of neurons in the absence or presence of Tβ4, Tβ10, and NP24. a,** Representative images (bright field) of neurons cultured with Tβ4, Tβ10, and NP24 at DIV12. **b,** Summary of data for neuron morphology (terminal points of apical dendrites) cultured in the presence of Tβ4, Tβ10, and NP24 (supplied with 0.4% B27; n = 50 neurons per group). Scale bars, 50 µm (a). Two-tailed unpaired *t-*test, *\*\*\*p < 0.001.* Error bars, s.e.m.

**Supplementary Fig. 24**

**RNA-seq of cultured neurons incubated with individual peptides Tβ4, Tβ10, or NP24.** Cultured neurons were collected for RNA-seq at DIV4 after they were incubated for 2 days with individual peptides Tβ4, Tβ10, or NP24 at DIV2. Gene Set Enrichment Analysis showed that the pathway ‘Positive regulation of neuron projection development’ was active in neurons incubated with Tβ10, with a higher mRNA abundance of the genes *Ngf* (Ctrl, 28.61±16.62; neurons incubated with Tβ10, 60.78±4.62; n = 4 samples per group) and *Ptn* (Ctrl, 4915.56±362.20; neurons incubated with Tβ10, 6821.95±547.73; n = 4 samples per group).

The pathway ‘Positive regulation of cell population proliferation’ was active in neurons incubated with Tβ4, with a higher mRNA abundance of the genes *Lin28a* (Ctrl, 0.53±0.54; neurons incubated with Tβ4, 9.06±13.20; n = 4 samples per group) and *Notch3* (Ctrl, 266.11±46.25; neurons incubated with Tβ4, 392.80±68.72; n = 4 samples per group). The pathway ‘Positive regulation of cell death’ was inactive in neurons incubated with NP24, with a lower mRNA abundance of the genes *Casp9* (Ctrl, 323.74±17.49; neurons incubated with NP24, 244.19±60.58; n = 4 samples per group) and *Atf4* (Ctrl, 1637.93±265.16; neurons incubated with NP24, 1143.21±89.32; n = 4 samples per group). Two-tailed unpaired *t-*test, *\*p < 0.05, **p < 0.01.* Error bars, s.e.m.

## Methods

### Mice

The use and care of animals followed the guidelines of the Institutional Animal Care and Use Committee at Chinese Institute for Brain Research, Beijing. *DTA* (Stock No. 009669) and *Ai14* (Stock No. 007914) mice were purchased from the Jackson Laboratory. *Sspo-Cre, Sspo-CreER, Car3-Cre,* and *Spdef-Cre* mouse lines (C57BL/6 background) were developed by our laboratory with the assistance from Biocytogen Pharmaceuticals (Beijing). The C57BL/6 control mice were purchased from Charles River Laboratory. *Ai47* mice were originally from Dr. Hongkui Zeng’s lab at Allen Institute.

### Brain excision and preparation of thin sections

Mice (**≥**P10) were anesthetized with isoflurane using a vaporizer and were perfused with phosphate-buffered saline (PBS) at room temperature followed by cold 4% paraformaldehyde. Mice younger than P10 were euthanized by hypothermia (P0–7) before decapitation or were euthanized by decapitation directly (P0–9). Each brain was excised and postfixed in 4% paraformaldehyde at 4°C overnight and then dehydrated in 30% sucrose at 4°C for 2–3 days. After embedding in OCT compound (Tissue-Tek4583, Sakura), each frozen brain was sectioned with a cryostat (Leica, CM3050S) at a thickness of 30 μm for immunostaining or 15 μm for RNAscope mFISH.

### SCO isolation

Male adult mice (8–10 weeks old) were euthanized with isoflurane. Each brain was resected and placed in cold PBS (4°C). Neonatal pups (P0–7) were anesthetized on ice and decapitated for brain dissection. All brains were placed in cold PBS. Hippocampus and SCO were isolated under a stereoscope (Leica M205 FA). All samples for bulk RNA-seq or peptidomic analysis were stored at –80°C.

### Bulk RNA-seq and data analysis

Total RNA was extracted from the SCO and hippocampus or cultured neurons using the RNeasy Micro kit (Qiagen, 74004). The DNA library for sequencing was prepared following the protocol for TruePrep DNA Library Prep kit V2 (SCO and hippocampus) or SMARTer Stranded Total RNA-Seq kit v2 (cultured neurons). RNA-seq raw data were initially filtered with “FastQC” to obtain clean data after quality control. Clean data were aligned to the reference genome (Version: GRCm38) downloaded from Gencode by HISAT. Raw counts for each gene were calculated by FeatureCounts. The expression level of detected genes was estimated by DESeq2. A volcano plot was generated by ggplot2 in R, with a cutoff *p*-value of <0.01 and absolute value of log_2_ (Foldchange) of >4, measured by DESeq2. Gene Set Enrichment Analysis was performed using the ClusterProfiler package in R, with an adjusted *p*-value of <0.05.

### RNAscope mFISH

The RNAscope mFISH assay was performed with the RNAscope multiplex fluorescent reagent kit v2 (Cat. No. 323100, Advanced Cell Diagnostics) and HybEZ II hybridization system (Advanced Cell Diagnostics). Advanced Cell Diagnostics designed the RNAscope probes for *Car3* (578571-C3, target region: positions 2–1002 of RNA complementary to the sequence NM_007606.3), *Spdef* (544421-C2, target region: positions 471–1400 of RNA complementary to the sequence NM_013891.4), and *Sspo* (1134541-C1, target region: positions 344–1232 of RNA complementary to the sequence NM_173428.4). The positive control (Ms-Ppib, 313911, Advanced Cell Diagnostics) and negative control (DapB, 310043, Advanced Cell Diagnostics) were applied to evaluate the RNA quality of processed brain sections.

### Drug administration

For all CreER mice, tamoxifen (10 mg/ml, dissolved in a 1:9 mixture of ethanol and sunflower oil) was intraperitoneally injected into mice on or after P5 at 75 µg/g body weight for 5 consecutive days or was injected intragastrically into neonatal mice (P0–4) at a dosage of 50 µg (i.e., 5 µl) per mouse daily for 4 consecutive days. For induction of embryonic mice, tamoxifen (100 mg/ml) was administered to pregnant mice by oral gavage at E10 with a dosage of 5 mg (i.e., 50 µl) once every other day for 4 consecutive days.

### EdU staining

To label dividing cells in *Sspo-Cre;DTA* mice at E15, EdU (2 mg/ml, dissolved in 0.9% saline, Beyotime, ST067) was intraperitoneally injected into their mothers at a dosage of 3 μl/g body weight per mouse. The brain of each *Sspo-Cre;DTA* mouse at E18 was excised and sectioned (30 μm) after fixation with PFA. Brain sections were permeabilized with 0.5% (w/v) Triton X-100 for 30 min and incubated with EdU staining cocktail for 30 min at room temperature (cocktail: Tris-buffered saline (Solarbio, T1080), CuSO_4_ (4 mM, Sigma-Aldrich, C8027), sodium ascorbate (20 mg/ml, Sigma-Aldrich, A7631), and sulfa-cyanine3 azide (3 μM, Lumiprobe, B1330). Sections were then washed in PBS before being mounted with mounting medium (Electron Microscopy Sciences, 17985).

### TUNEL staining

TUNEL (terminal deoxynucleotidyl transferase dUTP nick end labeling) staining was performed after immunostaining using the One Step TUNEL Apoptosis Assay kit (producing red fluorescence; Beyotime, C1090). Terminal deoxynucleotide transferase (TdT) was diluted with an equal volume of TdT diluent (Beyotime). A TUNEL staining cocktail of diluted TdT (10%) and fluorescent labeling solution (90%) was prepared. Brain sections were incubated with the TUNEL staining cocktail for 1 h at room temperature, washed in PBS, and mounted with mounting medium.

### Scanning transmission electron microscopy

Mice were perfused with PBS followed by 4% paraformaldehyde. Isolated tissues were post-fixed with 2.5% glutaraldehyde. Tissue blocks containing the aqueduct and spinal central canal were then isolated and post-fixed with 1% osmium tetroxide. The samples were dehydrated through a graded series of ethanol concentrations (50–100%) and infiltrated before embedding in SPI-PON resin (90529-77-4). Araldite resin blocks were sectioned at a thickness of 1.5 µm and stained with 1% toluidine blue for light microscopy assessment. Ultrathin sections (70 nm) were obtained from selected regions and stained with uranyl acetate and lead citrate. Specimens were imaged with a Hitachi HT7800 transmission electron microscope.

### MRI and 3D reconstruction

MRI was performed with a 7.0T scanner (Pharmascan 70/16, Bruker; Germany) equipped with a 23-mm surface coil and a 12-cm diameter self-shielded gradient system. The user interface consisted of Paravision 5.1 software (Bruker BioSpin) and a Linux PC running Topspin 2.0. Mice were initially anesthetized with 5% isoflurane and 95% O_2_ and then maintained with 2% isoflurane and 98% O_2_ during imaging. T2-weighted imaging was performed on each brain at different time points with a method of relaxation enhancement using the following parameters: repetition time/echo time 3500 ms/33 ms, relaxation enhancement factor = 4, 21×21mm field of view, 256×256 matrix, 20–25 slices, and 0.5-mm slice thickness. A 3D slicer software was used for the 3D reconstruction of ventricles and for quantifying ventricle volume.

### Virus preparation, purification, and infection

To sparsely label neurons in brain, plasmid (*p*)CAG-EGFP was designed to express EGFP under the control of CAG promoter. Each AAV-CAG-EGFP was packaged together with helper plasmids (phelper) and *p*RC. To label cultured neurons, *p*FUGWH1-EGFP was applied, and lentivirus (LV)-H1-EGFP was packaged together with *p*sPAX2 (Cat. No. 12260, Addgene) and *p*MD2.G (Cat. No. 12259, Addgene). HEK 293T cells were transfected with the above three-plasmid system for AAVs or LVs using Neofect (TF201201). After purification, the AAVs (10^12^ gc/ml) and LVs (10^8^–10^9^ TU/ml) were stored at –80°C for further analysis.

### *In vivo* neuronal labeling and 3D reconstruction

AAV-CAG-EGFP were injected into the lateral ventricle of embryonic mice at a concentration of 10^9^–10^10^ gc/ml. Each brain was harvested at least 7 days after viral injection. Brain sections with EGFP-expressing neurons were imaged with a VS120 Virtual Slide Microscope (Olympus) and confocal microscope (Leica, SP8). Neuronal morphology was analyzed and reconstructed with Imaris ×64 9.7.1 software (Oxford Instruments). Imaris was used to analyze parameters regarding apical dendrites, including the length of the apical dendrites, the number of dendritic branch points, and the number of terminal points.

### Morphological analysis of neurons *in vitro*

Neurons labeled with anti-Tuj1 were imaged by a confocal microscope (Leica, SP8). To quantify neurite density, we first plotted circles along the neuronal soma, 20 μm and 40 μm away from the soma surface and then counted the numbers of intersections between neuronal processes and these circles. Finally, the reconstruction of neuronal morphology and the analysis of dendrites were performed as for EGFP-expressing neurons *in vivo*.

### Co-culture of neurons with SCO and *in vitro* neuron labeling with lentiviruses

Primary cortical neurons, SCO and hippocampus were isolated from P0–1 *Sspo-Cre;Ai47* pups. Neurons were cultured on coverslips coated with poly-D-lysine in 24-well plates at a density of 5×10^4^ cells per well. The culture medium contained Neurobasal-A (Thermo Fisher, 10888022) supplemented with B27 (2%, Thermo Fisher, 17504044) and GlutaMAX (2 mM, Thermo Fisher, 35050061) with penicillin and streptomycin. Transwells (0.4 µm pore membrane, Corning, 3470) were hung on the top of the coverslip, and an excised SCO from a *Sspo-Cre;Ai47* pup was placed on the pore membrane. Cultured neurons were infected with LVs-H1-EGFP at least 3 days for labeling to assess morphology. For immunostaining, cultured neurons were washed with PBS three times and fixed with 4% paraformaldehyde for 15 min.

### Proteomic data analysis

Data-independent acquisition processing was carried out with PEAKS against a refined non-redundant mouse UniProt database with 22,001 entries. Searches utilized a 10-ppm tolerance for peptide masses and 0.02 Da for HCD (higher-energy C-trap dissociation) fragment ion masses. Variable modification included oxidation (methionine), and carbamidomethylation (cysteine) was configured as a fixed modification. The PEAKS quantification module was used to conduct label-free quantitation based on extracted ion chromatograms. Quantitative data were normalized to the summed intensities of all peptide matches.

### Digestion of proteins isolated from mouse cerebrospinal fluid

CSF samples were diluted with 8 M urea. Each resulting denatured protein sample was chemically reduced with 100 mM dithiothreitol (Roche) at 56°C for 30 min and alkylated using 100 mM iodoacetamide for 1 h. Proteins were digested with trypsin using the Filter-Aided Sample Preparation technique^48^. Specifically, 10 µl of each protein mixture was loaded onto a 10 kDa molecular-weight cutoff filter unit that was then centrifuged (14,500 *g*, 15 min). Proteins in the recovered solution were then reduced with dithiothreitol and alkylated with iodoacetamide as noted above. The filter membrane was subsequently washed three times with 50 mM NH_4_HCO_3_ to eliminate any remaining urea, and proteins were digested with 1 μg trypsin per membrane overnight at 37°C. The resulting tryptic peptides were eluted from the membrane by centrifugation (14,500 *g*, 15 min). An additional 50 μl of 50 mM NH_4_HCO_3_ was added to the filter membrane, and the membrane was centrifuged again (14,500 *g*, 15 min). Finally, the two eluates were combined for subsequent analysis.

### Proteomics analysis via LC-MS

Samples of tryptic peptides were spiked with the iRT internal standard peptide mix (Biognosys, Switzerland) and analyzed using a nanoLC system (M class, Waters) coupled to an Orbitrap Exploris 480 Mass Spectrometer (Thermo Fisher Scientific). Each sample was loaded onto an Acclaim PepMap trap column (75 µm × 2 cm, 3 µm, C18, 100 Å, Thermo Scientific) and then separated on a nanoEase BEH C18 column (150 µm × 100 mm, 1.7 µm, C18, 130 Å, Waters) with solvent A (0.1% formic acid in water) and solvent B (0.1% formic acid in 100% acetonitrile) maintaining a constant flow rate of 600 nl/min. Peptides were eluted with the following gradient: from 1 to 5% solvent B over 1 min, from 5 to 10% solvent B over 2 min, from 10 to 30% solvent B over 43 min, from 30 to 95% solvent B over 2 min, followed by a 4-min retention at 95% solvent B. This was succeeded by a 1-min transition to 1% solvent B, which was then maintained for 7 min prior to analysis of the next sample.

The AGC (automatic gain control) target value for the full MS1 scan was set at 300%, with a maximum injection time of 100 ms. For the MS2 scan, the AGC target value was consistently established at 1000%, paired with a resolution of 30,000 and an injection time of 90 ms. Furthermore, the entire MS scan range was dynamically divided into 60 MS/MS isolation windows, each with a 1-Da overlap, and the collision energy was normalized to 30.

### Extraction of endogenous peptides from mouse brain

Individual SCO samples from 120 mice and control hippocampal samples were dissolved in water/methanol/acetic acid (1:90:9, v/v/v) (10 µl per mg tissue) After homogenizing with a Sonic Dismembrator (8 s on, 15 s off, three cycles in total, Fisher Scientific Model FB120), the mixture was centrifuged at 20,000 *g* (20 min, 4°C). The clarified supernatant (200 μl) was aliquoted and loaded onto a 30 kDa molecular-weight cutoff filter (Millipore Amicon Ultra, Burlington, MA) that had undergone pre-rinsing as follows: (i) 0.1 M sodium hydroxide, 200 μl (ii) water/methanol/acetonitrile solution (50:30:20, v/v/v), 200 μl, repeated one time, and (iii) water/methanol/acetic acid (1:90:9, v/v/v), 200 μl. For each pre-rinse, the filter was centrifuged at 15,000 *g* (5 min, 4°C). The supernatant-loaded filters were centrifuged at 15,000 *g* (30 min, 4°C), and the filtrates were collected as the neuropeptide samples (<30 kDa). The flow-through was combined with the previously collected peptide samples (total volume ∼600 μl). All samples were then aliquoted and lyophilized. C18 Zip tips (Millipore Sigma, St. Louis, MO) were used for desalting. After lyophilization, the peptide samples were stored at –80°C until LC-MS analysis.

### Extraction of peptides from mouse CSF

CSF was collected from 20–30 embryos or neonatal pups of C57BL6 or *Sspo-Cre;DTA* mice and transferred into two different tubes, with a 55 µl final volume for each sample for subsequent analysis. Each sample was combined with 600 µl water/methanol/acetic acid (1:90:9, v/v/v) and kept on ice for 15 min, then centrifuged at 20,000 *g* for 10 min. Each supernatant was dried in a SpeedVac vacuum concentrator. Each peptide sample was then resuspended in 100 µl water containing 0.1% formic acid, followed by desalting with C18 tips.

### LC-Orbitrap MS peptidomics

The peptides extracted from the SCO or control samples were analyzed using a nanoLC system (Dionex UltiMate 3000, Thermo Fisher Scientific) coupled to an Orbitrap Fusion Lumos mass spectrometer (Thermo Fisher Scientific). The samples were reconstituted in a loading solvent of 3% acetonitrile and 0.1% formic acid (v/v). The analytical column was self-made with an integrated emitter tip and dimensions of 75 μm inner diameter × 17 cm length, 3.0 μm particle size, packed with 1.7 μm, 150 Å, BEH C18 material (Waters, Milford, MA). Solvent A was water containing 0.1% formic acid, and solvent B was acetonitrile containing 0.1% formic acid. The flow rate was 0.3 μl/min, and the gradient for peptide elution was as follows: 0–16 min, 3% solvent B; 16–20 min, 3–25% B; 20–30 min, 25–45% B; 30–50 min, 45–70% B; 50–56 min, 70–95% B; 56–60 min 95% B; 60–60.5 min, 95–3% B; 60.5–70 min, 3% B. Data were acquired in top-speed data-dependent mode. Other parameters included: precursor scan AGC, 1×106; MS/MS scan AGC, 5×104; isolation window, 1 m/z; normalized collision energy, 30%; with both HCD and electron-transfer/higher-energy collision dissociation MS/MS analysis.

Peptides extracted from CSF were analyzed with an Orbitrap Q-Exactive HF mass spectrometer (Thermo Scientific) coupled with an online Easy-nLC 1200 nano-HPLC system (Thermo Scientific). Briefly, 5 µl of peptide mixture was loaded on a reversed phase precolumn and then separated on a reversed phase analytical column at a flow rate of 600 nl/min with a 65-min gradient as reported previously^49^. Peptides were analyzed in data-dependent MS/MS acquisition mode and fragmented by HCD for MS/MS analysis. Detailed parameters for instrument settings can been found in our previous report^49^.

### Peptidomic data analysis

Raw data of peptides extracted from the SCO, CSF, and corresponding control samples were searched against the mouse neuropeptide database combining Neuropep and SwePep (337 entries) with PEAKS 8.5 for neuropeptide identification. A mass tolerance of ±10 ppm was used for precursors; monoisotopic mass tolerance was set to ±0.02 Da for product ions. HCD fragmentation type and electron-transfer/higher-energy collision dissociation fragmentation were selected individually. An advanced setting that searched against 313 build-in modifications was used for more accurate searching of post-translational modifications. Parameters for confident neuropeptide identification were Ascore (post-translational modification site confidence) higher than 20, false-discovery rate lower than 1%, and the presence of at least one unique peptide. Besides the mature neuropeptide database, a custom-built candidate preprohormone database was also used to discover novel neuropeptides, as reported previously^50^.

Raw data for peptides extracted from cerebrospinal fluid were processed with PEAKS Studio v8.5 against the neuropeptide database combining Neuropeptides.nl, SwePep, Neuropred.com, and Neuropedia.com (487 entries) or UniProt database (downloaded December 9, 2021, encompassing 17,547 entries). Mass tolerance for searches was set to 10 ppm for peptide masses and 0.02 Da for HCD fragment ion masses. Enzyme was set to none. The pyroglutamylation (N-terminal glutamine), oxidation (methionine), amidation (C-terminus), phosphorylation (serine or threonine) and acetylation (lysine) were set as variable modifications. Label-free quantitation based on extracted ion chromatograms was performed using the PEAKS Q module, and quantitative information was exported as .cvs files for further bioinformatics processing.

### Peptide-based rescue *in vitro* and *in vivo*

For the *in vitro* rescue assay, synthetic Tβ4, Tβ10, or NP24 (500 ng/µl, dissolved in 0.9% saline) was mixed with cultured neurons every 3 days from DIV0, and treated neurons were collected between DIV11–12 for staining. For the *in vivo* rescue assay, a cocktail containing Tβ4, Tβ10, and NP24 (1 µg/µl for each, dissolved in 0.9% saline) or single Tβ4, Tβ10, or NP24 (0.5 µg/µl or 3 µg/µl for each peptide], dissolved in 0.9% saline) was injected into the lateral ventricle of *Sspo-Cre;DTA* mice from P0 for three constitutive days. The control groups included non-injected *Sspo-Cre;DTA* mice and cocktail-injected wild-type mice from the same litter. The body weight and overall survival rate were then calculated. MRI was performed, and each brain was sectioned to measure the volume of lateral ventricles. AAV-CAG-GFP was applied for neuronal labeling for morphological analysis in the brain of *Sspo-Cre;DTA* and control mice treated with SCO peptides.

### Statistics and reproducibility

All quantitative data were analyzed using GraphPad Prism (9.0). The two-tailed unpaired Student’s *t*-test was used to evaluate the statistical significance of differences between two groups, and the mean ± s.e.m. are presented with the individual data points shown simultaneously. Statistical significance was analyzed by p-value: *p < 0.05, **p < 0.01, ***p < 0.001.

