## Supplementary figures and images for "The subcommissural organ regulates brain development via secreted peptides"

### Supplementary Fig. 1

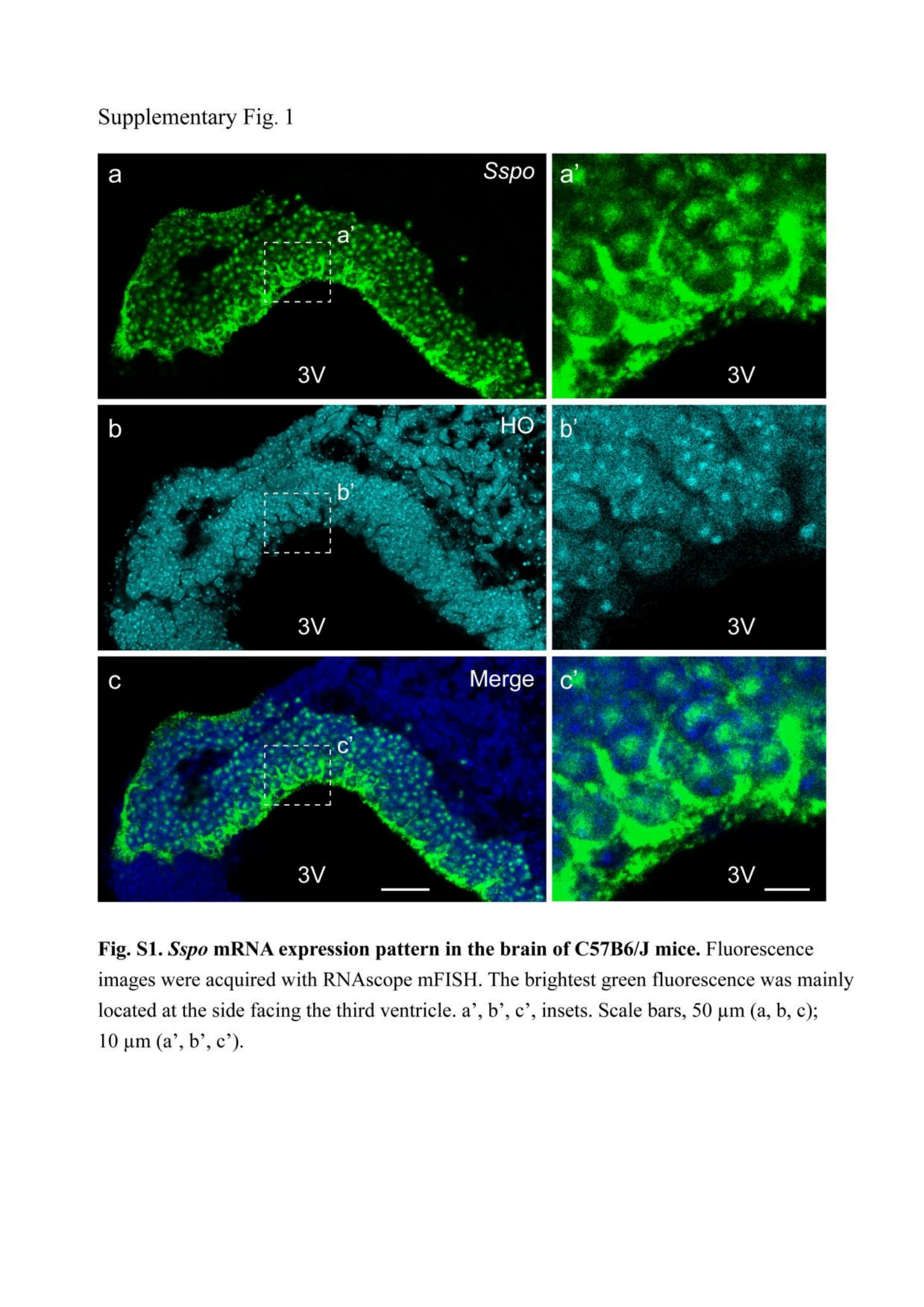

### Supplementary Fig. 2

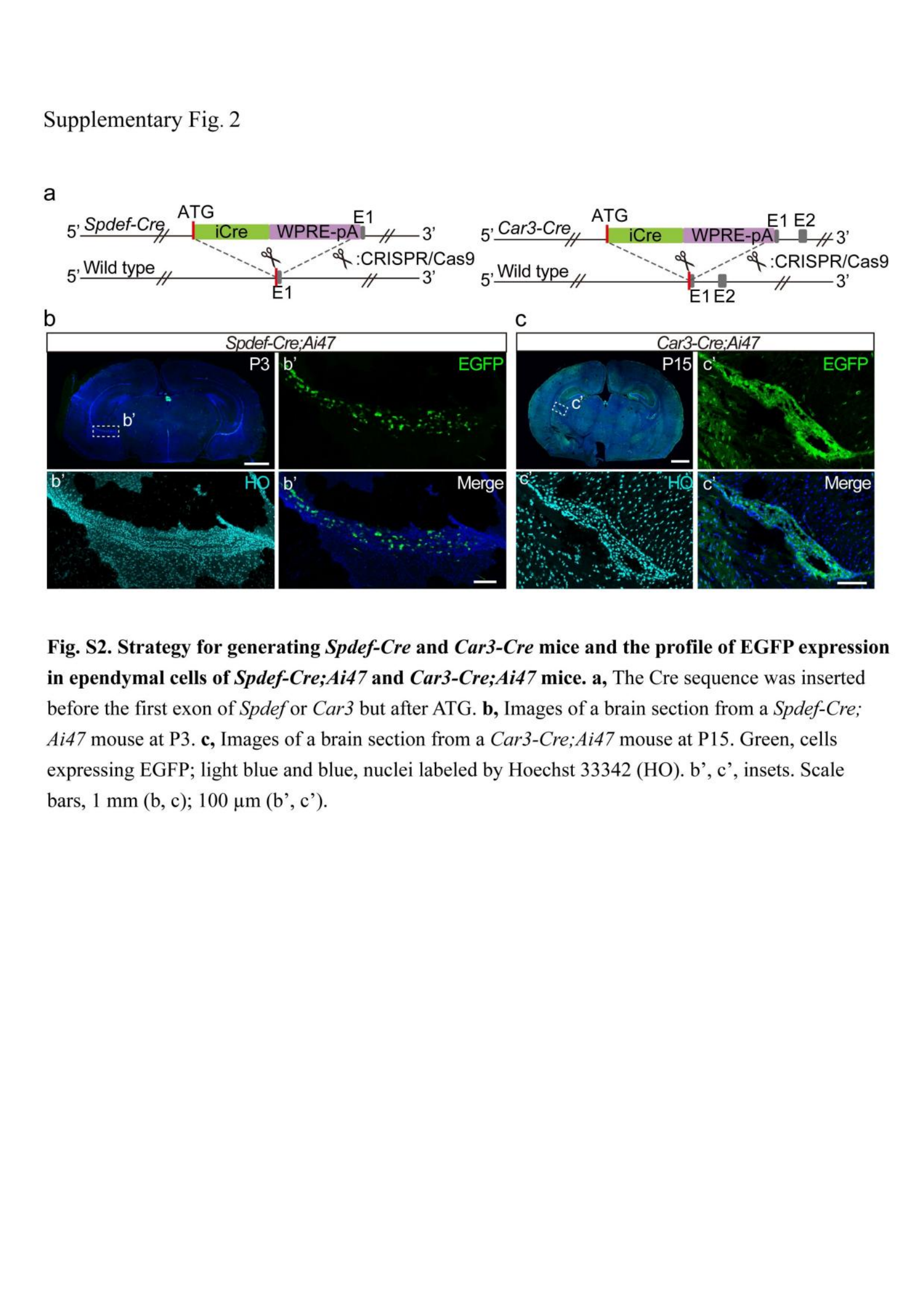

### Supplementary Fig. 3

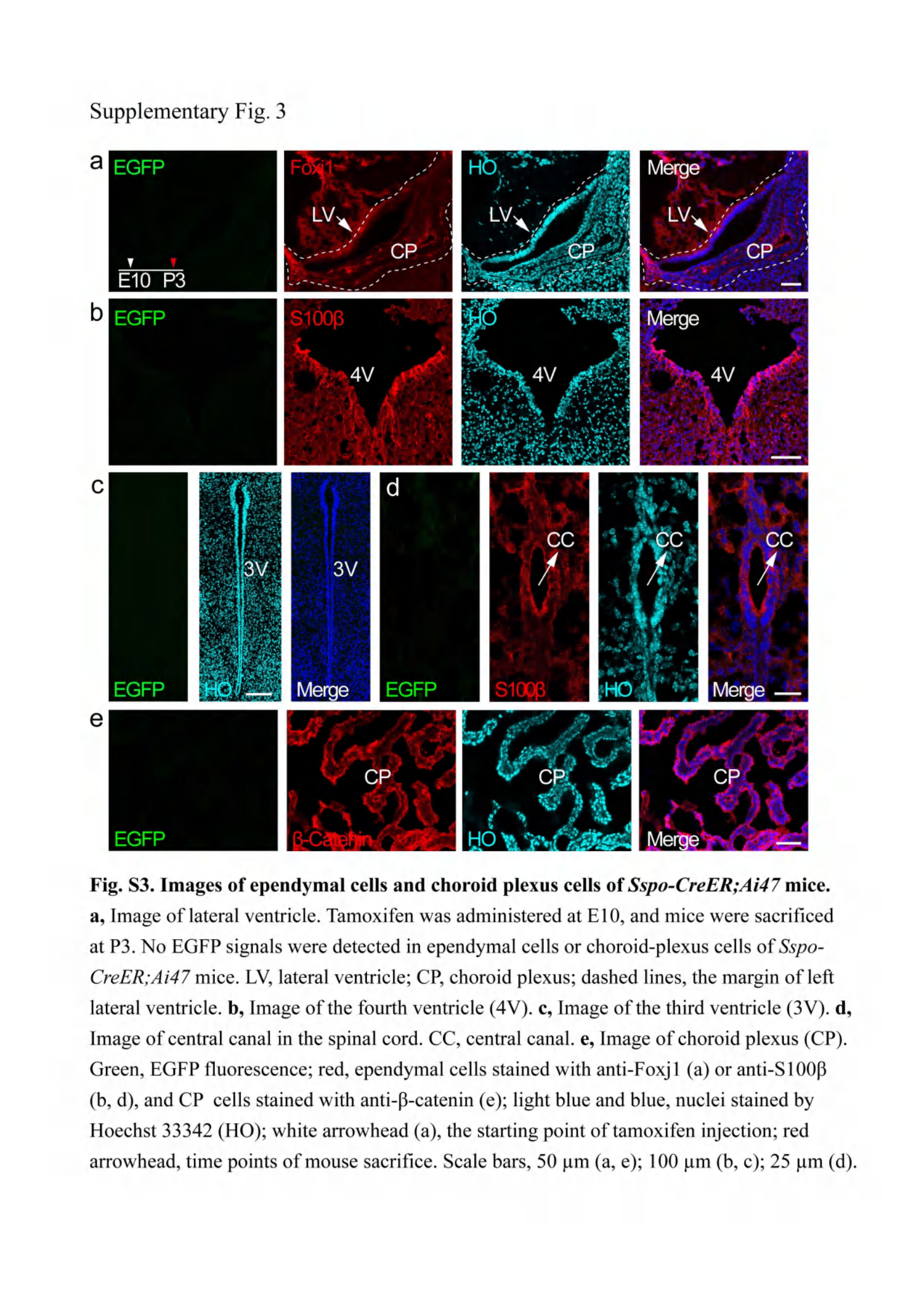

### Supplementary Fig. 4

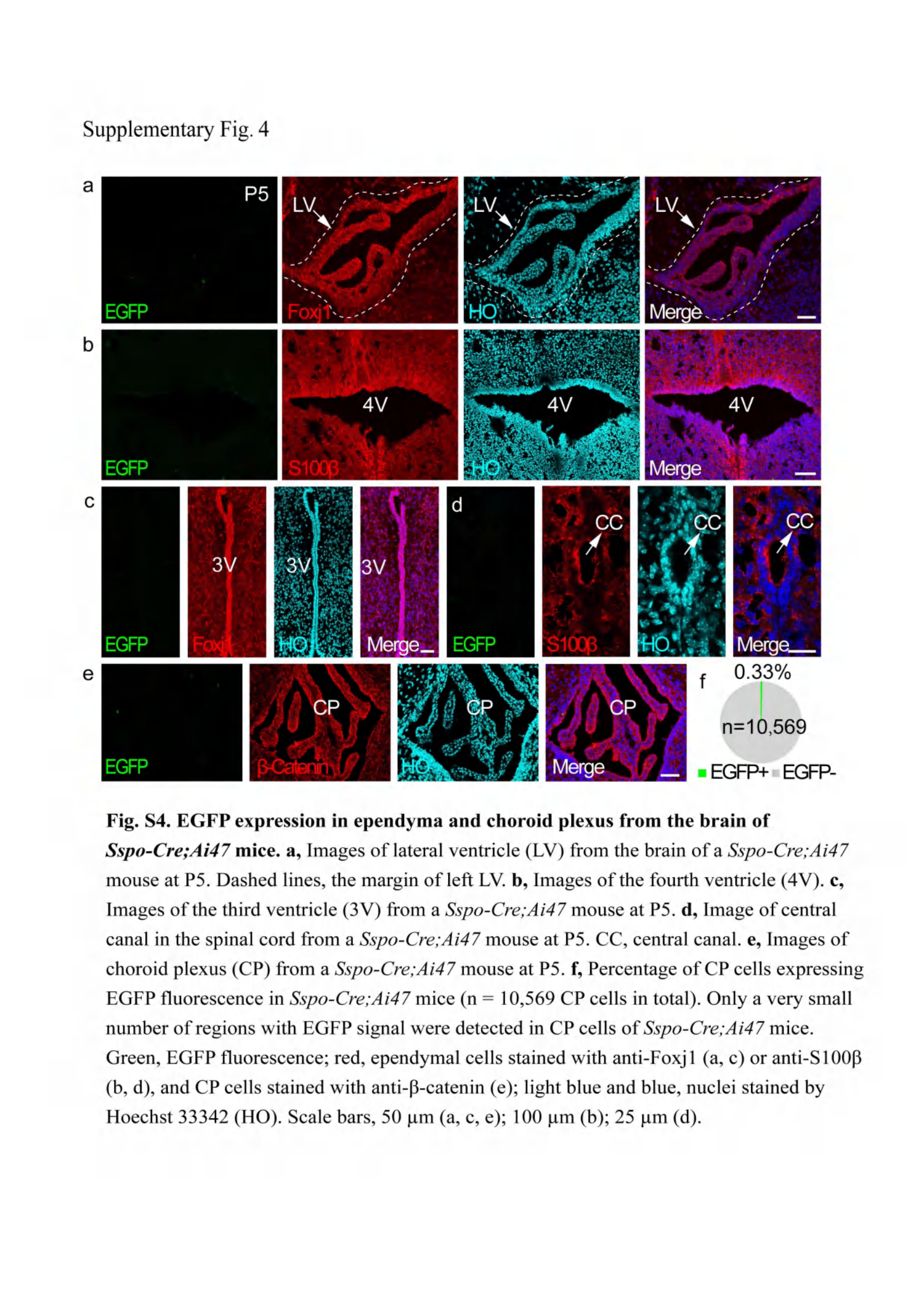

### Supplementary Fig. 5

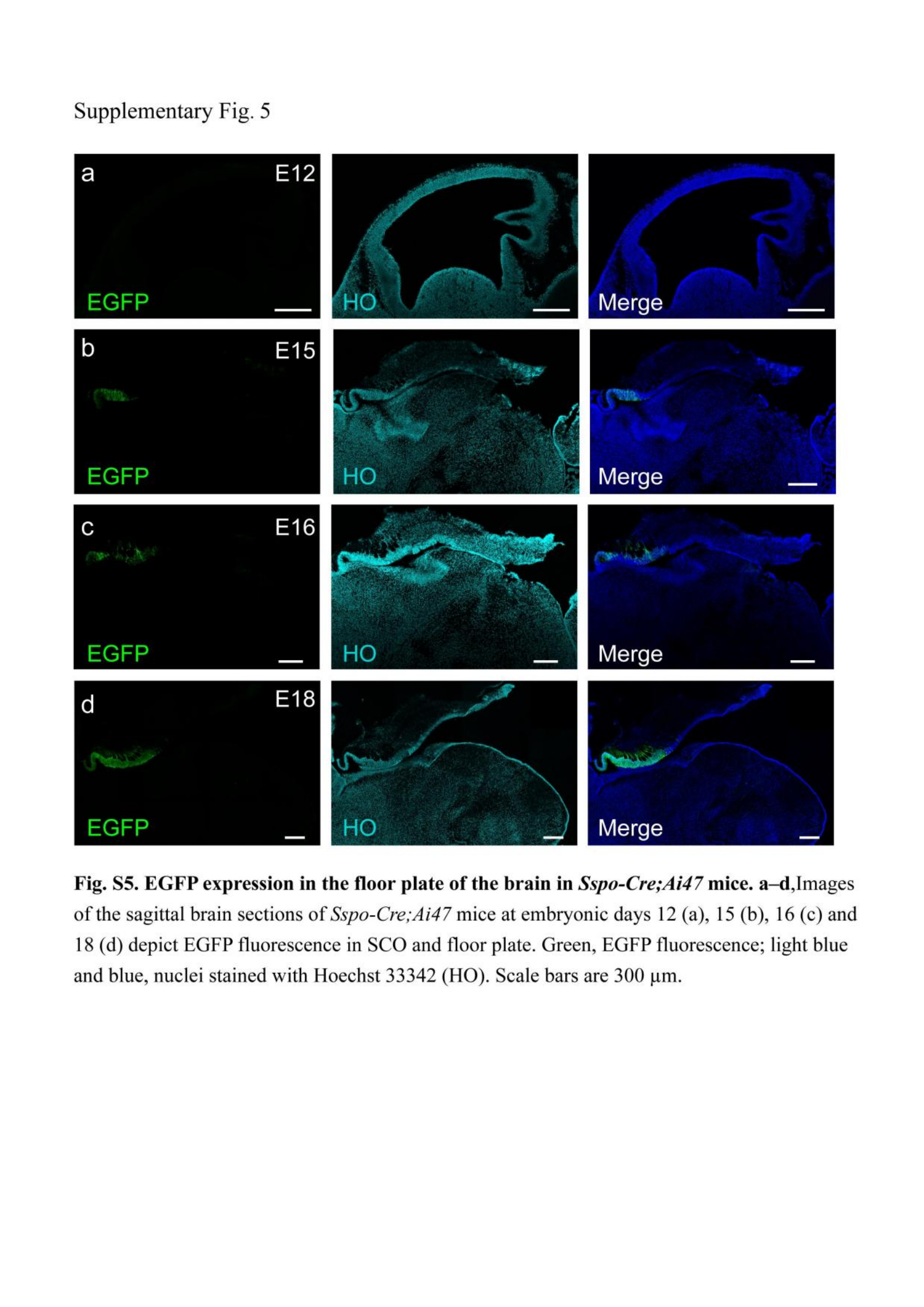

### Supplementary Fig. 6

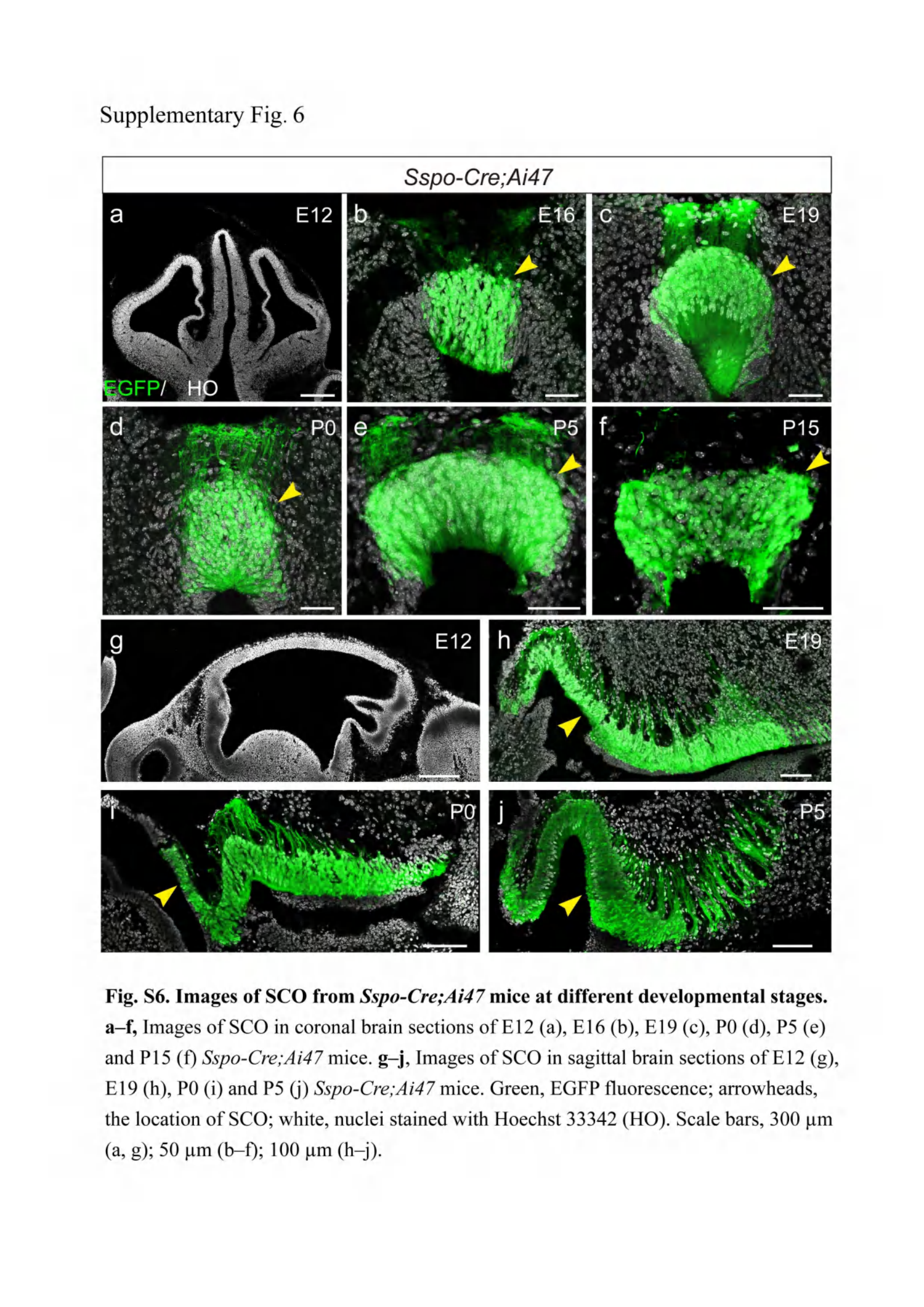

### Supplementary Fig. 7

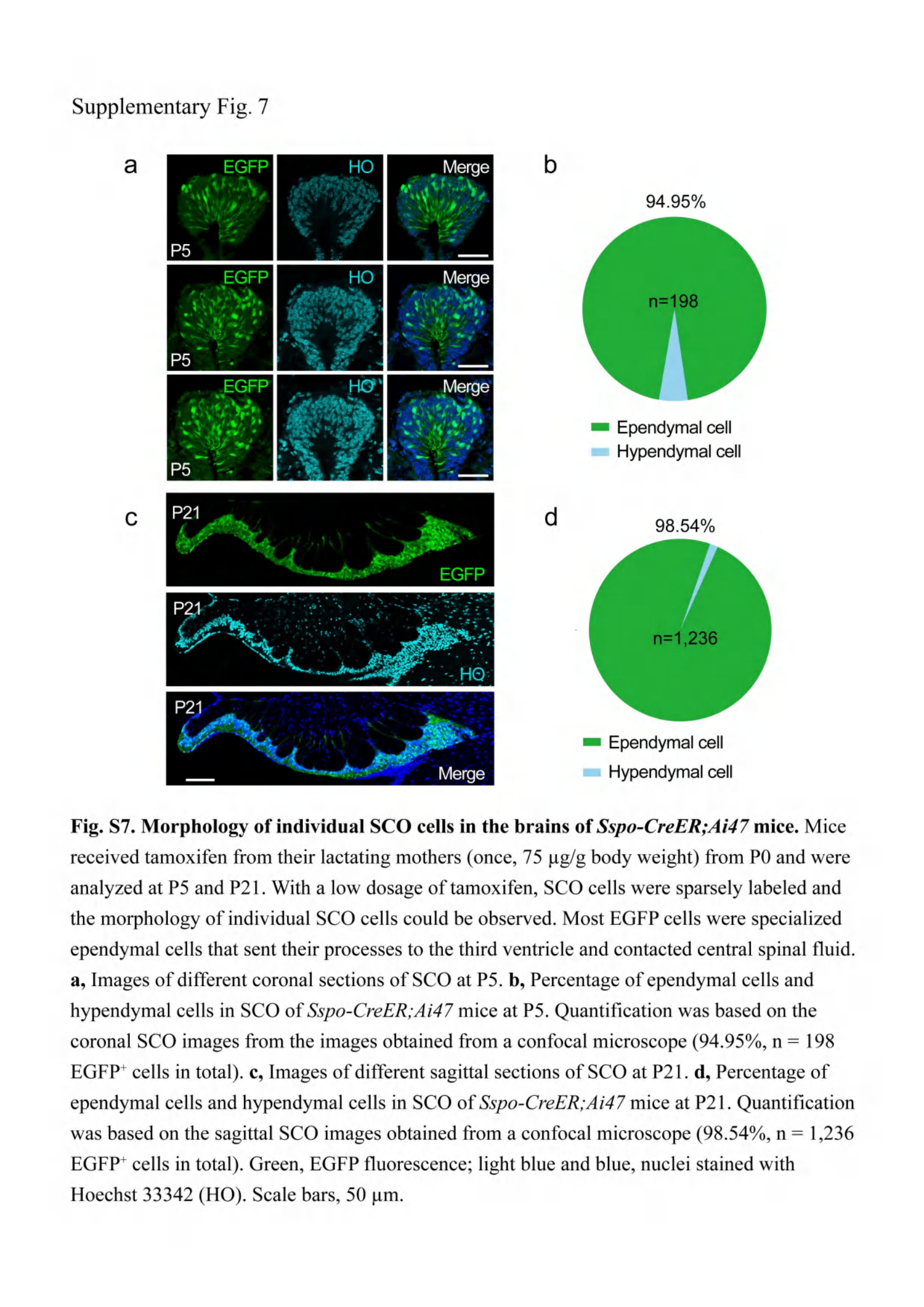

### Supplementary Fig. 8

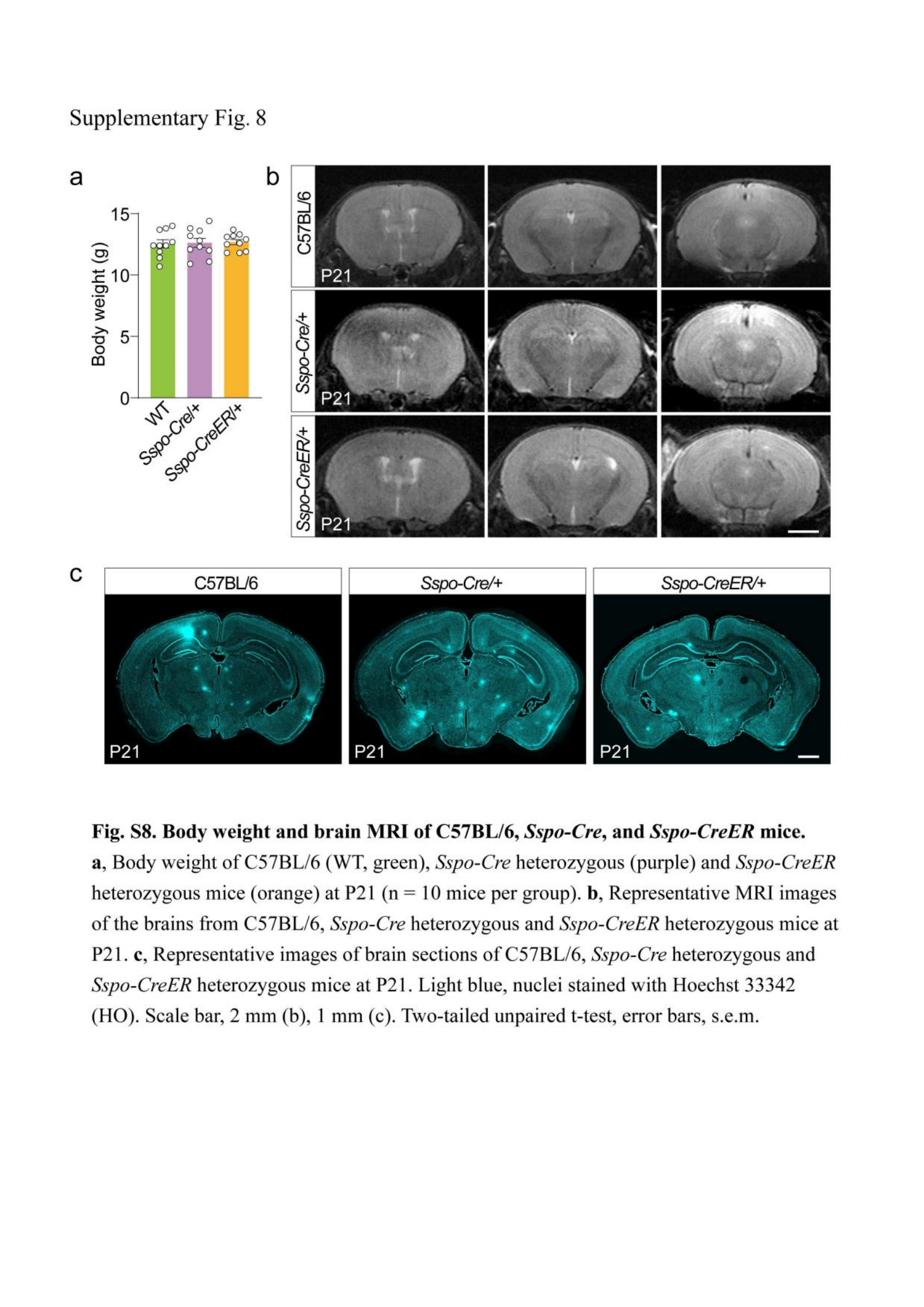

### Supplementary Fig. 9

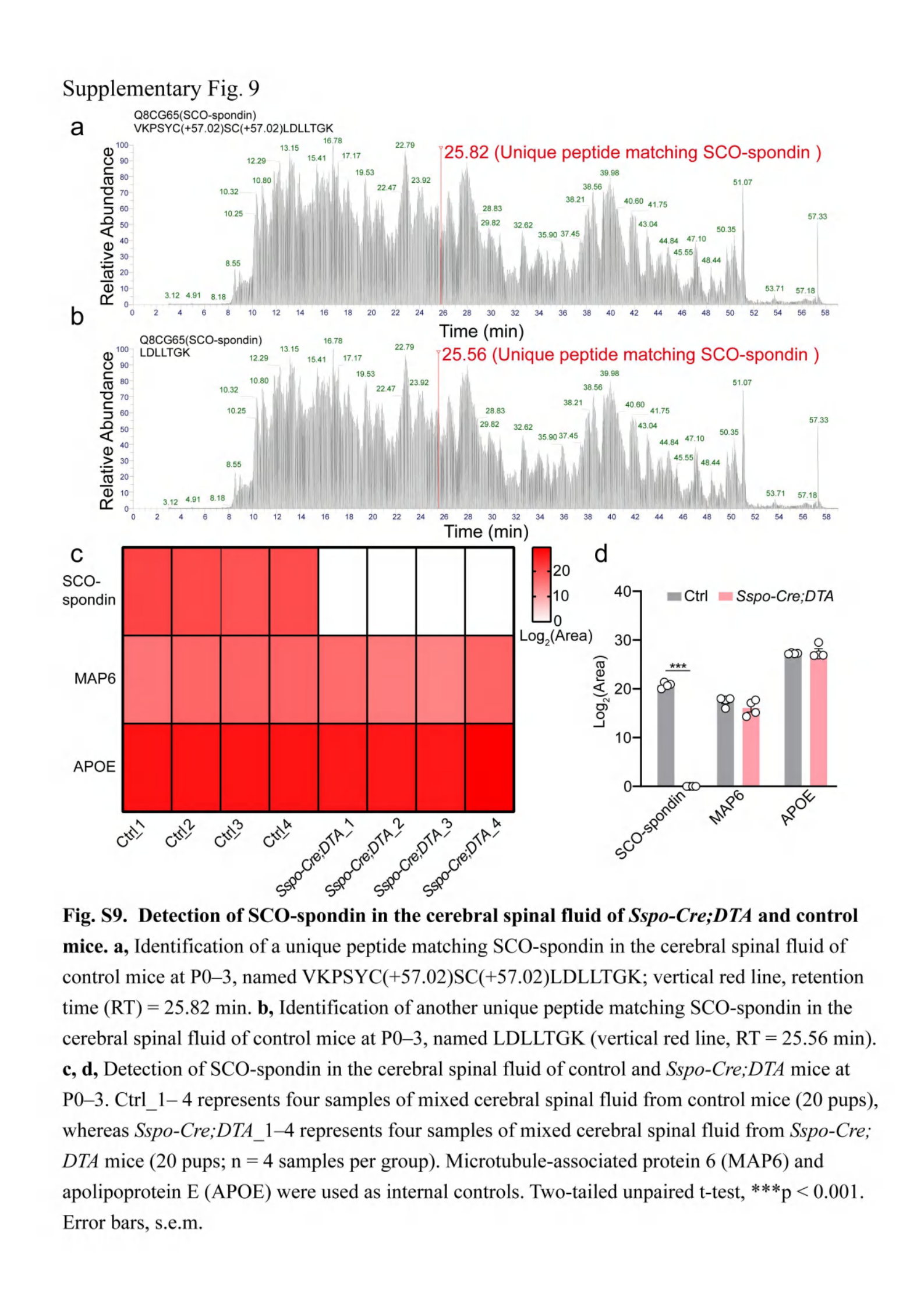

### Supplementary Fig. 10

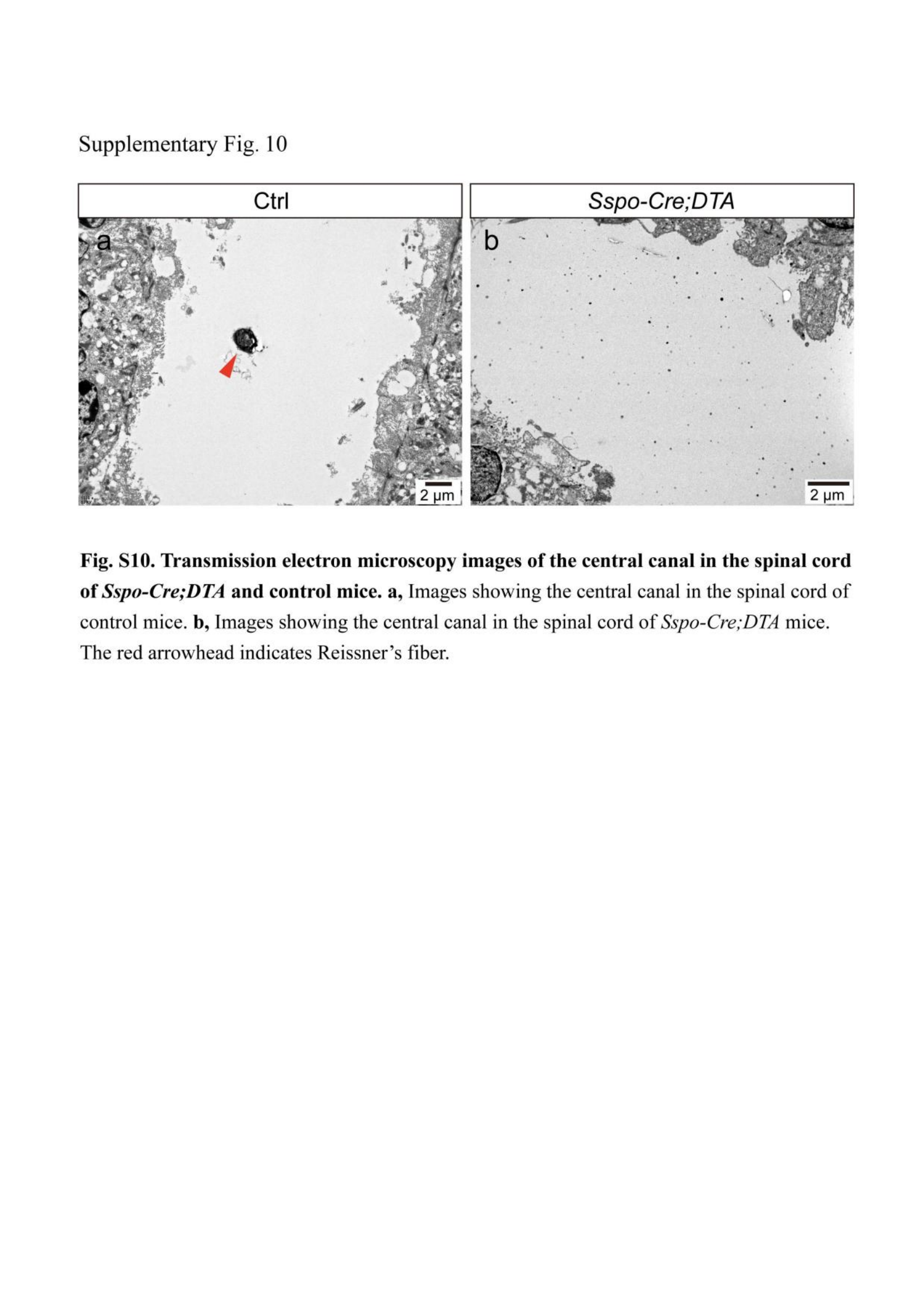

### Supplementary Fig. 11

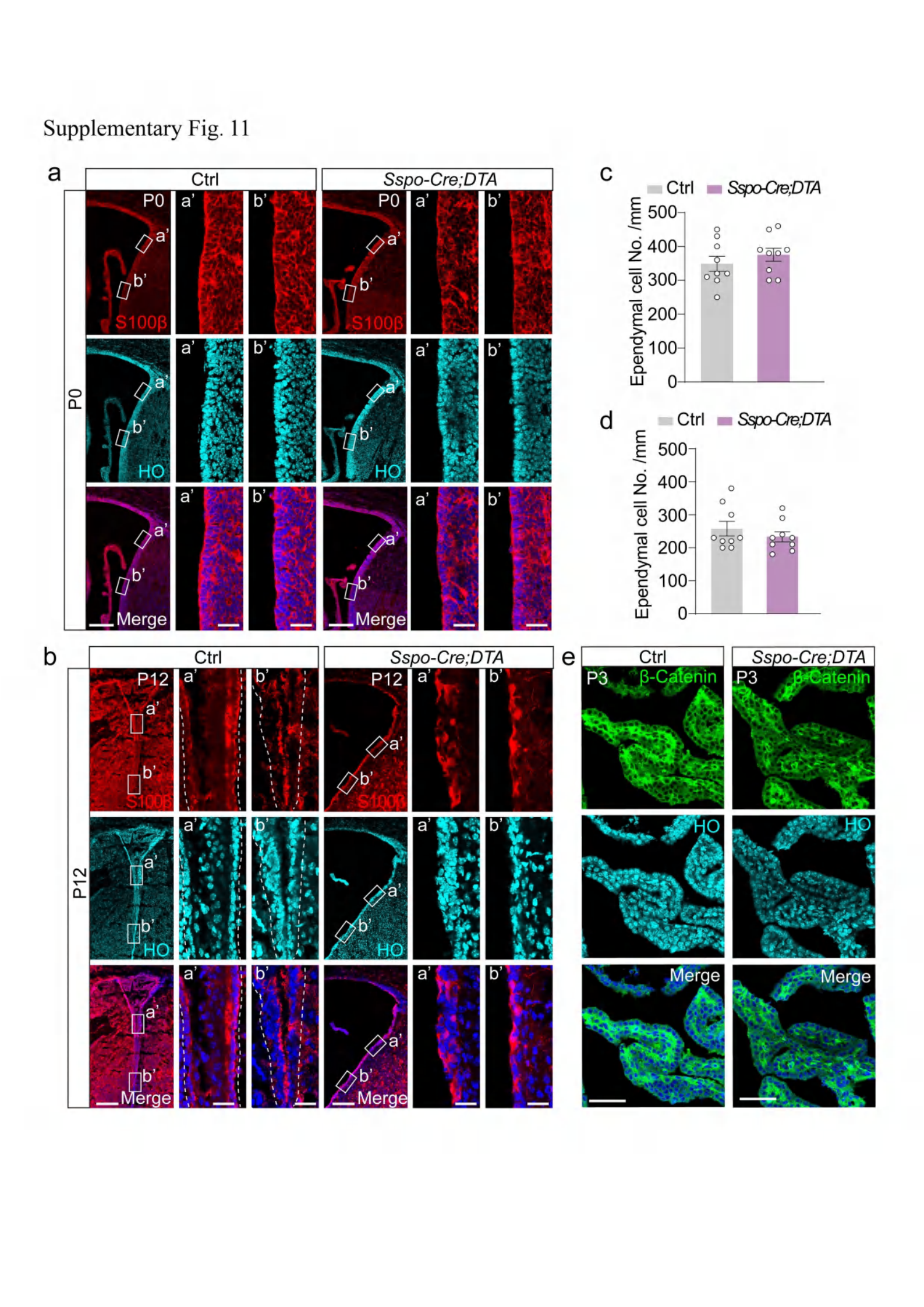

### Supplementary Fig. 11_legend

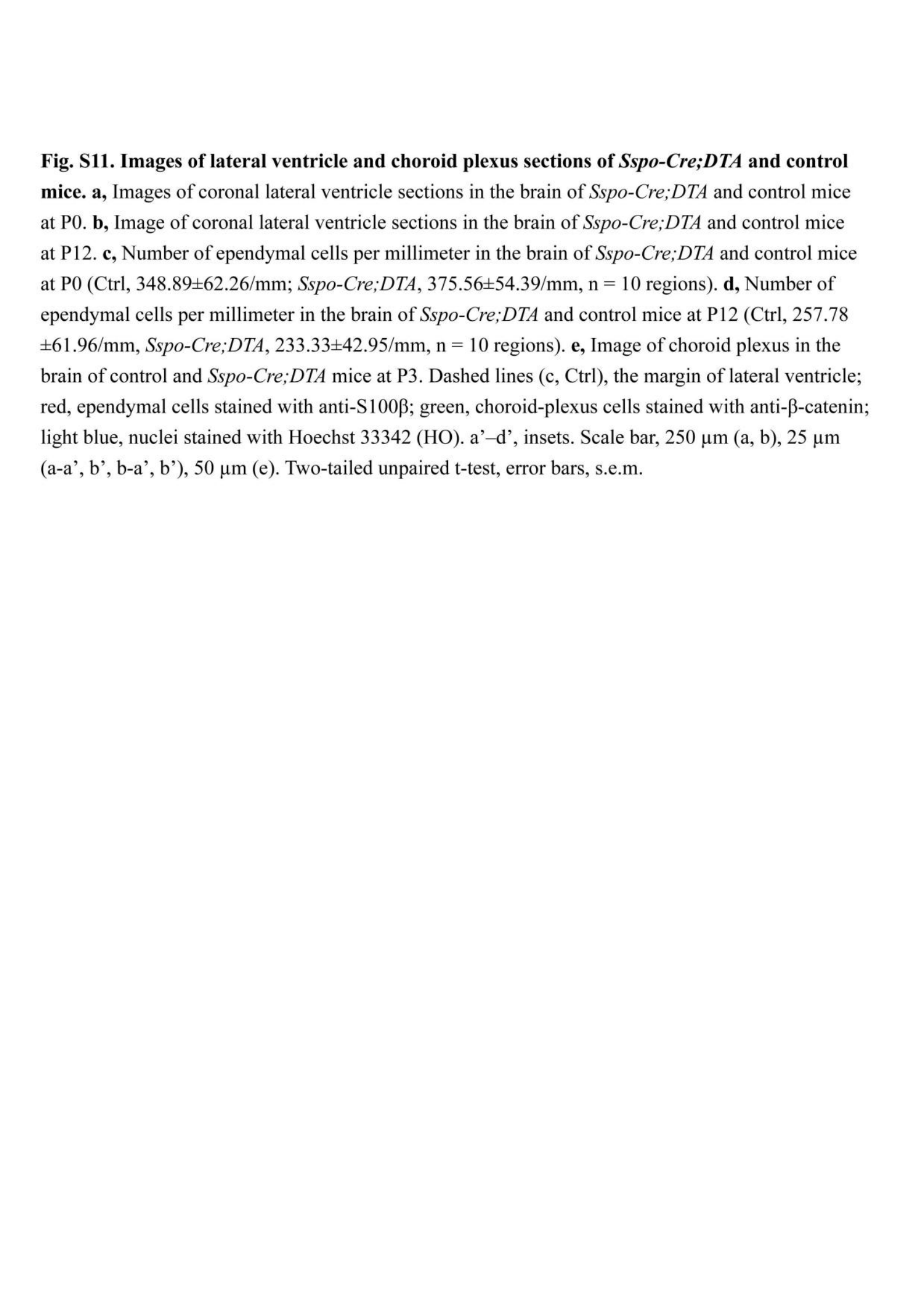

### Supplementary Fig. 12

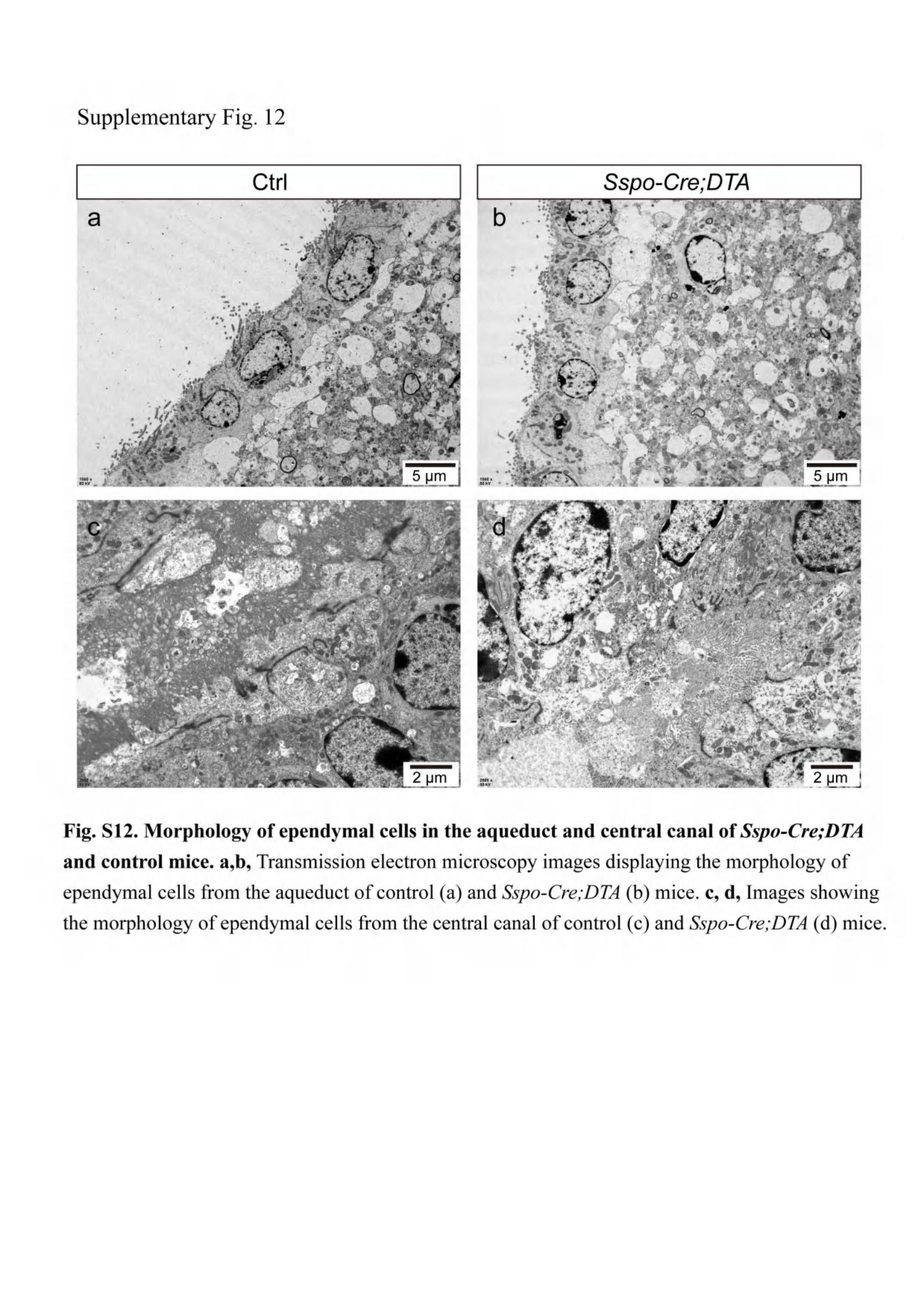

### Supplementary Fig. 13

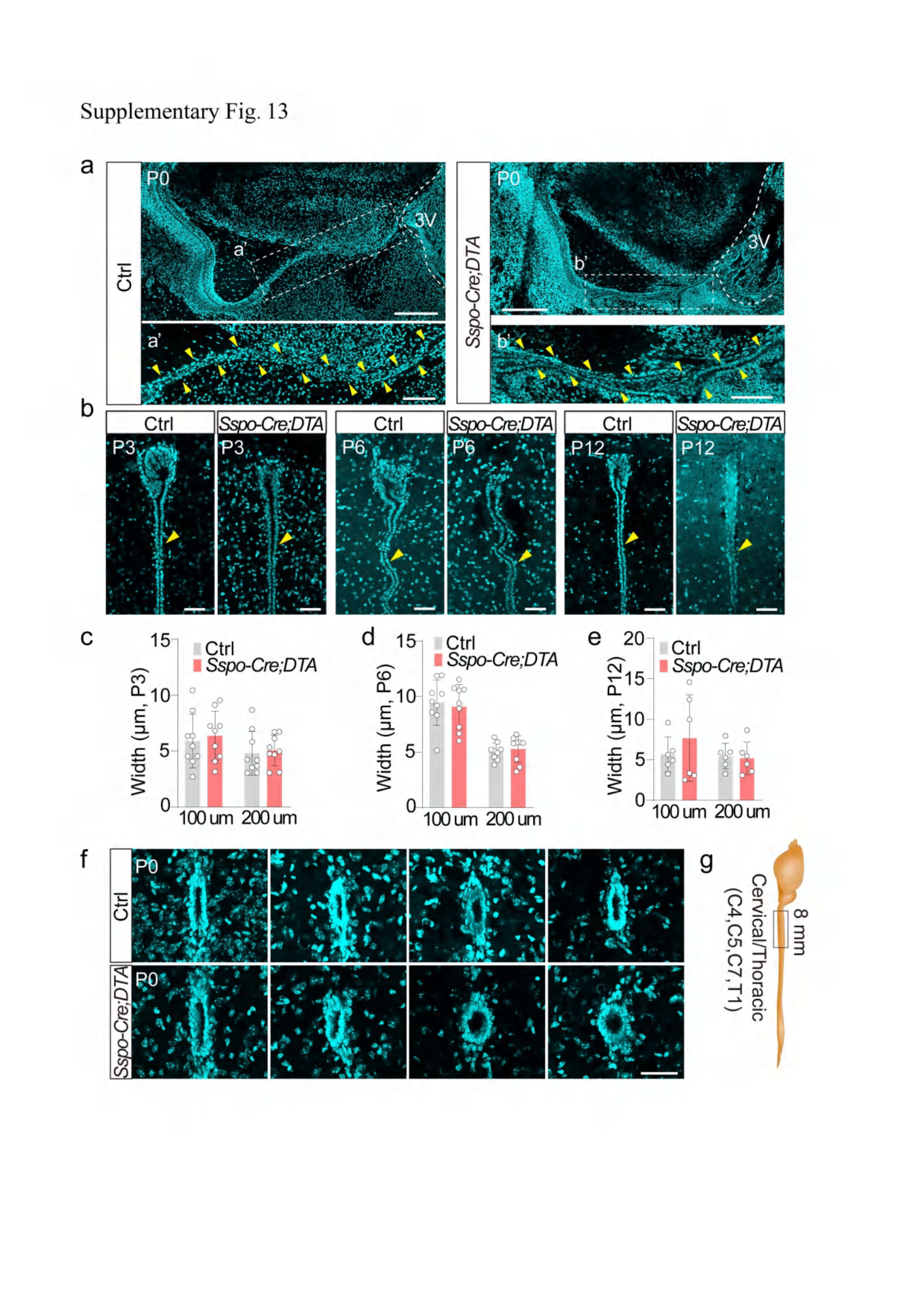

### Supplementary Fig. 13_legend

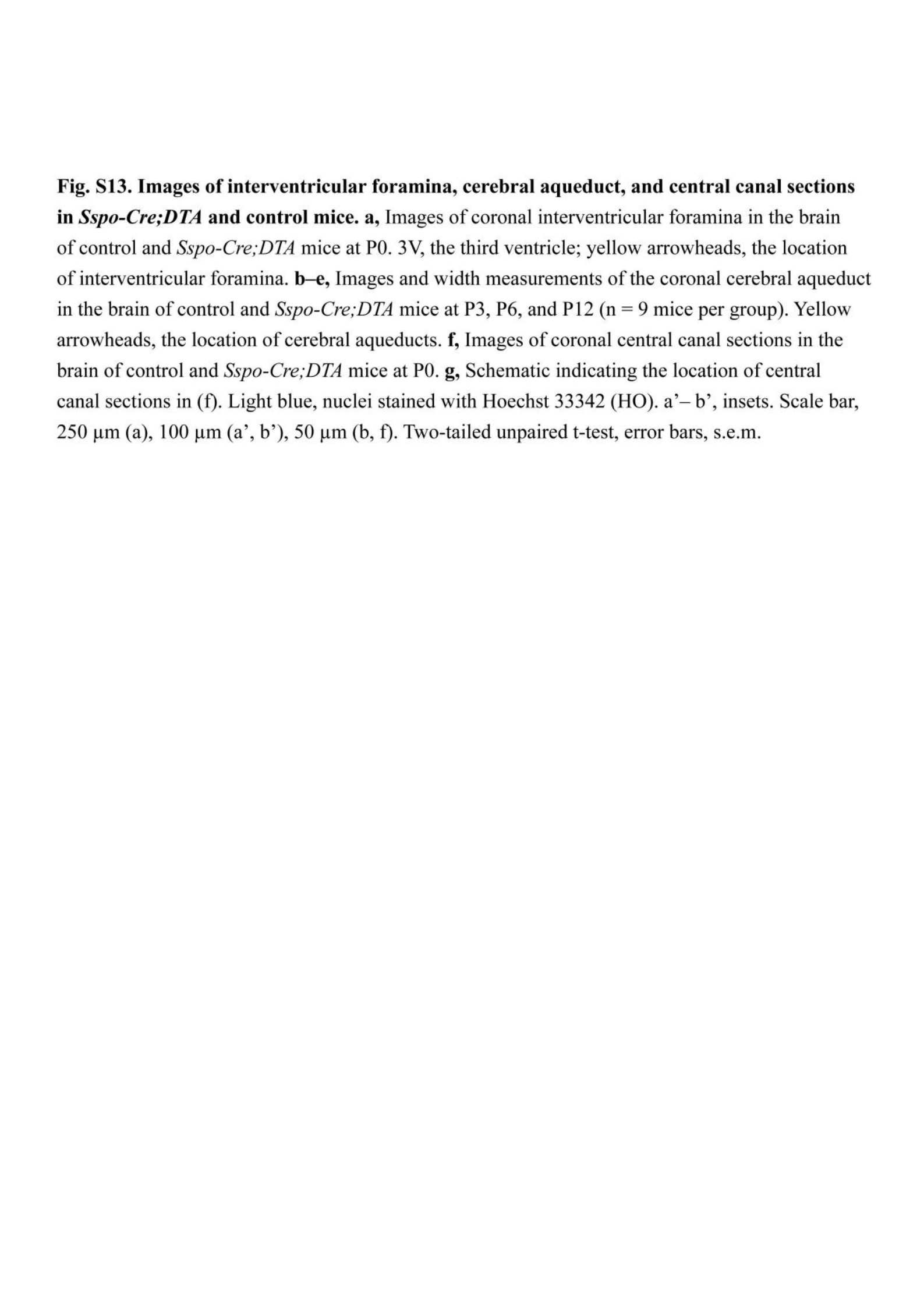

### Supplementary Fig. 14

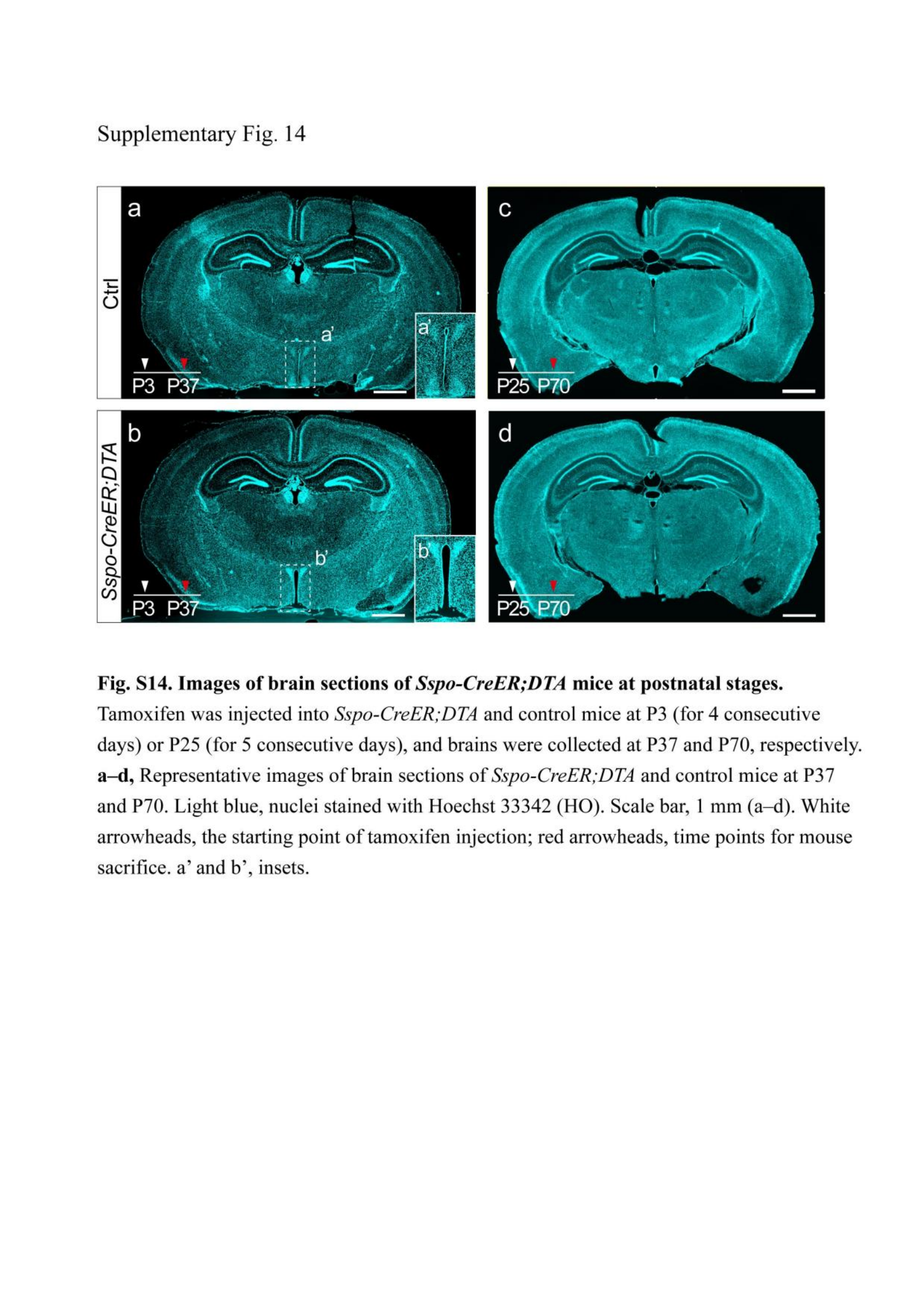

### Supplementary Fig. 15

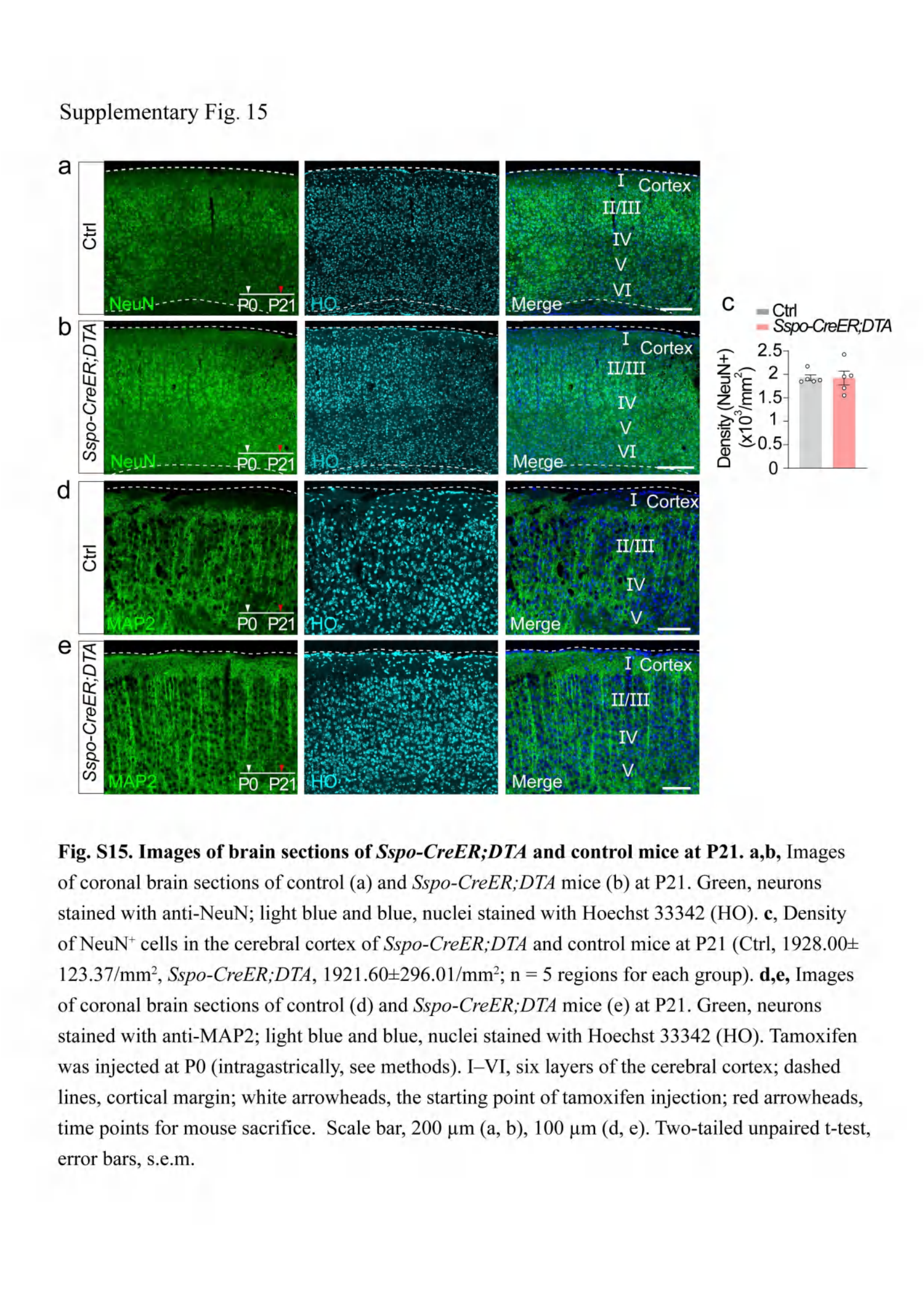

### Supplementary Fig. 16

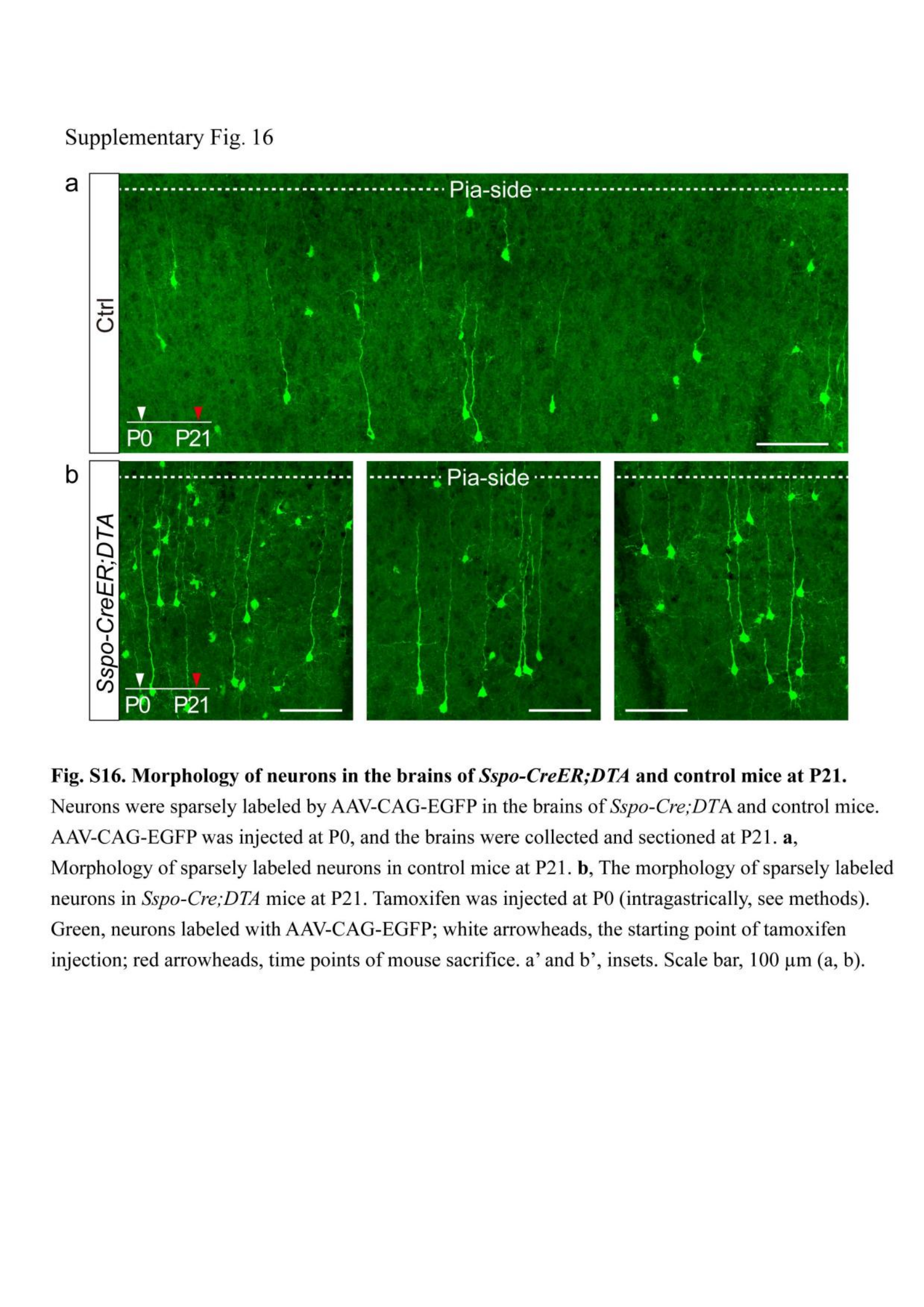

### Supplementary Fig. 17

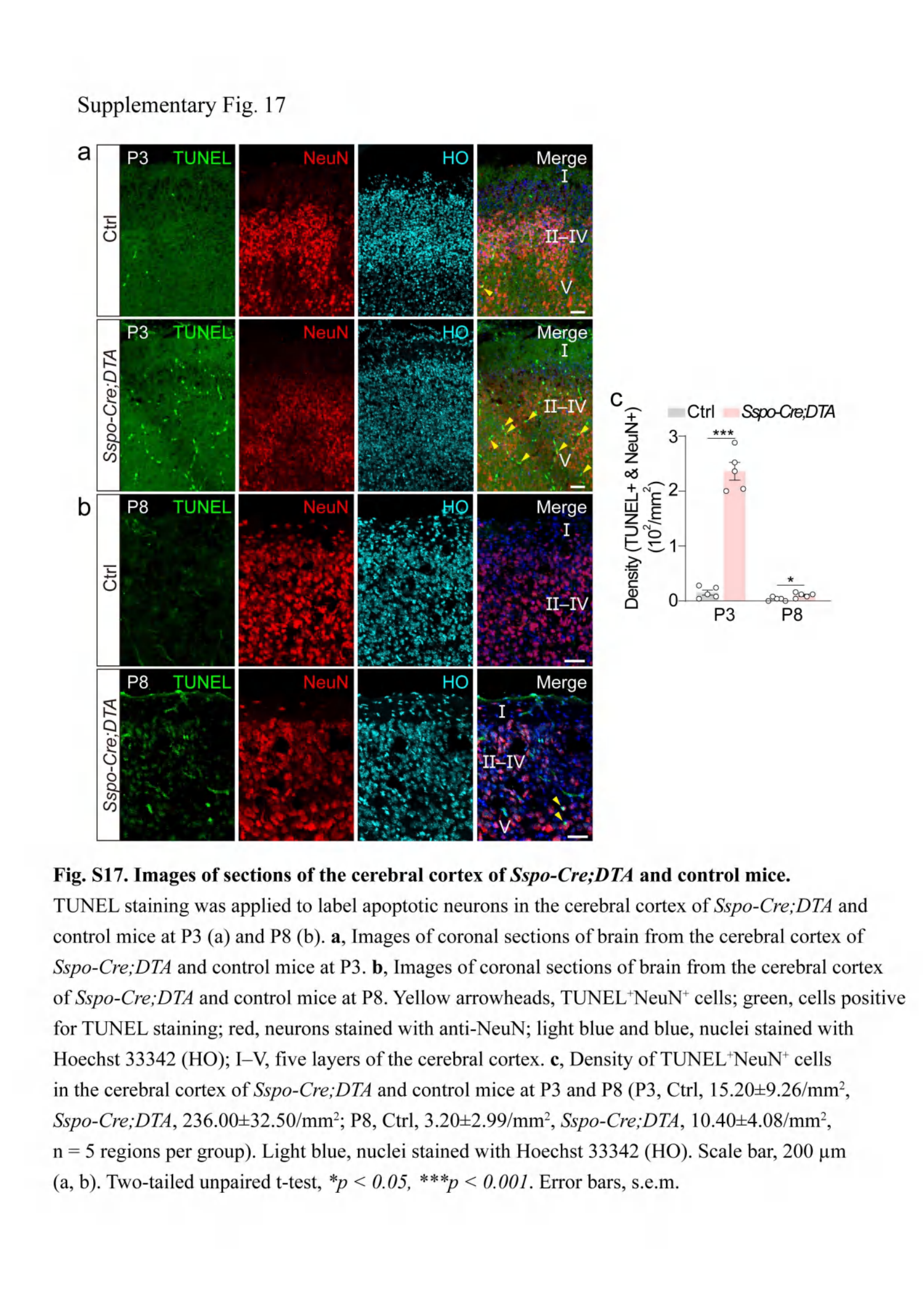

### Supplementary Fig. 18

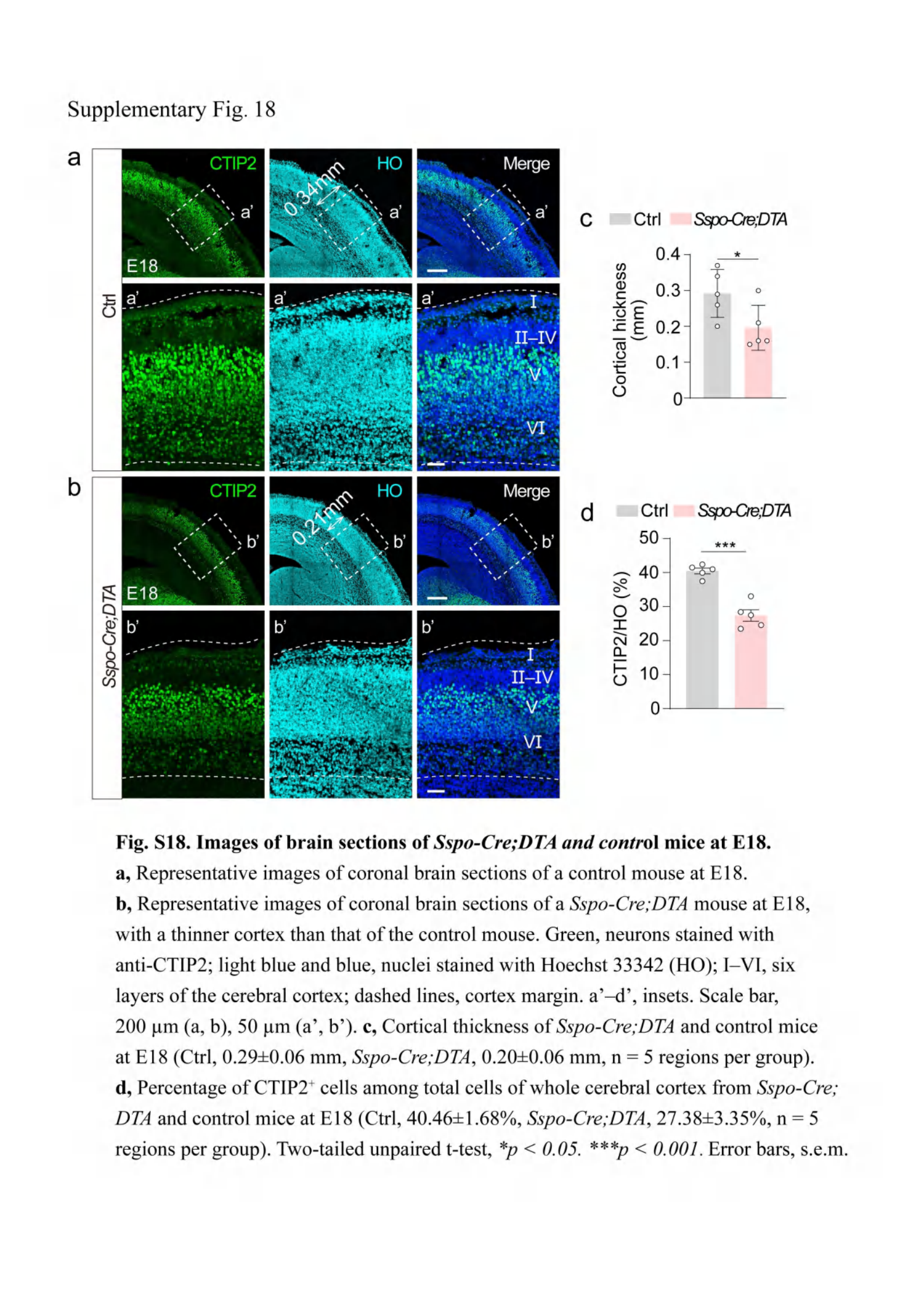

### Supplementary Fig. 19

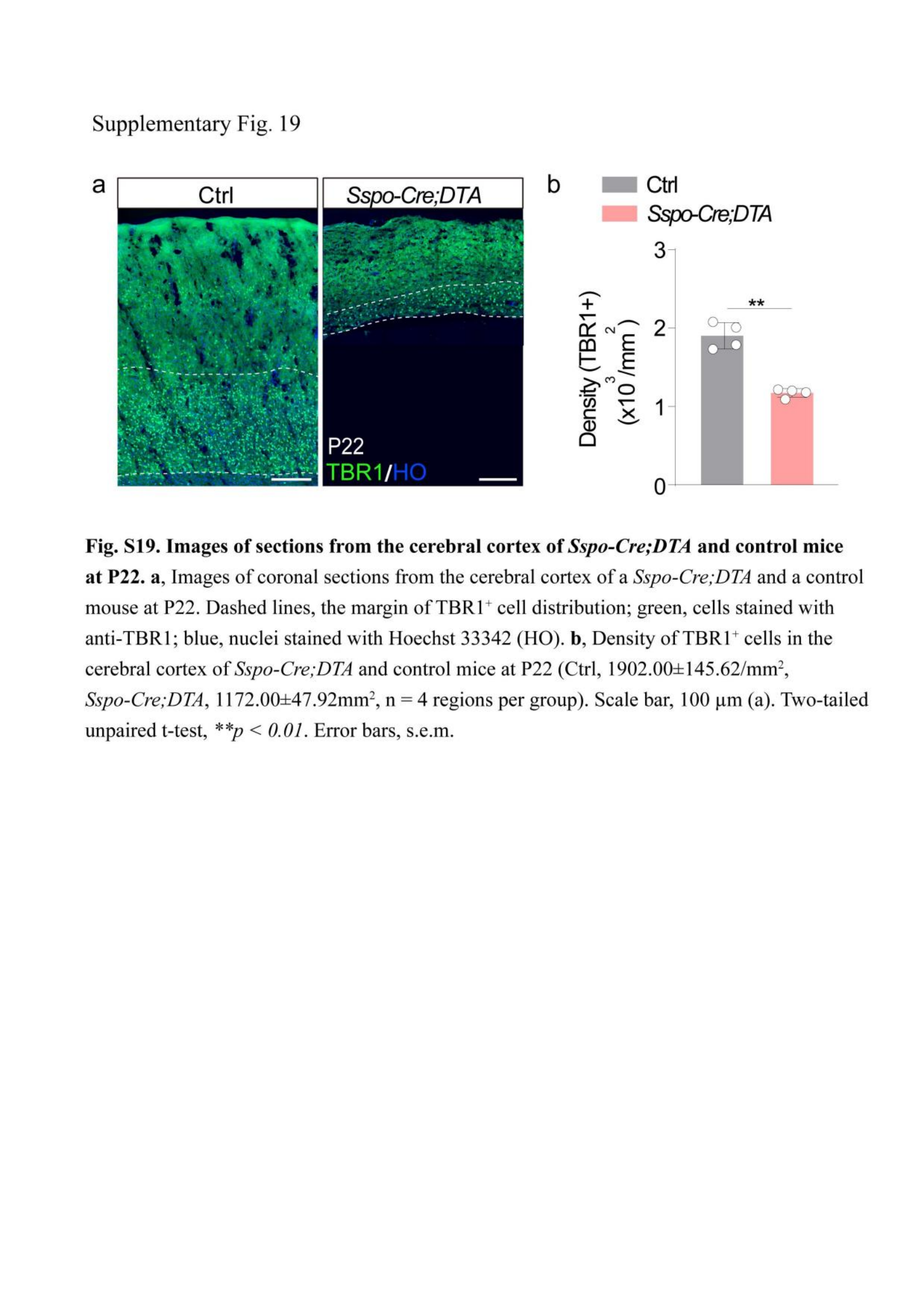

### Supplementary Fig. 20

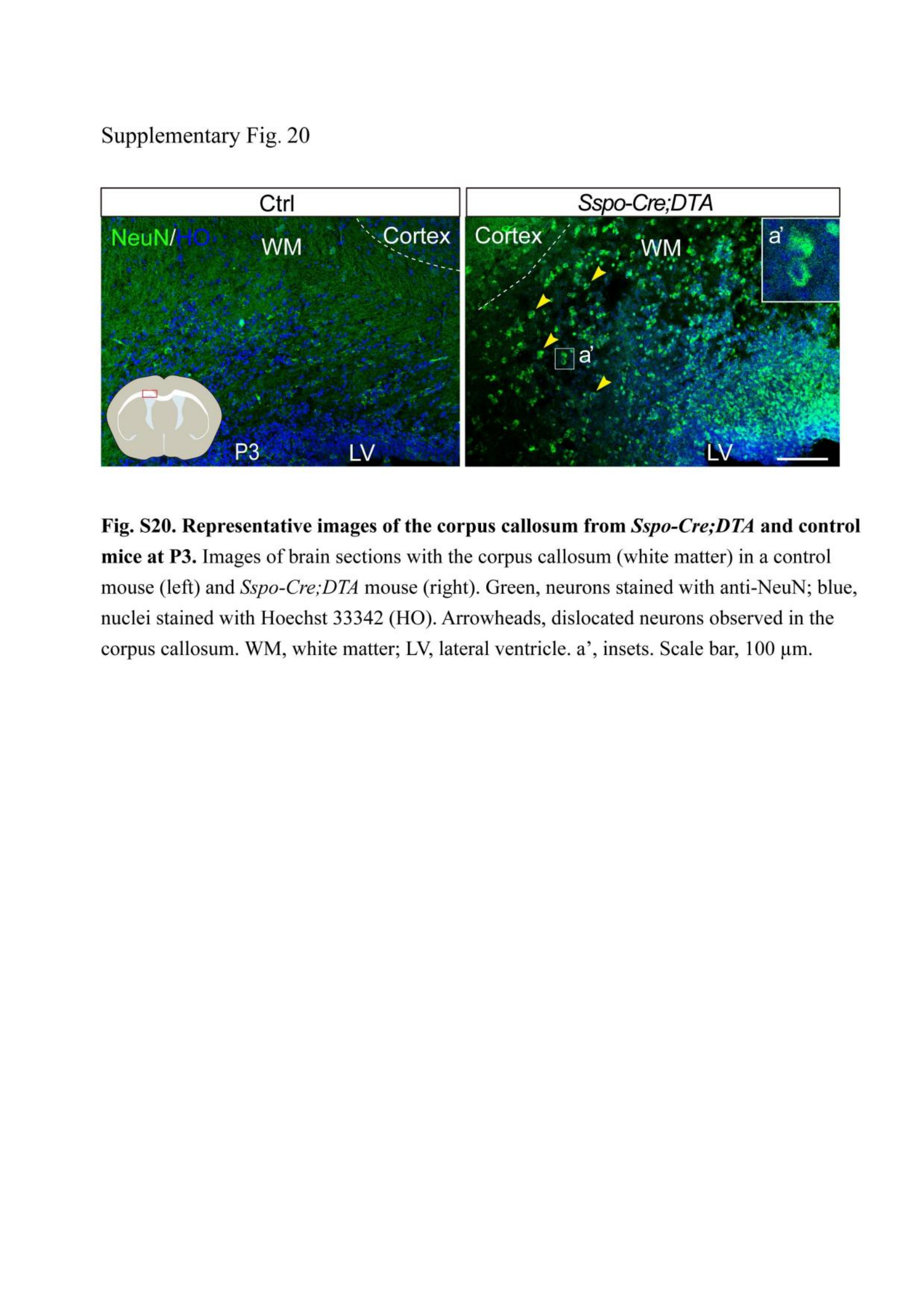

### Supplementary Fig. 21

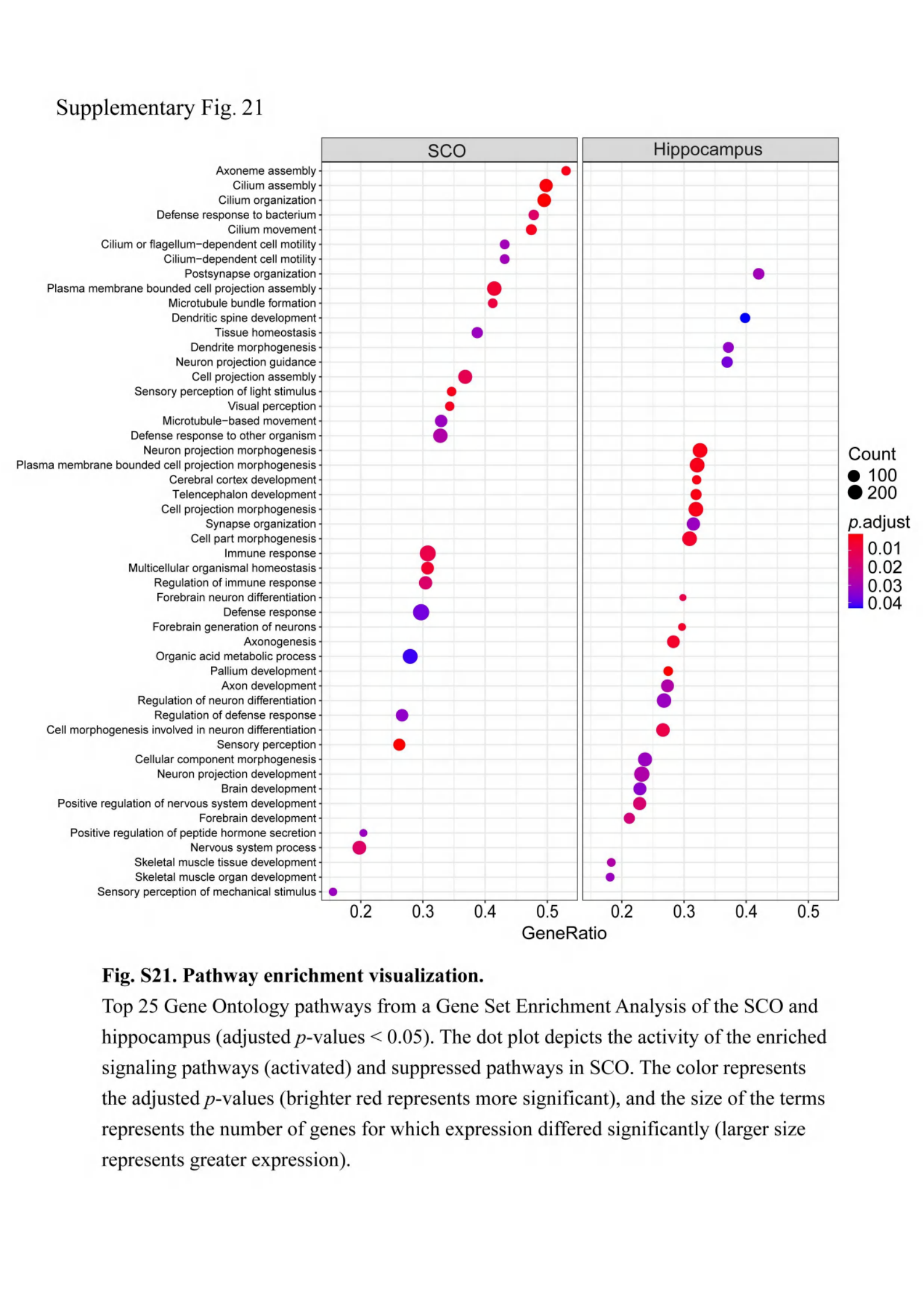

### Supplementary Fig. 22

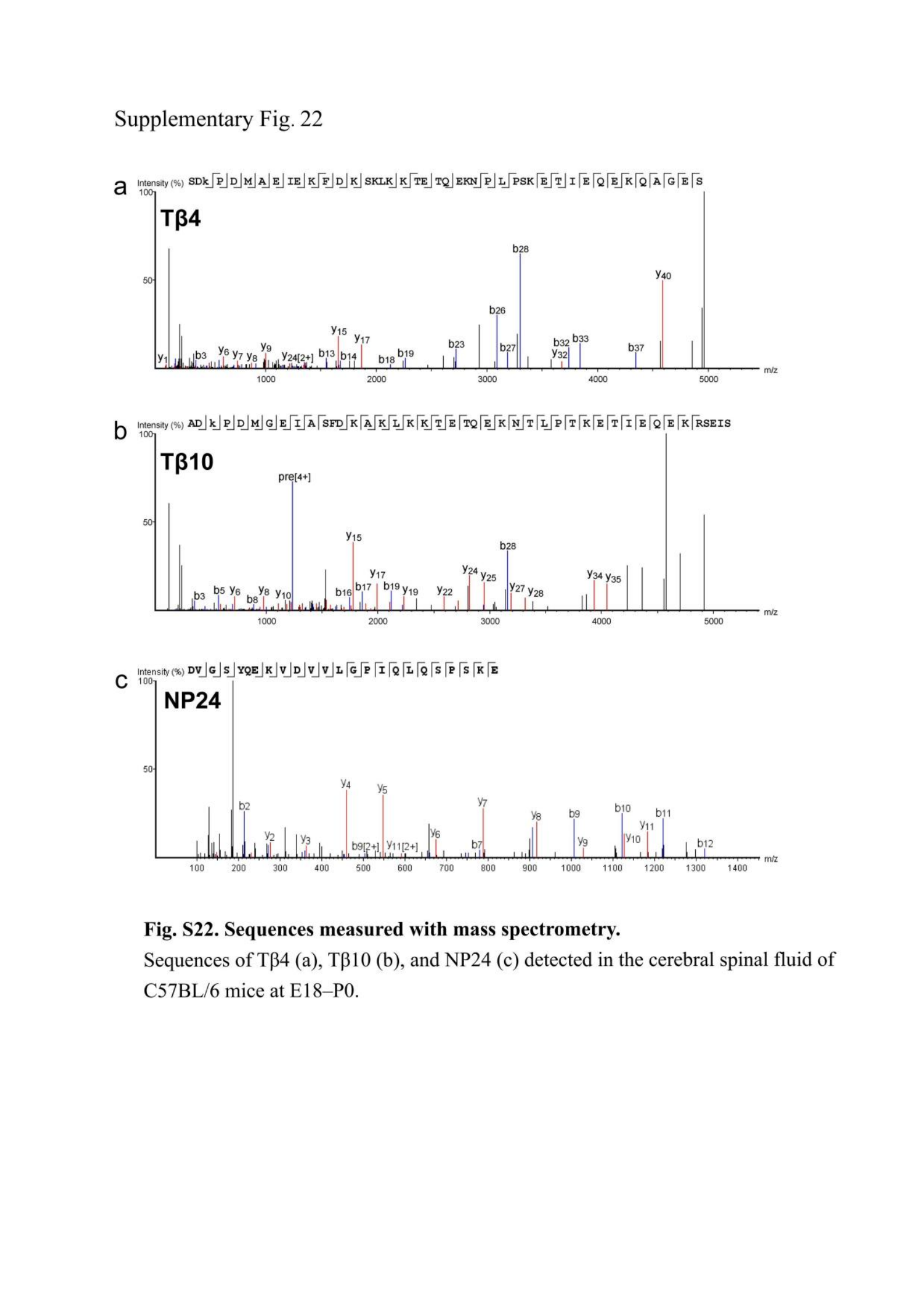

### Supplementary Fig. 23

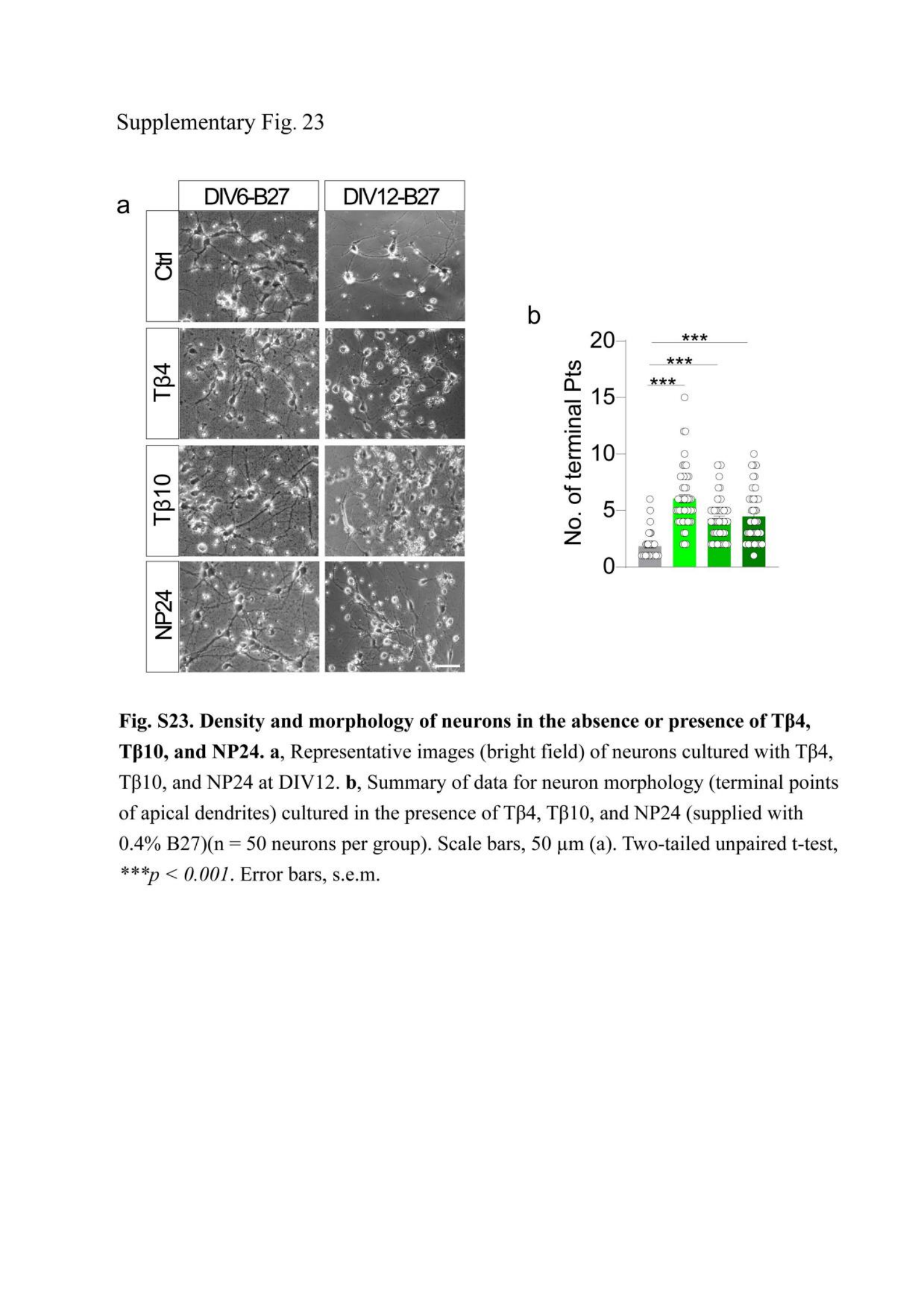

### Supplementary Fig. 24

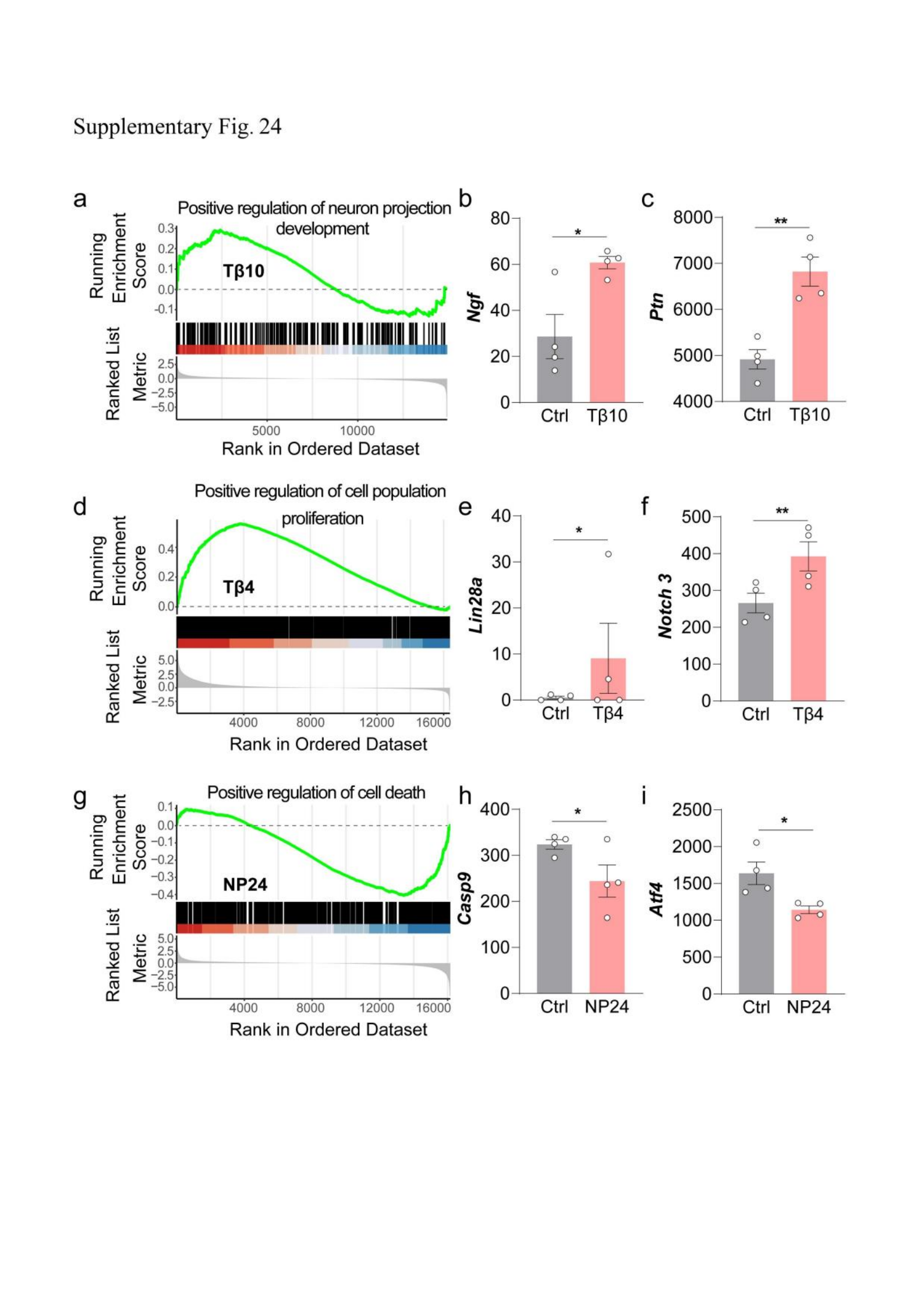

### Supplementary Fig. 24_legend

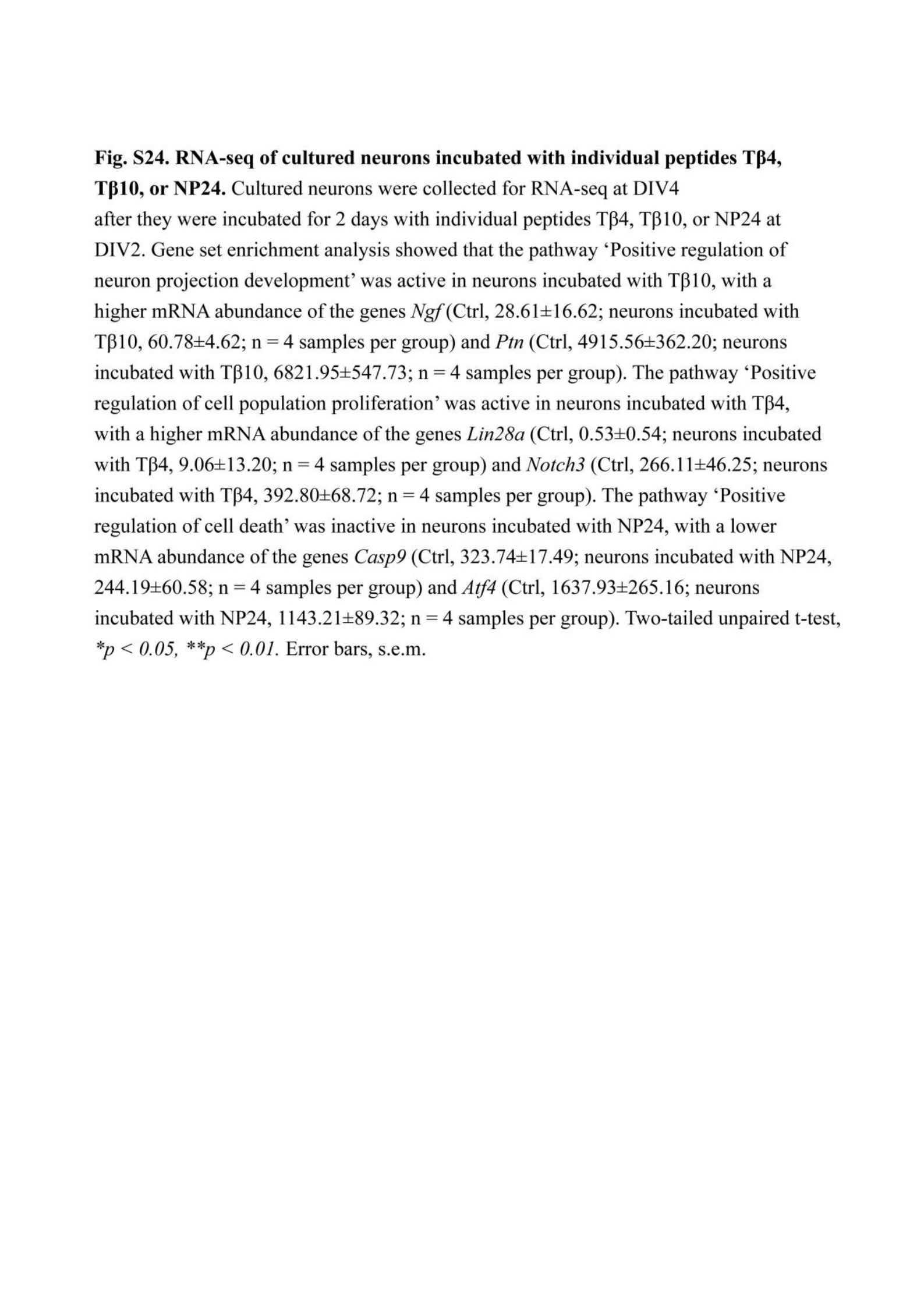

### Supplementary table 1

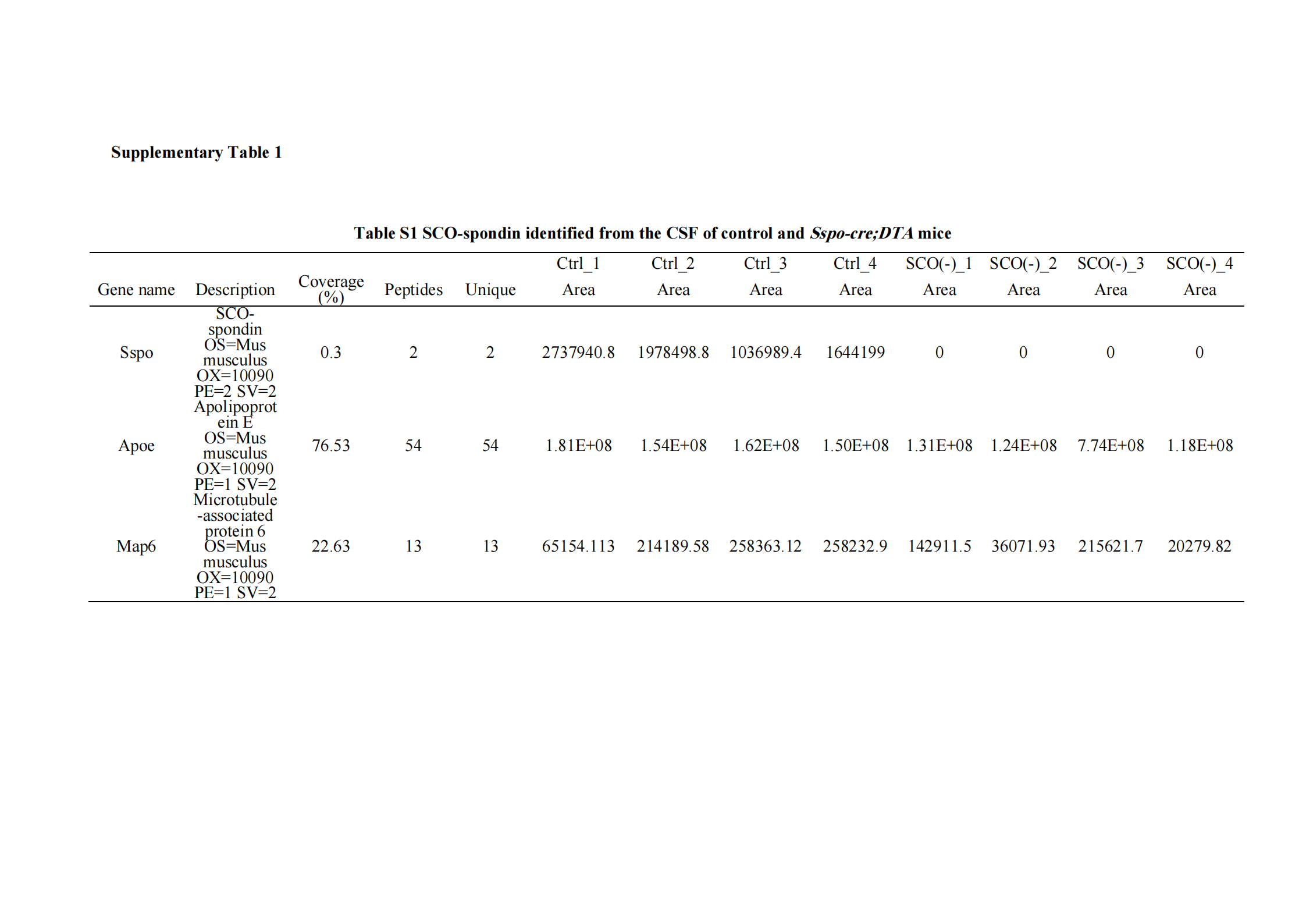

### Supplementary table 2

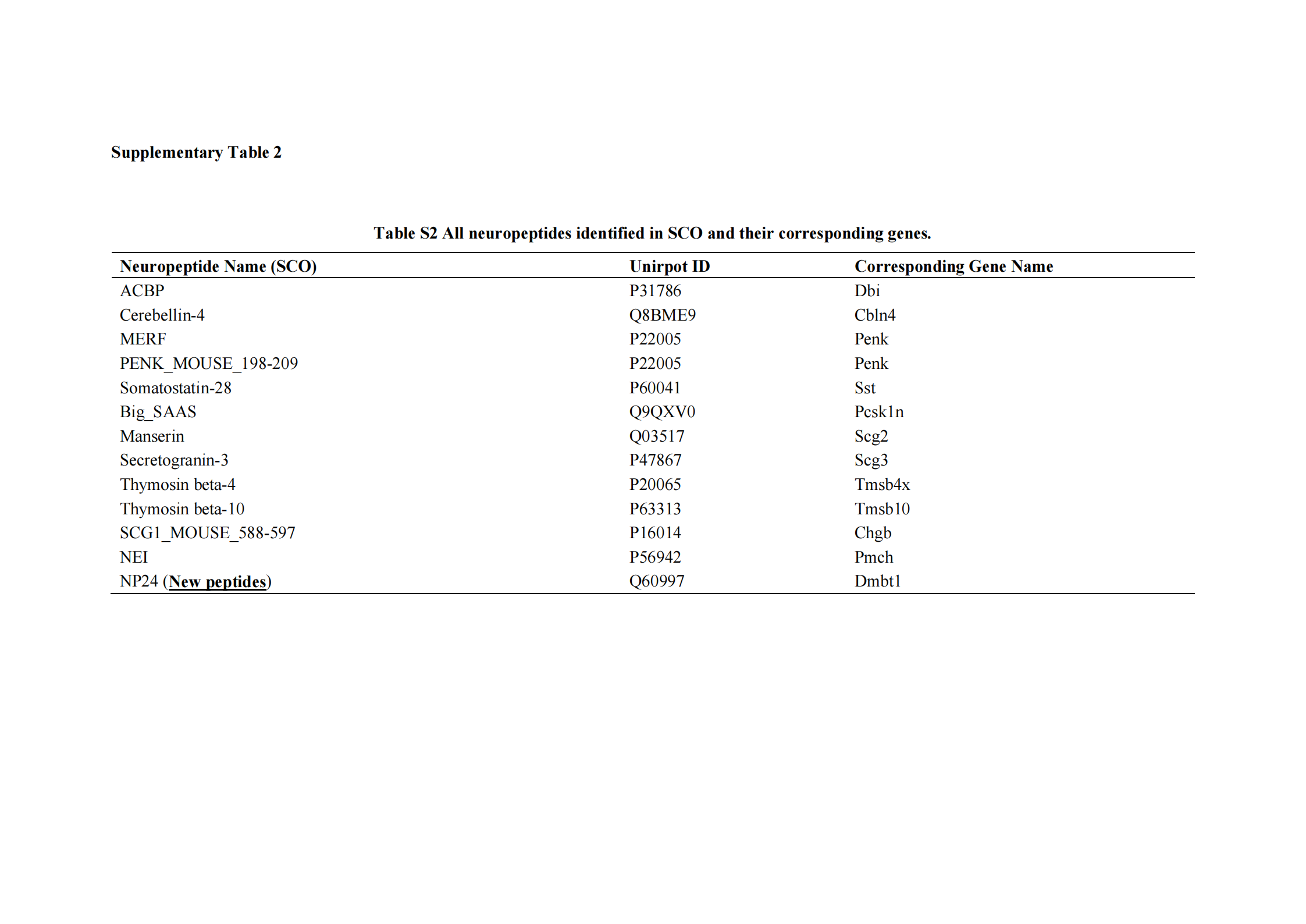

### Supplementary table 3

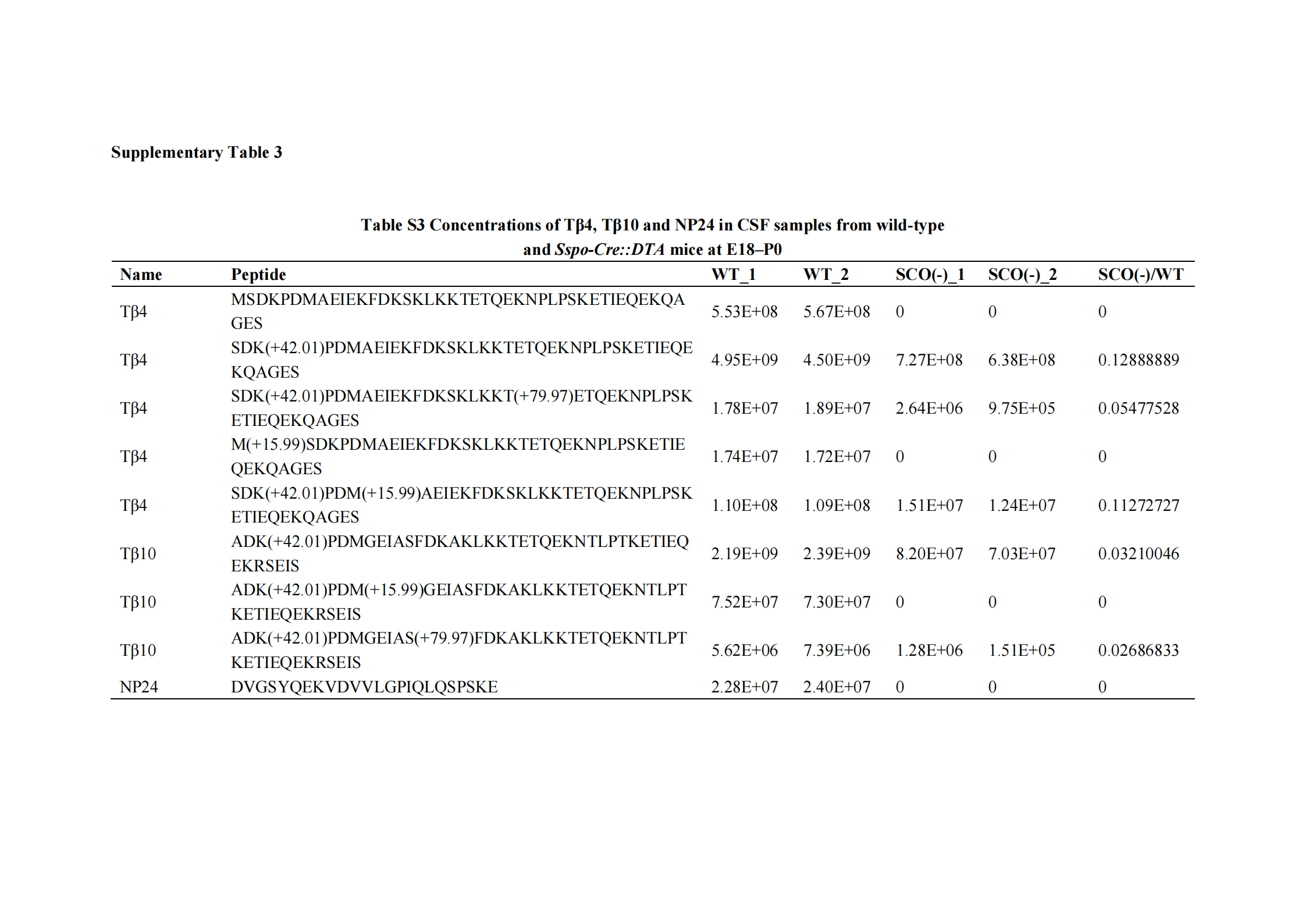
